# A long-chain heparan sulfate capture mechanism directs paracrine GDNF-GFRα1 signalling through RET

**DOI:** 10.64898/2026.05.20.726209

**Authors:** Mia I. Zol-Hanlon, Beatrice Rix, Salomé Bodet-Lefèvre, Miguel Zamora-Porras, David C. Briggs, Andrea Nans, Annabel C. Borg, Sarah L. Maslen, Antonio Di Maio, Ten Feizi, Yan Liu, Pradeep Chopra, Geert-Jan Boons, Jelizaveta Pavljuk, Ralf P. Richter, Benjamin Schumann, Neil Q. McDonald

**Affiliations:** Signalling and Structural Biology Laboratory, The Francis Crick Institute, 1 Midland Road, London, NW1 1AT, United Kingdom; Structural Biology Science Technology Platform, The Francis Crick Institute, 1 Midland Road, London, NW1 1AT, United Kingdom; Proteomics Science Technology Platform, The Francis Crick Institute, 1 Midland Road, London, NW1 1AT, United Kingdom; Glycosciences Laboratory, Department of Metabolism, Digestion and Reproduction, Faculty of Medicine, Imperial College London, London, United Kingdom; Complex Carbohydrate Research Center, University of Georgia, Athens, GA, 30602, USA; Department of Chemistry, University of Georgia, Athens, GA 30602, USA; Department of Chemical Biology and Drug Discovery, Utrecht Institute for Pharmaceutical Sciences, and Bijvoet Center for Biomolecular Research, Utrecht University, 3584 Utrecht, The Netherlands; The School of Biomedical Sciences, Faculty of Biological Sciences, School of Physics and Astronomy, Faculty of Engineering and Physical Sciences, Astbur Centre for Structural Molecular Biology, and Bragg Centre for Materials Researcher, University of Leeds, Leeds, LS2 9JT, UK; Faculty of Chemistry and Food Chemistry, TUD Dresden University of Technology, Dresden, Germany; Chemical Glycobiology Laboratory, The Francis Crick Institute, 1 Midland Road, London, NW1 1AT, United Kingdom; Department of Chemistry, Imperial College London, London, UK; Institute of Structural and Molecular Biology, School of Natural Sciences, Birkbeck College, Malet Street, London, WC1E 7HX, United Kingdom

## Abstract

The receptor tyrosine kinase RET is activated by GDNF and GFRα1 together as a bipartite ligand, driving receptor activation and signalling in developmental and neuroprotective contexts. Evidence from developmental and cell models has suggested that heparan sulfate (HS) functions as a fourth component in RET signalling by binding to GDNF, but the molecular details remain unclear. Here, we present the cryo-EM structure of the heterohexameric RET:GDNF:GFRα1 complex with a fully resolved heparin ligand, revealing an unexpected extended HS binding site spanning all three proteins. The architecture of the complex and binding mode of the HS chain in this complex enables the formation of a higher order 4:4:4 assembly bound to a single 30-saccharide HS chain which bridges two intimately bound complexes. This multi-protein interface selectively binds the highly sulfated domains of HS over other GAG classes, and is essential for RET activation *in trans* with soluble GFRα1, but not *in cis* when GFRα1 is membrane-bound. Our data suggest that HS shapes the dynamics of RET signalling at every stage, from ligand diffusion to signalling complex formation. Thus, GDNF-GFRa1 paracrine signalling reveals a surprising dependence for long-chain GAG function in which the glycan engages in both receptor complex assembly and clustering.

## Introduction

Glial cell-derived neurotrophic factor (GDNF)-driven signalling through RET is essential for the growth, guidance and survival of multiple central and peripheral neuronal populations^1–5^, as well as for kidney morphogenesis^6^ and spermatogonial differentiation^7^. GDNF, a constitutively dimeric member of the TGFβ superfamily^8^, activates RET through the dimerisation and transphosphorylation mechanism typical of receptor tyrosine kinases (RTKs), but uniquely requires the binding of GDNF family receptor α1 (GFRα1), a third protein which plays a dual role as both a co-receptor and ligand. GFRα1 is comprised of 3 α-helical domains (D1-D3) arranged spatially out-of-order with D3 positioned centrally due to an extended D1-D2 linker, and is associated with the plasma membrane via a glycophosphatidylinositol (GPI) anchor. GFRα1 binds GDNF via D2, forming a U-shaped 2:2 co-receptor-neurotrophic factor complex^9^. The 2:2 GDNF:GFRα1 complex engages the RET extracellular domain (ECD) through two spatially separated interfaces: a polar “wedge” via GFRα1 D3, and a hydrophobic “saddle” with GDNF^10–12^ **(Fig. 1A,B)**. RET is the sole RTK with a cadherin-like domain (CLD)-based ECD, with four such domains (CLD1-4) followed by a cysteine-rich domain (CRD) forming a flexible C-shaped architecture^13^. A family of RET ligands have arisen from gene duplication, providing structural variability and mechanistic diversity to support a range of biological processes: Four paralogous pairs of GDNF family ligands (GFLs) (neurturin (NRTN), artemin and persephin) and GFRαs (GFRα2-4), as well as the more divergent GDF-15 and GFRAL^14^ all engage RET. The geometry of RET complexes formed between the different GFL:GFRα pairs varies significantly through differences in the “batwing” angle made between the two halves of the 2:2:2 complex, although the significance of this finding remains unclear.

**Figure 1.**
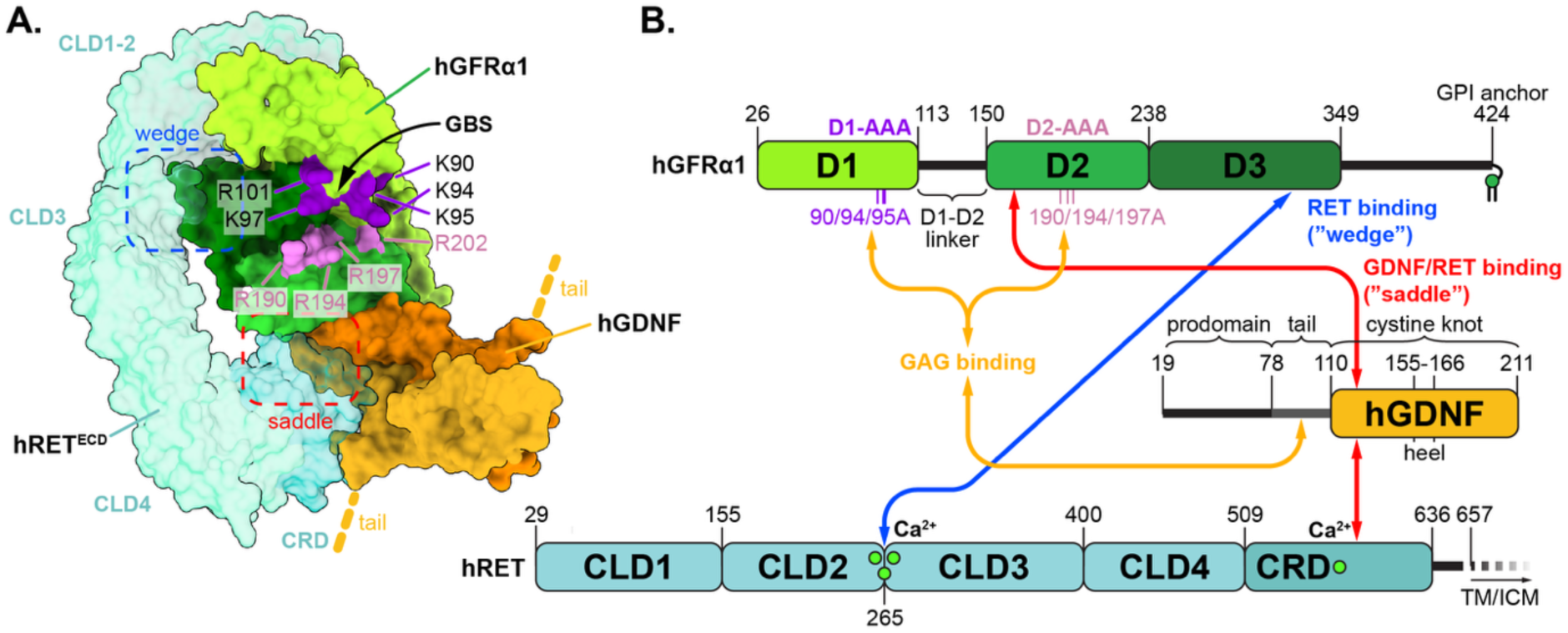
Overview of the structural basis for hRET^ECD^ receptor-ligand complex formation and GAG binding of hGFRα1 and GDNF. **(A)** Surface rendering of a composite model of a partial hRET^ECD^-hGFRα1-GDNF complex. Models for hRET^ECD^ (PDB: 6Q2N), hGFRα1 (AlphaFold 2 structure prediction, residues 26-349) and hGDNF (PDB: 2V5E) were used and are coloured according to the colour scheme in B). **(B)** Schematic domain diagrams of hRET, hGFRα1 and hGDNF, highlighting the sites important for complex formation and GAG binding.

While the requirement of heparan sulfate (HS) for RET signalling *in vivo* is known, the principles underlying its function and role in signalling are not understood. Mice deficient for HS-2-O-sulfotransferase or HS-glucuronyl C5-epimerase exhibit the same bilateral agenesis of the kidney as knockouts for RET, GDNF or GFRα1^4,15–18^. A deficiency of HS-2-O-sulfotransferase in humans results in a similar developmental disorder^19^. These findings imply that HS plays a key role in supporting RET signalling in the ureteric bud, which requires GDNF from the surrounding tissue to initiate kidney organogenesis^6^. Additionally, extracellular HS endosulfatase SULFs are required for GDNF-dependent innervation of the esophagus in mice, demonstrating that RET signalling is regulated by the sulfation structure of HS^20^. Both GDNF and GFRα1 bind HS with high affinity. GDNF binds HS via a basic region in the N-terminal region outside of the folded domain^21^. The glycan binding site (GBS) of GFRα1 is formed at the junction of D1-D2-D3 and faces inwards in the 2:2 complex with GDNF^9,22^. The GFRα1 GBS is conserved between GFRα1-3 **(Fig. S1A)**, but is not present in GFRα4, which has lost D1, or GFRAL, which has a divergent arrangement of its domains in the complex formed with RET ECD and GDF-15^10^. No structure of the RET ternary complex bound to HS has been captured, making its mechanistic roles in receptor activation unclear. While the role that HS proteoglycans (HSPGs) such as glypicans play in signalling, such as shaping the spread of HS-binding signalling molecules from their source, has been well investigated for instance in the Wnt system^23^, little information is available on how the interactions between RET ligands and HS affect their diffusion and regulates signalling.

As well as classical RTK signalling via ligand-driven dimerisation of the RET receptor, additional modes of signalling and RET-independent signalling have been established for GDNF and GFRα1, establishing these proteins as multifunctional factors which facilitate a wide range of regulatory cues. In addition to signalling *in cis* with RET resident on the same cell, GFRα1 can be shed from the cell surface by phospholipases to function *in trans* in a non-cell autonomous signalling mode, enabling the activation of RET by GDNF on cells which do not express GFRα co-receptors^5,24–26^. RET activation *in trans* can cause differential signalling outputs^27^, as GFRα1 induces translocalisation of the signalling complex into lipid raft microdomains via its GPI anchor^28,29^. RET-independent receptors for GDNF and GFRα1 include neural cell adhesion molecule (NCAM), which can modulate its adhesion function^30,31^ and act as a guidance cue for olfactory bulb precursors in response to GDNF-GFRα1^32^. Membrane-anchored GFRα1 also functions as a ligand-induced cell adhesion molecule with GDNF to induce synaptic development in hippocampal neurons^33^, likely mediated by the formation of a decameric “barrel” complex in which pentameric “caps” formed by the GFRα1 D2-D3 module on two membranes are bridged by five GDNF dimers^22^.

HS, a major constituent of the extracelluar matrix (ECM), has been implicated in a wide range of signalling processes, but the molecular mechanisms and structural details of HS function remain poorly defined in most systems. Like many glycans, HS exhibits a huge degree of heterogeneity due to the stochastic nature of its biosynthesis. The sequence of HS is characterised by regions of near-maximal modification (N-sulfated (NS) or highly sulfated domains) interspersed by longer, less modified N-acetylated (NA) domains, a property not shared by other glycosaminoglycans (GAGs) which exhibit a more even pattern of sulfation^34^.

However, protein-HS interactions outside of a few specific examples do not appear to be dependent on the recognition of specific subsequences, but instead occur through general recognition of the dense negative charge of the highly sulfated domains. *In vivo*, ablation of enzymes responsible for individual modifications causes compensatory changes in the sequence of the HS that is produced, for instance by changes in the NS/NA domain structure to maintain the overall charge density when a sulfotransferase is knocked out^35,36^, making clear structure-functional relationships for individual HS modifications difficult to establish. HS-protein binding motifs have arisen independently through convergent evolution^34^, and functionally relevant interactions appear to generally be charge-mediated interactions to the NS domains. Hence, a variety of HS-protein binding modes are found in the interactions that have been structurally characterised, for instance in FGF2-FGFR1, in which the receptor-ligand complex is dimerised by binding to a single chain^37^, the activation of antithrombin by a large conformational change induced by heparin binding^38^, and the HS binding-dependent clustering of semaphorin-5A^39^. HS-binding proteins may also bind other GAGs, such as the structurally related chondroitin sulfate (CS). This mode of versatile GAG binding has been found to modulate opposing functional outputs of RPTPσ via clustering induced by HS but not CS^40^, so HS-binding protein function must therefore be considered alongside other GAGs.

In this report, we use a range of biochemical, biophysical and structural methods to describe an extended glycan binding interface in the RET:GDNF:GFRα1 ternary complex formed across all three protein constituents and is highly selective for heparin and the NS domains of HS. Furthermore, we find that HS is necessary for RET activation *in trans* in a neuronal signalling assay. We also report that a higher order tetrameric RET ternary complex can form on a single HS chain. These results explain the basis for GAG-driven facilitation of RET activation, and illustrate that sulfated glycans in the ECM shape the dynamics of RET and its ligands from their secretion to reception.

## Results

### Distributed elements across GDNF and multiple domains of GFRα1 contribute to a HS-binding site

To characterise the glycan binding capacity of GFRα1 in terms of its ligand specificity and sequence-level determinants forming the binding site, we used two glycan microarray platforms to screen recombinant hGFRα1-CHis (residues 26-424, spanning the entire mature sequence excluding the GPI anchor)**(Fig. S1B)** against a library of GAG oligosaccharide probes printed on a solid support. To assess the binding of GFRα1 to different classes of GAG, we used a neoglycolipid (NGL)-based GAG microarray, in which the glycan probes are lipid- conjugated and printed in a liposomal formulation^41^. The GAG NGL library comprised 77 probes, spanning all major classes of GAGs, including CS (-A and -C), dermatan sulfate (DS), HS and heparin, as well as keratan sulfate (KS) and the non-sulfated GAG hyaluronic acid (HA), with a chain length range of 2-20 saccharide units (degree of polymerisation, DP2-20)**(Table S3)**. Screening hGFRα1-CHis against this array indicated the co-receptor has a selective binding capacity for HS and heparin probes, which is dependent on the putative glycan-binding site (GBS) formed by domains D1 and D2 **(Fig. 2A)**. The most strongly bound probes were those in the heparin and HS classes, with a lower binding to DS and longer CSC probes. Comparatively very low binding was detected to CSA and KS probes, and no binding was detected for hGFRα1-CHis to any of the HA probes. A structure-guided triple alanine mutant construct of GFRα1 (hGFRα1-CHis D2-AAA) was prepared, targeting basic residues on the D2 domain identified from a structure in complex with the sulfated glycan mimic sucrose octasulfate^9^ (SOS)(PDB: 2V5E) **(Fig. S2A)**. This mutant showed greatly reduced binding across all sulfated GAG probes when screened on the same array, with detectable fluorescence intensity observed only for HS and heparin probes. Length-normalised class-wide mean binding of hGFRα1-CHis D2-AAA was reduced to 5% and 20% respectively compared to WT protein, demonstrating an overlap between the mutated residues and the GFRα1-GAG interaction.

**Figure 2.**
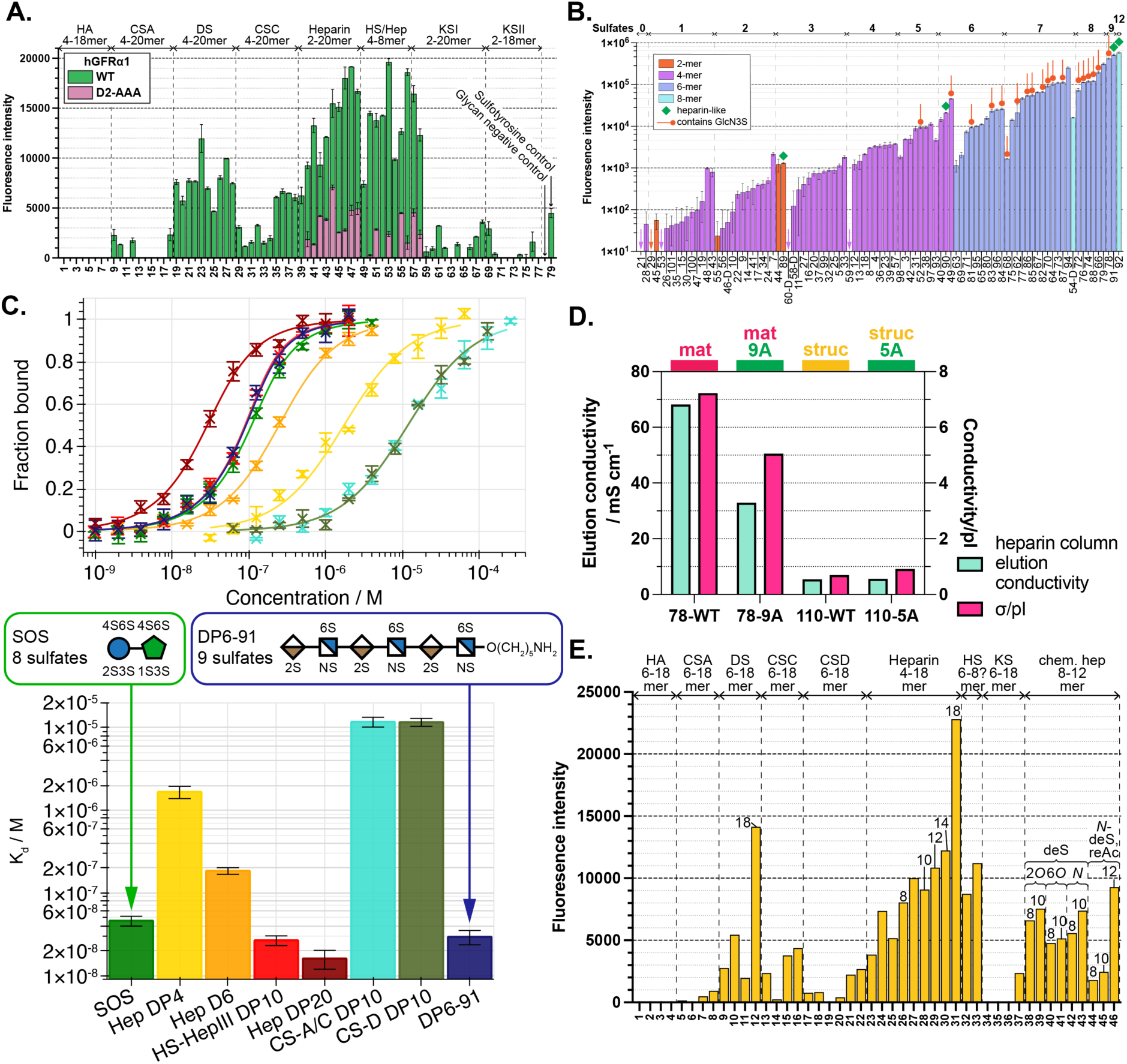
The RET signalling system components GDNF and GFRα1 share a specific, high-affinity and charge-based interaction with the sulfated glycosaminoglycan HS. **(A)** Binding analysis of hGFRα1-CHis (WT, D2-AAA) using a GAG NGL microarray. The fluorescence intensities shown are the mean values of duplicate spots (error bars represent half the difference between independent readings of the same probe). **(B)** Binding analysis of hGFRα1-CHis to synthetic HS oligosaccharides by microarray. **(C)** Binding curves and calculated *K*_d_ values for hGFRα1-CHis to natural-derived GAG oligosaccharides measured by MST. Binding data points are plotted ±SD as the mean for each concentration, normalised to the fraction bound of the binding model. Derived *K*_d_ values for each ligand are plotted with ±the standard error of the least-squares fit of the 1:1 binding model to the data. **(D)** Binding strength of hGDNF proteins to a heparin-functionalised resin as expressed by the ionic strength required to elute the protein, in conductivity and conductivity divided by the calculated theoretical isoelectric point. **(E)** Binding analysis of hGDNF^mat^ using a GAG NGL microarray.

Having established a binding selectivity of GFRα1 for HS-type GAGs, we next sought to determine the specific ligand requirements of this interaction at the level of HS sequence. We used a second glycan microarray platform, containing a library of 95 synthetic HS probes with lengths between DP2 and DP8^42^. Binding experiments on this synthetic array corroborated that hGFRα1-CHis required a WT GBS for binding to HS **(Fig. S3)**. A length- and sulfation-dependent selectivity was observed for hGFRα1-HS recognition **(Fig. 2B)**. The highest binding probes of each length class (DP2, -4, or -6) had heparin-like sulfation patterns, with each disaccharide unit bearing GlcN N- and 6-O- sulfation and IdoA 2-O-sulfation. This indicates that this “even” pattern of sulfation, which is likely to be common in the highly sulfated regions of HS, is beneficial for strong binding to GFRα1 GBS. GFRα1-probe binding did not appear to be greatly dependent on the presence of GlcN 3-O sulfation. The binding of GFRα1 to the heparin-like DP8 probe (92) was similar to that of the DP6 probe with the same sulfation pattern (91), indicating that the ligand requirements of the GBS are satisfied by a length of DP6 and longer ligands do not engage a larger surface. Overall, these results establish that GFRα1 has a GBS-mediated, charge-based and selective binding capacity for heparin and the highly sulfated domains of HS.

To confirm the GBS-dependent interaction of GFRα1 with highly sulfated HS, we measured its binding affinity to glycan ligands using microscale thermophoresis (MST)**(Fig. 2C)**. The dissociation constants (*K*_d_) measured by MST for the interaction of GFRα1 with SOS (47±6 nM) agrees well with that previously measured by ITC^22^ (34±9 nM)**(Fig. S4A-E.)**. The binding affinities of GFRα1 GBS ligands display a clear relationship to the length and GAG class of the glycans tested, in agreement with the microarray results. A 38-fold enhancement in affinity was measured between heparin DP4 and DP10, greater than what would be expected solely from the increase in effective concentration of binding sites due to the increase in size between the two ligands (*K*_d_ ∝ 1/DP), implying that the longer ligands interact with a larger surface of GFRα1. DP6-91, the highest binding probe from the sequence-defined HS microarray, binds with a *K*_d_ of 29±6 nM, a 6-fold enhancement from the naturally derived heparin DP6, which has a 4,5-unsaturated uronic acid on the non-reducing end due to being a product of lyase. Both assayed forms of CS DP10, A/C and D, bound less strongly than heparin DP4, corroborating the results from the GAG NGL microarray. Further comparing the participation of basic GBS residues on D2 in the glycan interaction, hGFRα1 D2-AAA had an 800- and 600-fold weaker affinity for SOS and fondaparinux, a synthetic pentasaccharide anticoagulant based on heparin, respectively compared to WT protein **(Fig. S5A,B)**.

To understand how the selective glycan binding capacity of GFRα1 functions alongside the protein which it binds and serves as a co-receptor for, we further characterised the glycan interactions of GDNF. Mature GDNF is secreted as a disulfide-linked dimer after cleavage of the signal sequence and prodomain (residues 1-77), and retains an unstructured N-terminal tail from residues 78 to 110 **(Fig. S6A)**. Confirming the previous finding that the major determinant for GDNF-HS binding is localised to residues 101-116^21^, targeting this region either by truncation to residue 110 (hGDNF^struc^) or point mutation of 9 lysine and arginine residues primarily in this region (hGDNF^mat^-9A) resulted in a protein with no binding to a heparin-functionalised resin, compared to the 1 M salt concentration required to elute protein with the full mature sequence (78-211) hGDNF^mat^ **(Fig. 2D)**.

We next explored the GAG binding specificity of hGDNF^mat^ by assaying its binding to a different version of the GAG NGL microarray. Similarly to GFRα1, hGDNF^mat^ exhibited a length-dependent preference predominantly for heparin and HS probes, with markedly lower binding of DS-type GAGs and little binding to other GAG probes on the array **(Fig. 2E)**. hGDNF^mat^-9A displayed effectively no binding to any probes, in accordance with its heparin resin retention. A set of chemically desulfated heparin probes showed a graded effect of reducing GDNF binding, from *N*-reacetylation > 6*O*-desulfation > *N*-desulfation ≈ 2*O*-desulfation. These data support a preferential and charge-mediated interaction of GDNF with HS mediated via the binding site in the N-terminal 101-116 region.

Given that GFRα1 and GDNF both engage RET^ECD^ and share an interaction with HS, we investigated whether glycan binding influences assembly of the RET signalling complex using MST. In this assay, RET (hRET^ECD^, residues 29-636)**(Fig. S2B)** was conjugated to a fluorophore and presented with or without hGDNF^mat^ and a variable concentration of hGFRα1 **(Fig. S5C)**. We detected a micromolar interaction between hRET^ECD^ and hGFRα1, which was enhanced to a *K*_d_ of 19±6 nM in the presence of hGDNF **(Fig. S5D,E)**, comparable to the 90±15 nM affinity of the zebrafish RET complex^10^. A modest affinity enhancement was detected in the presence of heparin DP20 (9±3 nM) but not SOS, indicating that the extended nature of the heparin chain was necessary for this effect.

### HS binding induces a conformational shift in GFRα1 D1 domain and the D1-D2 linker

Previous structures of GFRα1 have consistently indicated conformational flexibility between D1 and the D2-D3 module^12,22^, and that different conformational (or proteolytic) states of hGFRα1 are functionally relevant. Considering this alongside the participation of D1 in the GBS alongside D2, we hypothesised that glycan binding might induce a conformational change in GFRα1. To address this, hydrogen-deuterium exchange mass spectrometry (HDX-MS) was used to detect conformational changes in hGFRα1-CHis N59Q with and without fondaparinux. We detected lower deuterium uptake, corresponding to regions more protected from solvent, in the fondaparinux-bound conditions on hGFRα1-CHis N59Q peptides derived from the region between S119 and Y174, corresponding to the D1-D2 linker region and start of D2 **(Fig. 3A, S7)**. This is consistent with a conformational transition from an apo state in which D1 is flexible to an ordered state in which the linker is less solvent-exposed when D1 and D2 form the GBS to engage in sulfated glycan binding. Additionally, the second α-helix of D2, which forms the binding site for GDNF^9,43^ **(Fig. 3B),** overlaps the region found to have reduced exposure in response to sulfated glycan binding.

**Figure 3.**
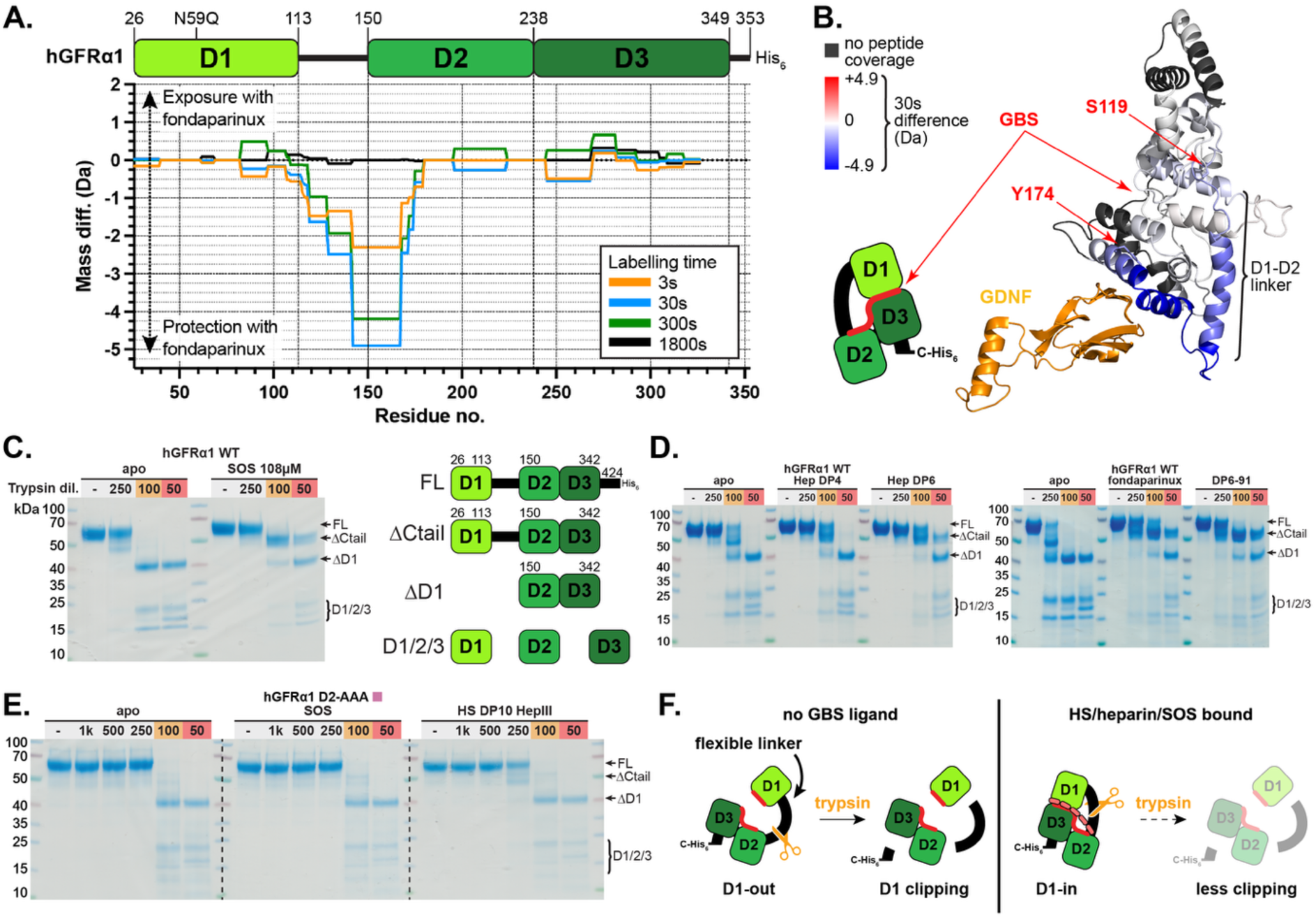
Conformational changes in hGFRα1 upon HS binding tethers the otherwise mobile D1 domain and D1-D2 linker. **(A)** Analysis of the solvent accessibility of the hGFRα1-CHis N59Q peptide backbone in the absence and presence of the heparin pentasaccharide fondaparinux by comparative HDX. For each pair of labelling reaction times, for each residue with detected peptides, the mean mass difference of peptides containing that residue between the apo and +1 molar equivalent fondaparinux conditions is plotted. **(B)** Per-residue protection to solvent exchange of hGFRα1-CHis N59Q caused by the presence of fondaparinux as analysed by comparative HDX for the 30 s labelling condition, mapped on to its structure. Regions where no peptide could be detected in both the apo and liganded conditions are coloured grey. An AlphaFold2 predicted model of GFRα1 was used with HDXViewer. One chain of a hGDNF dimer (PDB: 2V5E) is depicted bound to the binding site on D2. **(C)** hGFRα1-CHis proteolytic digestion pattern monitored by SDS-PAGE following incubation with trypsin at 4°C in the absence and presence of an excess of SOS. The fold dilutions of trypsin from a 1 mg/mL stock is indicated for each reaction. The indicated bands were identified by peptide mapping using relative intensity of peptides derived from each GFRα1 domain. **(D)** SDS-PAGE analysis of hGFRα1-CHis subjected to the limited proteolysis assay in the absence and presence of (left) natural-derived heparin oligosaccharides of DP4 and 6, and (right) synthetic HS oligosaccharides. Apo- and ligand-containing conditions were conducted simultaneously for each replicate. **(E)** SDS-PAGE analysis of hGFRα1-CHis D2-AAA subjected to the limited proteolysis assay in the absence and presence of SOS and natural-derived HS DP10. Band identities are inferred from peptide mapping of the wild type protein. **(F)** A schematic model for the conformational ordering of GFRα1 caused by binding of sulfated glycans to the GBS and the consequent protection from D1 clipping in the limited proteolysis assay.

To support our findings on the conformational dynamics of GFRα1 relating to GAG binding at the GBS, we used a limited proteolysis assay, in which GFRα1 was exposed to a varying concentration of trypsin. SDS-PAGE analysis of the limited proteolysis products of hGFRα1 indicated a protective effect of SOS^9^, in which bands identified as containing an intact D1-D3 unit remained even at the highest trypsin concentration used, compared to complete cleavage of D1 in the absence of a glycan ligand **(Fig. 3C)**. Further assays revealed that a heparin ligand of at least pentasaccharide length was necessary and sufficient for the D1 protective effect in this assay **(Fig. 3D)**, suggesting that the GAG ligand requirements for the conformational change are similar to those for high-affinity binding to the GFRα1 GBS. GBS-targeting hGFRα1 mutations (D1-AAA or D2-AAA) abolished the protective effects of either SOS or a HS DP10 ligand **(Fig. 3E)**, demonstrating the conformational change is dependent on GBS-mediated ligand binding. These data corroborate a model in which a flexible D1 adopts a more compact state upon GAG ligand binding and assembly of the two sections of the GBS.

### Cryo-EM structure of the RET^ECD^-GDNF^mat^-GFRα1-heparin DP20 quarternary complex reveals an extended, multi-protein glycan binding site

To capture the RET ternary complex bound to a HS-type ligand, a reconstituted complex was assembled consisting of hRET^ECD^ (residues 29-635), hGDNF^mat^, hGFRα1, and heparin DP20 (hereafter hRGα1-HP20) **(Fig. 4A,B)**. The complex was purified using size exclusion chromatography, eluting as a single peak **(Fig. S8A)**.

**Figure 4.**
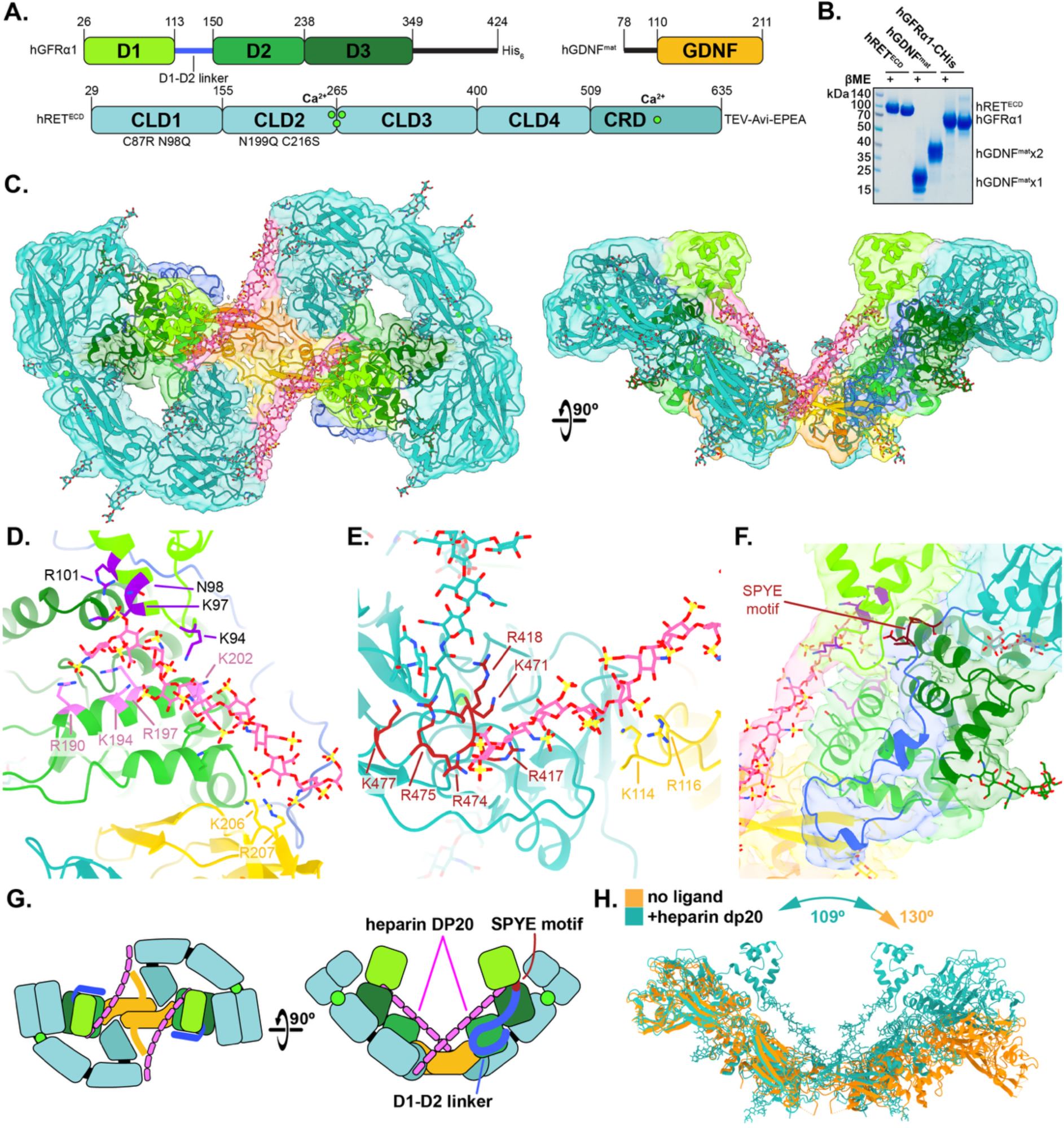
The cryo-EM structure of the hRGα1-HP20 complex reveals an extended multi-protein binding surface for HS-class sulfated glycans. **(A)** Domain diagrams for the recombinant proteins used to reconstitute hRGα1-HP20. **(B)** SDS-PAGE analysis of the recombinant protein preparations before mixing and SEC purification of the hRGα1-HP20 complex, boiled with and without DTT to analyse oligomerisation due to disulfides. **(C)** Orthogonal views of the reconstructed cryo-EM map, and molecular model, viewed down or normal to the symmetry axis. The map is presented at a threshold such that the density corresponding to the heparin ligand is visible and is coloured by proximity to the model. **(D)** Detail view of the interaction site between the heparin ligand at the GFRα1 D1/D2 GBS and with GDNF finger domain residues. **(E)** Detail view of the interaction site between the heparin ligand at the partially resolved GDNF tail and RET CLD4. **(F)** Detail view of the map and model of the hGFRα1 D1-D2 linker (dark blue). **(G)** Schematic representation of the hRGα1-HP20 complex depicting the overall architecture and path of the heparin ligand observed in the structure. **(H)** Comparison of the geometries of the structures of apo (PDB: 6Q2N) and heparin-bound complexes of RET:GDNF:GFRα1, aligned on one RET chain, showing the narrowing of the angle formed between the two batwings. Batwing angles were calculated by measuring the torsion angle formed between residues 61 and 480 in the two RET protomers.

Data collection of cryo-EM samples indicated the presence of ternary complexes with good distribution of particle orientations **(Fig. S8B, S10)**, indicating that the addition of the heparin ligand improved the view anisotropy encountered in previous cryo-EM studies on RET complexes^10,12^. Processing of the reconstructed 3D map with CryoSPARC^44^, as well as Bayesian polishing with RELION-3^45^ **(Fig. S9)**, allowed for determination of a map at nominal resolution (gold standard Fourier shell correlation > 0.143) of 2.74 Å for the whole 2:2:2 complex **(Fig. 4C**, **Table 1, S10)**, and 2.63 Å for local refinement using a focus mask enclosing approximately one copy of each of the 3 protein chains. This resolution improvement upon local refinement reflects flexibility between the two halves of the complex, where flexing largely within the GFL^22^ allows for the wings of the complex to adopt a range of angles, as previously observed in the cryo-EM structure of the zebrafish RET-GDNF-GFRα1 complex^10^. Subjecting the hRGα1-DP20 particle stack to 3D classification revealed this heterogeneity, directly indicating this flexibility.

**Table 1.**
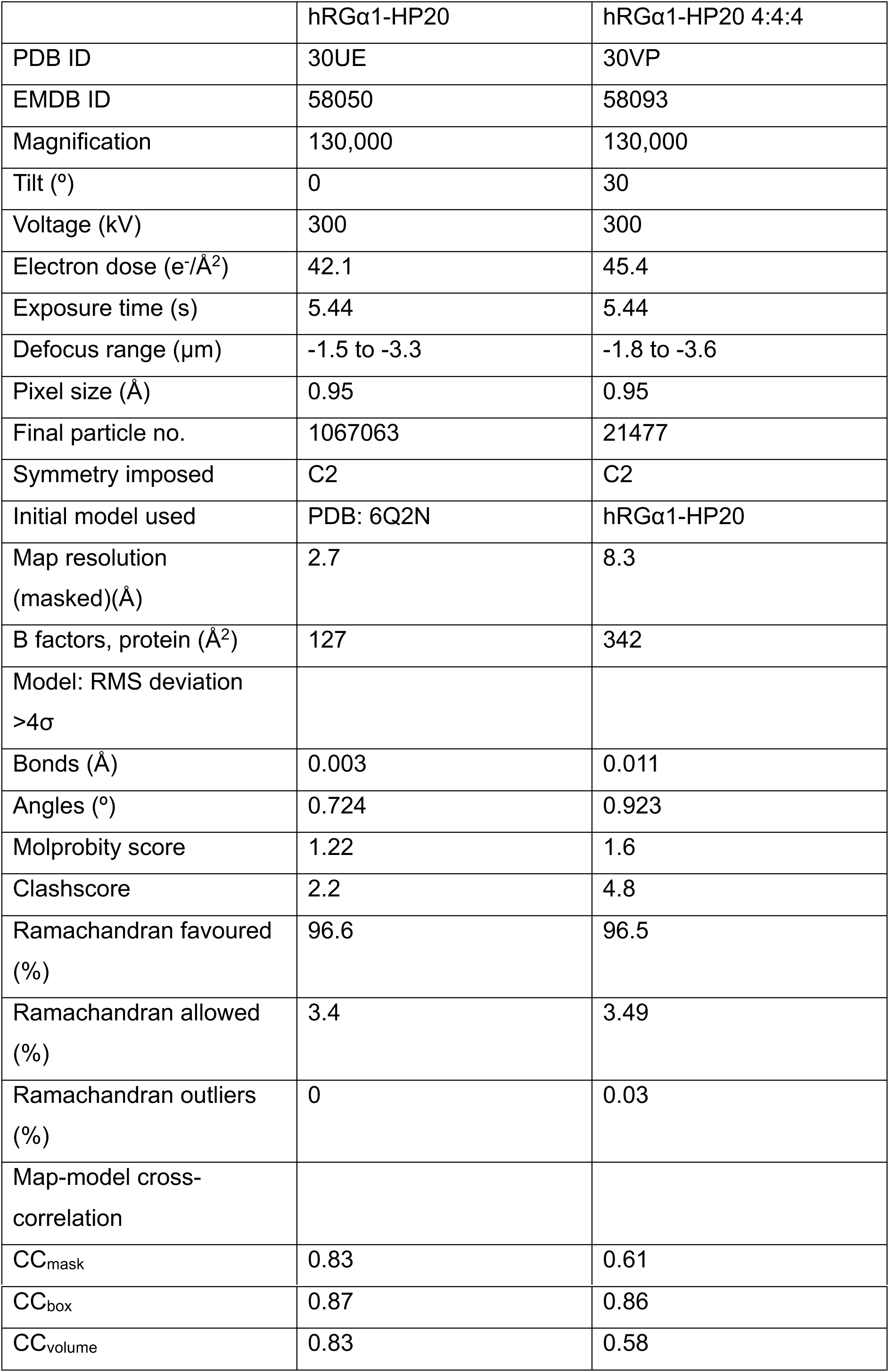
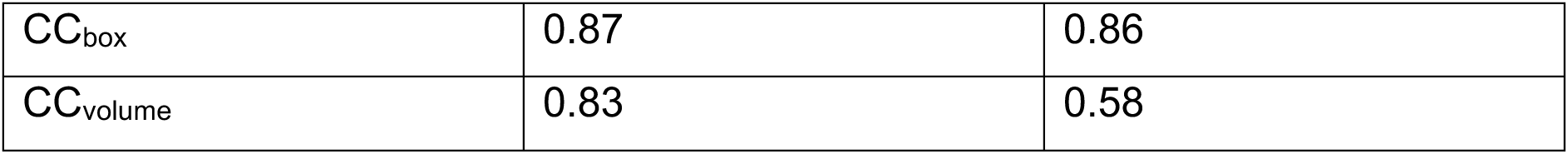
Cryo-EM data acquisition, processing and refinement statistics.

In the heparin bound structure, hRGα1-DP20 adopts the same figure-of-eight architecture previously determined for the glycan ligand-free complex^11^. Improved map density, partially due to the low view anisotropy, allowed us to build a molecular model of the protein based on a previous structure (PDB: 6Q2N) with real-space refinement in PHENIX^46^. The map has a region of elongated density not attributable to protein, present in both halves of the 2:2:2 complex, which extends from the hGFRα1 GBS, contacting the second GDNF finger domain and the GDNF N-terminus, and terminating at the “lower” face of hRET^ECD^ CLD4, where the overall geometry of the ECD turns ∼90° before the CRD (“elbow” region). We attribute this density to the heparin ligand bound to an extended glycan binding site formed across all three protein components of the complex, interacting with basic sidechains at the GFRα1 GBS, hGDNF N-terminal segment, and a previously unreported HS/heparin binding site on CLD4. The heparin in the map has a decreased local resolution (4-6 Å)**(Fig. S10C)**. The dimensions of the density permitted fitting a heparin ligand with 17 saccharide units with 3 sulfates per disaccharide based on an NMR solution structure of a heparin octasaccharide^47^ (PDB: 1HPN). Consistent with the relatively similar presentation of negative charge of heparin/HS S-domains when the molecule is oriented in either direction^47^, heparin can be fit into the map density in either direction, implying that the binding site can accommodate both directionalities of the ligand and that the particle stack used for 3D reconstruction is likely a sampling of combinations of directions.

Density corresponding to 12 N-glycans (per 1:1:1 half-complex) were evident in the hRGα1-HP20 map, permitting molecular modelling of most of these up to the first mannosyl residue to reveal the pose they adopt in the complex. The α1-6 antennary mannosyl residue of the conserved RET N361 glycan appears to make a contact with the loop formed between the second and third helices of GFRα1 D2, forming a previously uncharacterised glycan interaction surface between the ectodomain and co-receptor **(Fig. S12A,B)**. The heparin-bound structure shows an improved interpretability of the map compared to the GAG-free structure, in particular close to CLD1. This improvement reveals a small interaction surface between hGFRα1 D3 sidechain R272 and a CLD1 pocket formed between L119, S148 and F150 **(Fig. S12C)**, in a similar location to but smaller than the interaction site formed between GFRAL D3 and CLD1^12^.

The hRGα1-HP20 cryo-EM map exhibits density of sufficient quality for hGFRα1 D1 and the D1-D2 linker to permit an atomic model to be built for these regions, allowing for the structure of the complete GAG ligand-bound GBS in complex with a physiologically relevant ligand to be determined **(Fig. 4D)**. Comparison with the co-crystal structure of GDNF, GFRα1 ΔD1 and SOS^9^ (PDB: 2V5E) shows that the heparin ligand binds further towards the top of the view of the protein compared to SOS **(Fig. S12D)**, likely due to the additional participation of D1 GBS residues in the full D1-D3 construct used for the hRGα1-HP20 structure compared to the truncated protein used in the SOS structure. The heparin ligand at the hGFRα1 GBS is bound by the D1 loop-helix α4 and D2 helix α3, with additional contributions by K169 and Y170 on D2 helix α2. The ligand also contacts K206 and R207 on the second finger domain of GDNF, forming a nearly-continuous track of surface-exposed positive charge over the GDNF:GFRα1 unit to which heparin/HS ligands can bind.

The D1-D2 linker in the hRGα1-HP20 structure emerges from the D1:D3 interface, packing against the RET-distal face of the D2-D3 module, before extending close to the GDNF interaction site and doubling back towards the first helix of D2 **(Fig. 4F)**. The interaction of the linker close to the GDNF binding site may contribute to the stronger interaction of GDNF with D1-D3 constructs of GFRα1 compared with ΔD1 protein^48^. The start of the D1-D2 linker containing the conserved SPYE motif (residues 114-125) **(Fig. S1A)** is close to hRET^ECD^ CLD1 but makes no contacts with it, consistent with the finding that ΔD1 GFRα1 constructs are capable of forming ternary complexes^48^ and potentiating RET activation by GDNF^49^. Similarly to D1-D3 models of GFRα2^50^ and the zebrafish RET-GDNF-GFRα1 complex^10^, the SPYE motif is located centrally within the D1:D3 interface and projects the Y121 sidechain into a cavity formed between the D1 GBS loop and the interacting face of D3, where it forms a H-bond with R259, part of a RSR motif which is also conserved between D1-containing GFRαs. This central interaction at the D1:D3 interface likely makes the SPYE motif important in limiting the flexibility of the linker when a ligand is bound at the GBS.

### A higher-order assembly of the RET-GDNF-GFRα1 complex form a continuous extended GAG-binding surface

During 2D classification of particles in the hRGα1-HP20 dataset, we noticed a class corresponding to a higher order 4:4:4 assembly, featuring a “side-to-side” arrangement such that the side of a RET CLD4 domain contacts GFRα1 from the adjacent ternary complex **(Fig. 5A**, **Table 1)**. This complex resembles the tetrameric RET:NRTN:GFRα2 complex previously observed by cryo-EM^12^. As the images of the higher order complex displayed a high degree of view anisotropy approximately incident with the C2 dyad, we were unable to perform 3D reconstruction of the assembly from this dataset. To remedy this, we collected additional images at a stage tilt of 30° to obtain additional views of the complex, enabling 3D reconstruction of a map at 8Å resolution of a particle stack from the tilted and untilted datasets.

**Figure 5.**
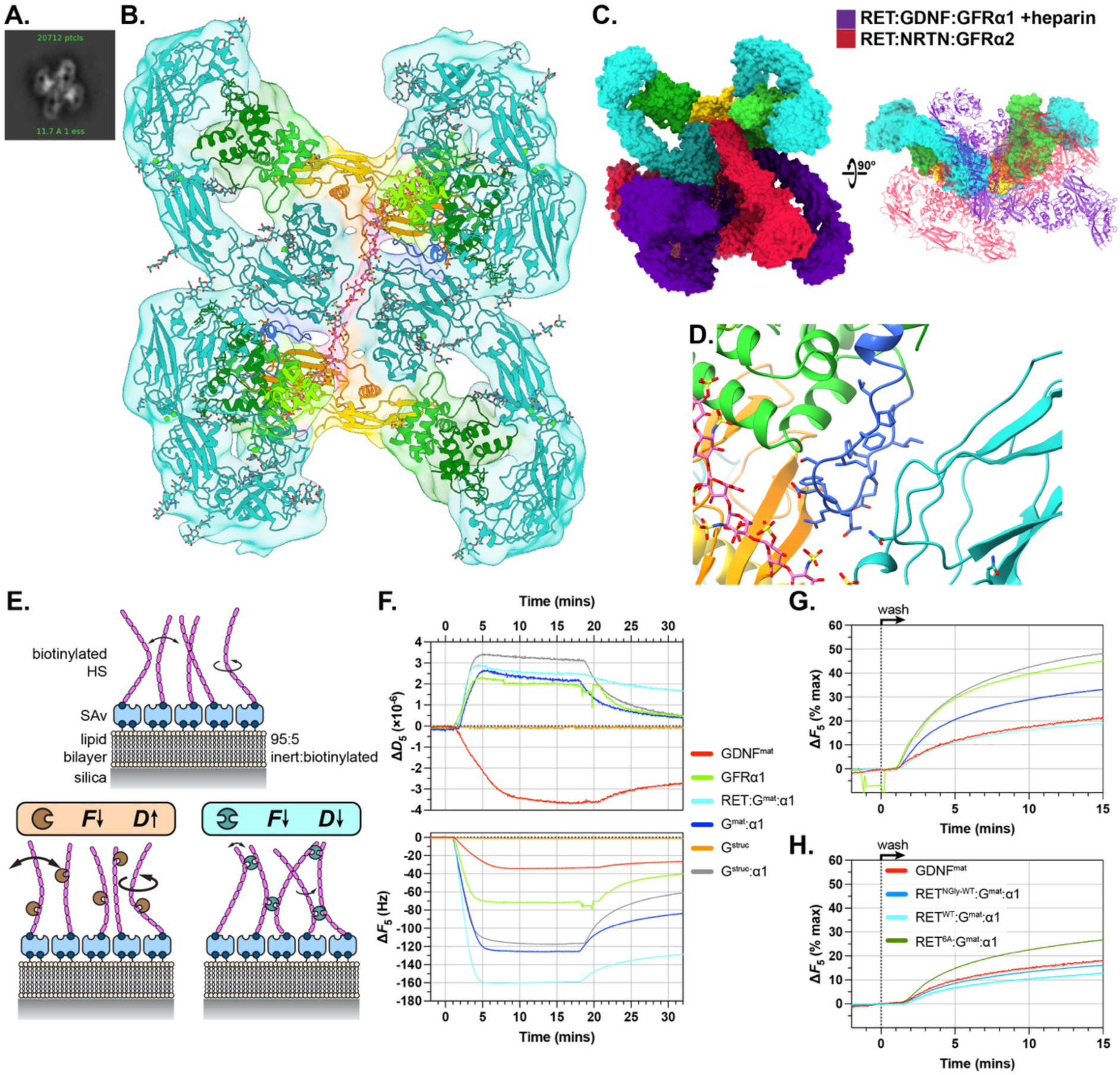
Longer HS chains permit assembly of a higher-order, tetrameric hRGα1 complex linked by binding to a single glycan chain and a GFRα1 linker-RET CLD4 contact. **(A)** 2D class averages of particles extracted from hRGα1-HP20 cryo-EM micrographs (non-tilted collection) containing a heterotetrameric complex. **(B)** Top view of the hRGα1-HP20 tetrameric map and model reconstructed from the combined tilted and untilted datasets. The map is coloured according to proximity to the model. **(C)** Comparison of the inter-complex tilts found in this structure and the tetrameric RET-NRTN-GFRα2 complex (PDB: 6Q2R). The structures were aligned on one 2:2:2 complex **(D)** Inset view hGFRα1 D1-D2 linker-RET CLD4 interaction. **(E)** Schematic of the HS MCS on the QCM-D sensor, alongside binding of a monovalent multivalent HS ligands and their expected effects on *F* and *D*. **(F)** QCM-D binding assays of components and complexes of the RET signalling system to the MCS. For each assay, the start of the binding phase is aligned. The beginning of each desorption phase varies due to the different timing of each assay. **(G)** Plot illustrating the desorption kinetics from the MCS. The function of Δ*F* vs time was aligned to the start of the desorption phase and normalised to 0% = Δ*F*_max_ at t = 0, and 100% = Δ*F* value of unliganded MCS. **(H)** Plot illustrating the desorption kinetics from the MCS including ternary complex formed with hRET^ECD^-6A.

Enforcing a C2 symmetry in the refinement improves the map density, but clear density for both ternary complexes which form the overall tetrameric complex is present in maps from refinement without symmetry. Analysis of the 3DFSC for this map indicates a limited resolution in the direction aligned with the dominant view angle as expected.

The 4:4:4 hRGα1-HP20 map features a non-protein density corresponding to a single heparin ligand bound to the extended glycan binding surfaces of the two interacting 2:2:2 complexes. No discontinuities are evident in this density, implying that it is occupied by a single DP30 heparin. A minority of DP30 heparin may be present in the preparation of heparin used due to the imperfect separation of different lengths when partially digested heparin is separated by HPLC. The configuration of heparin places a GFRα1 GBS interaction on each end of the heparin chain, and a sandwich-like interaction with two CLD4 surfaces at the centre. We docked one copy of each of the RGα-HP20 models with opposite heparin directionalities into this map, permitting building of a complete 4:4:4 model with a single DP30 heparin ligand **(Fig. 5B)**. No significant difference in the batwing angle is evident between the single 2:2:2 hRGα1-HP20 complex and the 4:4:4 complex, suggesting the narrowing in the angle upon binding of the glycan ligand may “prime” the complex for formation of this higher order assembly. Density for heparin bound to the copies of GFRα1 distal to the inter-complex interaction was not evident, and accordingly D1 or the D1-D2 linker on these protomers is not resolvable in the map.

The D1-D2 linker of the “inner” copies of GFRα1 is positioned such that it likely forms a protein-protein interaction surface with the outer face of RET CLD4 **(Fig. 5D)**, in addition to the interaction mediated via simultaneous interaction of the two 2:2:2 complexes with the single heparin chain. This face of CLD4 is not N-glycosylated, permitting the close contact with GFRα1 compared to the heavily glycosylated outer face of CLD3. The arrangement formed between the 2:2:2 complexes in the tetrameric assemblies of RET bound to GDNF:GFRα1:heparin and NRTN:GFRα2 differ **(Fig. 5C)**, with the latter complex tilting such that the glycan interaction site of the CLD4 domain is oriented towards the finger domains of NRTN and the outer face is exposed to solvent, compared to a shallower ∼30° tilt between the two complexes in the GFRα1-GDNF:heparin-bound assembly.

### Formation of hGDNF^mat^ complexes with GFRα1 and hRET^ECD^ alters the modality and kinetics of its binding to HS

HSPGs are critical for the travel and formation of concentration gradients of GAG-binding proteins in the ECM, but little is known about the effect of HS on the diffusion of GDNF and/or soluble forms of GFRα1. When expressing RET signalling system components in HEK293F cells, we noticed that while hGFRα1 readily accumulated in the conditioned media (CM), only HS-binding deficient constructs of hGDNF produced recombinant protein **(Fig. S6B)**. HS-binding hGDNF^mat^ only appears in the CM if soluble heparin is added to the media **(Fig. S6E)**, suggesting that the hGDNF^mat^ expressed by the cells remains bound to cell surface HSPGs and heparin allows it to exchange into solution, leading to accumulation of the protein in the CM.

To investigate this behaviour further, a model cell surface (MCS) on a fluid-supported lipid bilayer (SLB) was reconstituted with HS *in vitro* and used to analyse the binding and release of purified proteins. We used quartz crystal microbalance with dissipation (QCM-D) to monitor protein adsorption (negative frequency shift, Δ*F*) and protein-mediated rigidification (negative dissipation shift, Δ*D*) of the film of end-grafted HS on the MCS **(Fig. 5E)**. Both GFRα1 and GDNF bound to the HS film, but caused opposite dissipation shifts, with GFRα1 softening the film and GDNF inducing a stiffening **(Fig. 5F, S14B.)**. This indicates a different binding mode between the two proteins as reflected by their structures: GFRα1 is monomeric with a single GBS, while GDNF is dimeric with two spatially separated HS binding sites and can therefore crosslink and stiffen the HS film. While the N-truncated hGDNF^struc^ construct did not adsorb to the MCS, the combination of hGDNF^struc^ and GFRα1 induced a greater Δ*F* shift than hGFRα1 alone, indicating that increased binding occurs, likely due to the formation of the 2:2 complex even when the GDNF HS binding site is removed. Combinations of hGDNF^mat^, hGFRα1-CHis and hRET^ECD^ bound to the MCS but had no marked stiffening effect, indicating that GDNF:GFRα1 and ternary complexes bind to the matrix in a different mode to GDNF alone. This is illustrated by a parametric plot, where hGDNF^mat^ causes greater rigidification of the film at all surface densities compared to GFRα1 and complexes of GDNF, which share broadly similar trajectories **(Fig. S14D)**. Comparing the rates of desorption from the HS film reveals further differences in binding kinetics between the different proteins and their complexes **(Fig. 5G, S14C)**. During the desorption step in which buffer is flowed over the sensor, hGDNF^mat^ desorbed from the film at a slow rate, while hGFRα1 did so at a comparatively faster rate. The desorption of hRET^ECD^:hGDNF^mat^:hGFRα1 occured at a similar rate to hGDNF^mat^ alone, suggesting that the additional CLD4-HS contacts, or formation of the tetrameric complex identified by cryo-EM, when all three protein components are present has the effect of reducing the off-rate of the complex compared to the 2:2 complex. A ternary complex formed using RET with mutations targeting six CLD4 residues which contact HS (R417A/R418A/K471A /R474A/R475A/ K477A, hRET^ECD^-6A) desorbed from the HS film at a faster rate than complexes containing hRET^ECD^ protein with WT CLD4 domains **(Fig. 5H, S14E)**, indicating the RET CLD4-HS binding site described here increases the affinity of the complex to HS by slowing down the unbinding kinetics. These data reveal striking differences in the way different protein assemblies of GDNF interact with HS chains in comparison to GDNF alone.

### Soluble GDNF-GFRα1 requires cell surface HS and intact GAG binding sites for RET activation in SH-SY5Y neurons

To validate the function of the GDNF-GFRα1 binding site for HS GAGs *in cellulo*, we used two paradigms of endogenous RET signalling: the breast cancer cell line MCF-7^51,52^ and neuroblastoma line SH-SY5Y^53,54^.

For these experiments, cultured SH-SY5Y cells were differentiated with a 2-phase protocol in which the basal culture is exposed to all-trans retinoic acid, followed by addition of the TrkB agonist ZEB85^55^. This drives the SH-SY5Y cells into a neuronal phenotype in which RET is upregulated^54,56^. The differentiated SH-SY5Y cells express GFRα3 but not GFRα1, meaning they are responsive to GDNF only when presented with soluble GFRα1 but not to either protein alone, via the *in trans* signalling paradigm^25^. RET activation by GDNF:GFRα1 was monitored by quantifying downstream Akt phosphorylation. To establish the role that HS plays in RET activation at the cell surface, we treated differentiated SH-SY5Y with heparinase III to selectively remove cell surface HS, then added GDNF:GFRα1. Treatment with heparinase desensitised the cells to WT GDNF/GFRα1 signalling and to combinations of the GBS-targeting mutants of each protein **(Fig. 6B)**, demonstrating that HS facilitates *in trans* RET activation in this system.

**Figure 6.**
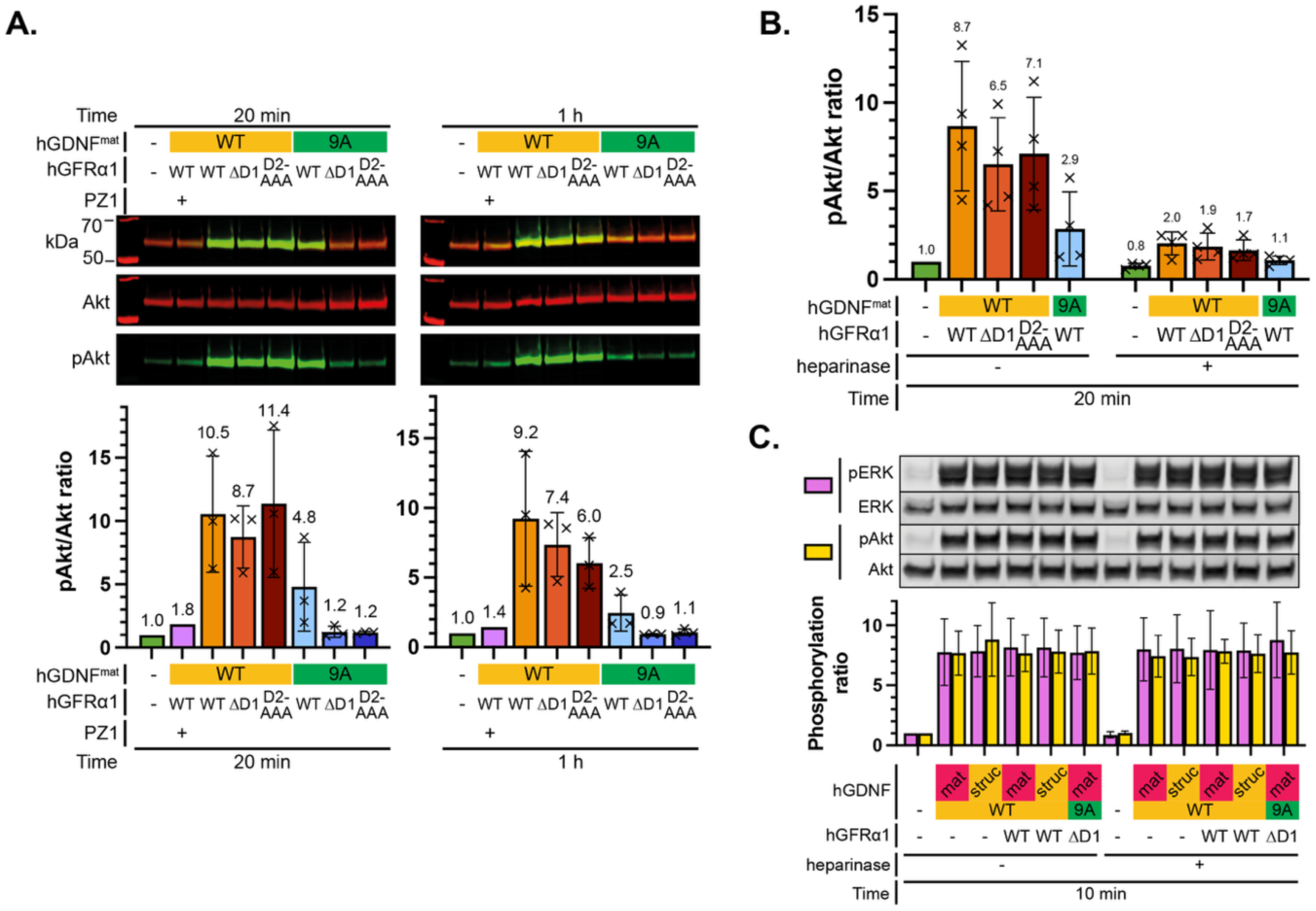
The binding of ligands to cell-surface HS is required for the activation of RET on cells *in trans* but not *in cis*. **(A)** Stimulation of AKT phosphorylation in differentiated SH-SY5Y cells by soluble hGDNF and hGFRα1 protein at 7 nM for 20 min and 1 h. Total AKT and pAKT in cell lysate were analysed by western blotting. Quantification of the phosphoprotein/protein signal intensity ratios normalised to the media-only control for each biological replicate (n=3), represented as the mean ± SD. **(B)** Stimulation of Akt phosphorylation in differentiated SH-SY5Y cells pretreated with heparinase III by soluble hGDNF and hGFRα1 for 20 min. **(C)** Stimulation of Akt and ERK phosphorylation in MCF7 cells with and without pretreatment with heparinase III.

We exposed differentiated SH-SY5Y cells to WT or glycan binding-impaired mutants of GDNF and GFRα1 for different periods and monitored RET activation. hGDNF^mat^ and hGDNF^mat^-9A were chosen to examine the effect of eliminating the glycan binding of full-length mature GDNF. At 20 min and 1 h timepoints, Akt phosphorylation was elevated for all conditions containing hGDNF^mat^ WT with hGFRα1 **(Fig. 6A)**. Substantially lower phosphorylation was observed for the conditions containing hGDNF^mat^ 9A with hGFRα1 WT, while Akt phosphorylation was effectively abolished when hGDNF^mat^ 9A was added alongside either hGFRα1 D2-AAA or ΔD1. Treatment of the cells with the RET-selective inhibitor PZ1^57^ mostly blocked Akt phosphorylation in response to hGDNF^mat^ with hGFRα1. These results indicate that while the GDNF HS binding site has a crucial impact on RET activation, the glycan binding of both proteins is required for full RET activation by GDNF:GFRα1 *in trans*, and the composite GBS across both GDNF and GFRα1 is necessary for the facilitative effect that HS binding plays in receptor activation.

To assess the contribution of HS to RET signalling in an *in cis* context where the GFRα component of the signalling complex is anchored to the same membrane as the RET, we used cultures of MCF-7 cells, which express GFRα1 and RET and respond to GDNF without soluble GFRα1. The response, quantified by downstream Akt and ERK phosphorylation, of WT and HS binding-impaired hGDNF was essentially identical **(Fig. 6C)**. The addition of soluble hGFRα1 constructs had no effect on either downstream response to RET activation, demonstrating that GDNF *in cis* at this concentration results in a saturated RET response, and addition of soluble co-receptor does not result in additional downstream activation. Assaying MCF-7 cells after exposure to the same panel of proteins following pretreatment with heparinase III did not change the response. The activation in the absence of cell surface HS, as well as the loss of downstream kinase phosphorylation when RET knockout MCF-7 cells were assayed **(Fig. S13C)**, excludes the possibility that the signalling response is due to a RET-independent receptor or direct HSPG interaction of GDNF^58^. Together, these results validate the HS binding site observed by cryo-EM and demonstrate that signalling *in trans* requires cell-surface HS on the SH-SY5Y cells. In contrast, signalling *in cis* where GDNF is added to MCF-7 cells displaying RET and GFRα1 anchored on the same cell surface is not dependent on HS for RET activation. We conclude that a major difference in ligand capture mechanism operates between *cis* and *trans* signalling through RET, and that that the latter is HS-dependent.

## Discussion

The critical role of HS in morphogen gradients and signalling during development has been long established but the underpinning mechanisms remain less clear. Studies on FGF, Wnts, Shh and even chemokines have all shown binding sites exist for short HS-chains. However, insights into the impact of HS-glycan interaction on morphogen signalling has been limited, due to the lack of tools to manipulate or image HS-glycan chains by biochemical or structural means. The availability of fractionated heparins as surrogates of HS-chains has shown their flexible nature and mechanical properties, their roles in ligand transportation and clustering of ligand-receptor complexes. Here we describe the extended glycan binding site of the RET:GDNF:GFRα1 signalling complex, providing a structural rationale for existing evidence that HS plays an important role in multiple aspects of RET activation.

HS binds all three components of the RET signalling complex simultaneously, making it likely that HSPGs function as a co-receptor which is associated with the active ligand-receptor assembly in a quaternary complex, as well as participating in ligand capture to the plasma membrane. That the presence of cell-surface HS and HS binding-capable ligands was found to be required for RET activation of GFRα1/GDNF *in trans*, but not by GDNF *in cis*, implies that there is some difference in ligand capture and/or receptor activation between the two regimes, or that HS binding compensates for the loss of anchored GFRα1 in some way. One possibility is that activation requires *in cis* linkage of RET to a GPI anchor to facilitate translocation of the receptor to lipid rafts. Glypican-class HSPGs have HS sites situated spatially close to the membrane (between the folded domain and C-terminal GPI anchor) (in contrast to syndecans, which have GAG attachment sites close to the N-terminus^59^), making them a likely binding partner if translocation of signalling complexes to rafts is indeed important. Equally, binding to cell-surface HSPGs is likely to capture unanchored GDNF:GFRα1 complexes and restrict their motion to 2D diffusion close to the membrane, and the participation of RET in the multi-protein HS binding site may prolong the lifetime of signalling complexes to overcome mechanisms of kinetic proofreading^60^. Further investigation of RET on living membranes is required to build a full model of how HS participates in receptor activation.

While multiple reports have found that sites of RET signalling are exposed to shed GFRα1 *in vivo*^5,24,25,61^, the importance of signalling *in trans* relative to signalling *in cis* is not entirely clear. Although “cis-only” mouse knockouts in which GFRα1 is only expressed in RET-expressing cells do not display any of the developmental defects associated with RET loss-of-function, such mice gain ectopic GFRα1 expression in RET+ cells, implying there is a feedback mechanism capable of compensating for the deficit when GFRα1 is not available *in trans*^62^. This also does not discount any role for shed GFRα1 signalling *in trans*, such as activating RET on the cell that produced it in a cell-autonomous manner, binding to GDNF in the ECM and influencing its diffusion, and the formation of RET-independent complexes. In kidney organogenesis, the invading ureteric bud expresses both RET and GFRα1, implying an *in cis* activation mechanism, but the developmental process fails when certain HS modifications are lost^15,17^. This contrasts with our findings in cell culture models of RET activation, suggesting there is some additional complexity *in vivo* that has been difficult to capture. Whether factors such as the role of the ECM or other cell type-specific components account for this difference requires additional study.

The finding that GFRα1 and GDNF have a highly specific interaction with HS/heparin-class GAGs mirrors previous reports that removing cell surface HS, but not CS, inhibits RET activation^63^. We used heparin, which is produced by the same biosynthetic process as HS but has an overall higher degree of sulfation and an accompanying more even pattern of modification to the NS/NA structure of HS^34^. The use of heparin favours a structural approach due to its lower heterogeneity, aiding the reconstitution of the RET:GDNF:GFRα1:glycan structure. While the more physiologically relevant HS ligand has a more distinct domain pattern, structural elements in the RET ternary complex appear poised to accommodate for differential positioning of the highly sulfated domains; including the presence of basic regions along nearly the entire length of the extended GBS, and the HS-binding site of GDNF, which is largely unstructured and can likely accommodate binding sulfate groups at a range of positions.

Investigating hGDNF^mat^-GAG binding specificity revealed a relatively minor reduction of binding for 2O-desulfated HS. This contrasts with the knockout of HS 2O sulfotransferase in mice, which causes the failure of kidney morphogenesis^15^. Equally, while HS 6O sulfotransferase-deficient mice develop normal kidneys^64,65^, 6O-desulfation resulted in a relatively large reduction of GDNF binding in the array. These discrepancies may be explained by differences in the charge density of HS between the two methods, which is reduced by chemical desulfation but is relatively stable when sulfotransferases are ablated *in vivo*^35^. Whether differential patterns of HS sulfation regulates the signalling of RET or other HS-dependent receptors remains to be seen.

HS binding causes a conformational shift in GFRα1, where the otherwise flexible D1 domain forms the composite GBS with D2 upon GAG ligand binding. The region we found to be conformationally dynamic overlaps with the GDNF binding site on D2, rationalising a previous finding that a ΔD1 construct of GFRα1 has a weaker interaction with GDNF in a cell-binding assay (in which cell-surface HS is presumably present) as ΔD1 protein cannot stabilise the GDNF binding site^48^. This property of GFRα1, alongside the structure of the D1-D2 linker, is also consistent with the inhibition of GDNF-GFRα1 mediated cell adhesion by soluble HS. The linker occupies the D3 interface which is used for formation of the pentameric caps of the “barrel” complex which mediates this adhesion^22^, making it unavailable for complex formation in the HS-bound state. As D1 is required for the interaction of GFRα1 with NCAM^66^, it is possible that HS binding also regulates this interaction, placing HS binding as a central regulator for GDNF-GFRα1 complex formation via this conformational switch.

HSPGs are critical in shaping the effects of signalling molecules which bind them, through influencing their diffusion and the concentration gradient that is formed in the extracellular space. HSPGs can either facilitate or restrict the spread of proteins, based on the nature and strength of their interaction^23^. A relevant example to illustrate this principle is the morphogen Hedgehog, which exhibits two HS binding sites that counterintuitively facilitate faster diffusion through a “switching” mechanism between chains^67^. We show that GDNF has different kinetics and modes of binding to HS depending on the complex it participates in, indicating that the spread of RET ligands through tissue may dynamically change based on the combinatorial availability of GDNF and GFRα1 and the formation of 2:2 complexes. This is relevant for clinical applications of these proteins, as the poor spread of GDNF and NRTN from their point of delivery due to their affinity for HS is likely a factor in its low efficacy in human clinical trials^68,69^. Overall, our findings illustrate that HS shapes GDNF signalling even in the stages before the ligands reach RET-expressing cells.

A single, 30-saccharide chain of heparin was found to bind to a tetrameric form of the RET:GDNF:GFRα1 complex, implying that HS chains can organise higher-order assemblies of RET. The figure-of-eight architecture of the RET complex appears poised to tesselate into this assembly, with the GFRα1 D1-D2 linker forming an interaction surface for an additional copy of RET. The structure of a similar 4:4:4 RET:NRTN:GFRα2 complex has been characterised in the absence of a GAG ligand, the formation of which was linked to slower internalisation kinetics potentially due to the local negative curvature imposed upon the membrane by the complex^12^. The different geometries of the two tetrameric complexes may therefore reflect differential effects on internalisation or other aspects of signalling. Speculatively, the functional outcomes of receptor activation by each of the GFRα/GFL pairs may be shaped by the higher order complexes which they form with RET, which are in turn regulated by HS binding. This may function in addition to the observed variations in batwing angle formed by the RET ternary complexes with different GFRα/GFL pairs^12^, resulting in a system where aspects such as signalling kinetics or downstream signalling effectors are shaped by the geometry of the complex which is formed. Notably, RET ECD with a C634 mutation frequently found in multiple endocrine neoplasia type 2 (MEN2A), which aberrantly dimerises the receptor with an intermolecular disulfide leading to constitutive activation, forms a structure which resembles the configuration found in the HS-arranged 4:4:4 RET:GDNF:GFRα1 complex in which the CLD4 domains are close^70^, as opposed to the conserved 2:2:2 architecture in which RET^ECD^ has a separation distance imposed by the GFL **(Fig. S15)**. This suggests that this cancer mutation mimics an activation mechanism utilised by the wild type receptor.

In summary, we establish that HS is indeed a critical component of the RET system, influencing ligand function and receptor activation in multiple contexts and stages. Our work establishes a novel mechanism for GAG-receptor function, in which a single long heparin chain organises the higher-order assembly and clustering of a receptor-ligand complex by mutual protein-glycan and protein-protein interactions, which may also function along similar principles in other receptor systems with HS-binding components. Further work is necessary to build a more complete model rationalising how regulatory processes and the mechanism of RET activation can be explained by structural aspects of the complexes it forms with its ligands and HS.

## Methods

### Protein expression and purification

Proteins were expressed in Expi293F and harvested from conditioned media. Expi293F cells were grown in Freestyle 293 Expression Medium in vented flasks at 37°C, in 8% CO2 humidified atmosphere with shaking at 125rpm. Cells at a density of 1-1.5×10^6^/mL were resuspended in fresh media and transfected with 1 µg plasmid per 1 mL culture using linear polyethylenimine (MW 25,000)(PEI)(Polysciences) at a mass ratio of 1:3 DNA:PEI. Transfected cells were cultured as above for 4-5d before conditioned media was harvested by centrifugation at 600×g at 4°C for 30 min., followed by the addition of imidazole to 10 mM, Tris pH 7.5 to 20 mM and Ni-NTA agarose resin (Qiagen) and incubation with continuous rotation at 4°C for 1-2 h.

Recovered agarose resin was washed with cold 500 mM NaCl 20 mM HEPES (pH 7.8) buffer followed by elution of protein with three serial incubations with 150 mM NaCl, 20 mM HEPES (pH 7.8), 500 mM imidazole buffer, followed by concentration of eluent using a centrifugal concentrator (Amicon Ultra, Merck). Proteins were further purified using size exclusion chromatography, by injection on to a Superdex 200 Increase 10/300 GL column (Cytiva) on 130 mM NaCl, 20 mM HEPES (pH 7.8) (+1mM CaCl_2_ if RET was present)(gel filration buffer, GFB/GFB+Ca). Peak fractions containing protein as monitored by reducing SDS-PAGE were pooled, concentrated and stored long-term at -70°C following flash freezing.

### GAG NGL microarray

GAG-focused NGL microarrays were prepared as previously described^1^. The microarray-printed slides were fit with a 16-pad FAST frame multi-slide plate aligned to the printed probe sets in a FAST slide 16-well incubation chamber (GVS Filter Technology), pre-wet with HPLC-grade water and flicked dry. Arrays were blocked with 3% BSA solution in PBS (Sigma) for 1 h at RT, followed by flick-dry and washing with PBS. Proteins were bound at 60 µg/mL for 90 min at RT, and detected with mouse anti-His antibody (70796-M, Sigma) at 10 µg/mL for 1 h at RT, followed by incubation with a biotin anti-mouse antibody (10 µg/mL), then streptavidin-Alexa Fluor 647 (Molecular Probes) at 1µg/mL, with four PBS washes carried out between overlay steps. After a final wash with HPLC grade water, the slide was dried in the dark at 37°C, and imaged with a GenePix 4300A (Molecular Devices) with a resolution of 10 µm/pixel at a wavelength of 635 nm.

### Sequence-defined HS microarray

Synthetic HS oligosaccharides with reducing-end functionalisation with a primary amine and pentane spacer of length 2- to 8-mer were dissolved in 50 µL buffer (225 mM Na_2_PO_4_ pH 8.5) to a concentration of 100 µM. Microarrays Kwere printed using a sciFLEXARRAYER S3 (Scienon) microvolume dispenser with a PDC80 nozzle (Scienon) on NHS-ester activated glass slides (NEXTERION Slide H, Schott Inc.) in replicates of 6 with 300 pL spot volume, at 20°C, 50% humidity. After overnight incubation, slides were blocked with ethanolamine buffer (5 mM in 50 mM Tris pH 9) for 1 h, rinsed with DI water, spun dry and stored at RT in a desiccator.

Microarrays were incubated in the dark with hGFRα1-CHis (WT, D2-AAA) at 3 µg/mL for 1 h at RT, followed by washing by sequential dipping for 2 min in TSM wash buffer (20 mM Tris-HCl, 150 mM NaCl, 2 mM CaCl_2_, 2 mM MgCl_2_, 0.05% Tween, pH 7.4), TSM buffer (20 mM Tris-HCl, 50 mM NaCl, 2 mM CaCl_2_, 2 mM MgCl_2_, pH 7.4) and DI water twice. Protein was detected after incubation with Alexa Fluor 635 anti-His antibody for 1 h at RT followed by repetition of the slide washing sequence.

Slides were imaged using GenePix 4000B (Molecular Devices) at 635 nm with a resolution of 5 µm/pixel ensuring the array spots remained in the linear response range of the detector. Images were analysed with GenePix Pro 7 (v7.2.29.2, Molecular Devices) with an Excel macro to yield binding results. The highest and lowest-response spots out of each set of 6 replicates of each probe were excluded for analysis. Microarray screening was performed in triplicate.

### MST affinity

hGFRα1-CHis was labelled using His-Tag Labelling Kit, RED-tris-NTA 2^nd^ gen (NanoTemper). The dye was dissolved in PBS-T (Na_2_HPO_4_ 8.1 mM, KH_2_PO_4_ 1.5 mM, NaCl 137 mM, KCl 2.7 mM, 0.05% w/v Tween 20) to 5µM and mixed to a concentration of 50 nM with protein at 250 nM, incubated at RT for 30 min., centrifuged for 10 min. at 13,000×g at 4°C, and the supernatant separated and used. Solutions for loading into capillaries (premium capillaries, NanoTemper) were made by mixing the dye-protein solution 1:1 with solutions of the serially diluted analyte with or without any additional components. Fluorescently labelled hRET^ECD^ (hRET^ECD^*) was produced using Monolith Protein Labeling Kit RED-NHS 2^nd^ generation (NanoTemper) according to the manufacturer’s protocol. The final concentration of the labelled proteins in the capillaries was 125 nM for hGFRα1-CHis, and 50 nM for hRET^ECD^*. MST was performed with a Monolith instrument (NanoTemper) at 25°C. The fluorescence measured transverse across the capillaries showed a bell-shaped curve, indicating the fluorescently labelled proteins did not stick to the glass **(Fig. S4D)**. Excitation and IR power respectively were set to 40%/40% for hGFRα1-CHis experiments and 20%/60% for hRET^ECD^*. MO.Affinity software (NanoTemper) was used to analyse MST data by fitting to a 1:1 binding model to derive a disassociation constant (*K*_d_) with a confidence interval based on the least squares fit of the model to the data. The cold region was set to -1 s to 0 s for all experiments, and the hot region was set to 1.5 s to 2.5 s for hGFRα1-CHis and 14-15 s for hRET^ECD^*. *F*^MST^_norm_ is defined as the ratio of fluorescence at 670 nm between the hot and cold regions and is denoted per mille (‰). Δ*F*^MST^_norm_ is the difference in between the extents of the fit binding model **(Fig S4B, E)**.

### Heparin column affinity

HiTrap Heparin HP 1 mL columns (Cytiva) were used on an AKTA Pure liquid chromatography system (Cytiva). Columns were prepared by equilibration into buffer A (50 mM NaCl, 20 mM HEPES pH 7.8), washing with 5 column volumes of buffer B (2 M NaCl, 20 mM HEPES pH 7.8) then equilibration into buffer A. Protein was loaded by injection to the column, and if protein did not elute on the flow-through (no binding), the percentage of buffer B was increased with monitoring of absorbance at 280 nm and buffer conductivity.

### MCF7 culture and signalling assay

MCF7 cells were seeded at 300,000 cells/well in 12 well plates, in DMEM/10% FBS. The following day cells were washed x1 in warm DMEM, then serum starved for 24 h before ligand application. 2 h before ligand application, 3 AU Heparinase III (#HY-P79211, Cambridge Bioscience) was applied to 6 wells per plate. RET ligands were applied at 10 nM final concentration for 10 min before aspiration of media and addition of 150 ul lysis buffer (50 mM Tris, pH 7.5, 150 mM NaCl, 2 mM EDTA, protease inhibitor cocktail cOmplete EDTA, phosphatase inhibitor cocktail PhosStop) for 15 min on ice. Lysates were centrifuged for 10 min, 4°C, 13000×g. The supernatant was separated and stored at -20°C until analysis.

Samples were thawed on ice, 4x LDS Sample buffer and DTT (final concentration 50 mM) added, then run on SDS-PAGE (4-12% Bis-Tris gels) in MES buffer (27 min, 200V), with the Spectra™ Multicolor Broad Range Protein Ladder (Thermo scientific, Cat No. 26634). Proteins were transferred to nitrocellulose membrane via wet transfer (110V, 70 min), before PonceauS staining. Membranes were blocked in 5% BSA/TBST (1 h, RT). Primary antibodies were applied at the stated concentrations **(Table S5)** in 5% BSA/TBST, and membranes were incubated overnight at 4°C with gentle rocking. Membranes were washed ×3 in TBST (5 min per wash). Secondary antibodies were applied diluted in Blocking Buffer for Fluorescent Western Blotting (#ABIN925618, Antibodies Online)(1 h, RT). Membranes were washed ×3 in TBST, then imaged on a LICORbio Odyssey CLx Imager. Band intensities were analysed using the Image Studio software (LICOR).

### SH-SY5Y culture and signalling assay

SH-SY5Y cells were cultured in Dulbecco’s Modified Eagle Medium F-12 Nutrient Mixture [+] L-Glutamine (DMEM/F-12, Gibco), with 10% Fetal Bovine Serum (FBS, Gibco) and 1% Penicillin-Streptomycin (PenStrep, Gibco), in T-75 flasks incubated at 37°C, 5% CO_2_. They were passaged twice a week at a 1:3 ratio, using 4 mL 0.25% Trypsin-EDTA (1X) (Gibco). The cells were seeded at a 1:4 density in 12 well plates pre-coated with 10 µg/cm^2^ bovine Collagen I (Gibco). A two-step protocol was used to differentiate the cells. On day 0, fresh media was added which was supplemented with 10 µM all trans retinoic acid (ATRA, Sigma-Aldrich R2625). On day 5, the media was changed for DMEM/F-12 and PenStrep media without FBS, supplemented with 1 nM of the TrkB agonist ZEB85. The experiments were performed on day 12: the tested proteins were diluted in DMEM/F-12 and added to the cells, which were incubated a further 20 minutes, 1 hour, or 24 hours. The cells were then lysed on ice for 15 minutes (lysis buffer: 1% NP-40 alternative, 2 nM EDTA, 150 mM NaCl, 50 mM Tris, protease inhibitor cocktail (Roche) and phosphatase inhibitor cocktail (Roche)), centrifuged for 10 minutes, and the cell lysate was analysed by western blot using anti Akt and pAkt primary antibodies (Cell Signalling Technology, clones 40D4 and D9E respectively).

### Comparative HDX

hGFRα1-CHis N59Q, containing residues 26-353, removing an N-glycan site in D1 compared to the WT protein, was used to improve peptide resolution and data analysis in HDX. Purification of hGFRα1-CHis N59Q was similar to other constructs of hGFRα1.

hGFRα1-CHis N59Q was diluted to 5 µM in deuterated (D_2_O) buffer and incubated for 3, 30, 300 and 1800 s before reaction quenching by addition of cold 2.4% v/v formic acid in 2 M guanidinium chloride and immediate flash freezing in liquid nitrogen. Samples were stored at -80°C before analysis. Samples were rapidly thawed and added to pepsin protease, and injected to reverse-phase HPLC on an Enzymate BEH immobilised pepsin column (2.1 × 30 mm, 5 µM, Waters) at 200 µL/min for 2 min. Peptides were trapped and desalted with an Acquity BEH C18 Van-guard pre-column (2.1 mm × 5 mm, 1.7 µM, Waters), and eluted at 40 µM/min for 11 min with a gradient of acetonitrile (5-43%) in 0.1% v/v formic acid. UPLC was performed with an Acquity UPLC BEH C18 (100 mm × 1 mm, 1.7 µM, Waters) with detection using a Cyclic mass spectrometer (Waters) acquiring at m/z 300-2,000 with standard electrospray ionisation source, and lock mass calibration with [Glu1]-fibrinopeptide B at 50 fmol/µL. MS was operated at 80°C source temperature and 3.0 kV spray voltage, and spectra were collected in positive ion mode. The labelling reactions were performed in triplicate at the same time for all timepoints, and individual timepoints were acquired by MS on the same day.

Peptides were identified by MS^e^ (MS scan followed by all-ion fragmentation) using an identical gradient of acetonitrile in 0.1% v/v formic acid over 12 min ^2^. MSe data was analysed with Protein Lynx Global Server (Waters), with an MS tolerance of 5 ppm. Mass analysis of peptide centroids was performed using DynamX (Waters) software, only considering peptides with score >6.4. The first round of peptide identification was done automatically by the software, with additional manual verification of all peptides at all timepoints for correct charge state, presence of overlapping peptides and correct retention time. The derived mass changes were not corrected for back-exchange, and therefore represent relative, not absolute, changes in deuteration of the peptides. Changes in H-D amide exchange in any peptide may be due to one amide or multiple amides.

### Limited proteolysis assay

Trypsin (Proti-Ace) was reconstituted to 1 mg/mL, and diluted in buffer (130 mM NaCl 20 mM, HEPES pH 7.8) to ratios between 1/1000 to 1/10 (concentration range of 1-100 µg/mL). For each reaction condition, 5 µL protein at 1 mg/mL (with or without additional compound, preincubated on ice for 5 min.) was mixed with 5 µL diluted trypsin (or buffer) and incubated for 2 h at 4°C. Reactions were quenched by addition of water and reducing PAGE loading buffer 4X to a total volume of 20 µL, and heating to 95°C for 3 min. Reaction products were analysed by SDS-PAGE.

### Cryo-EM sample preparation

hRET^ECD^, hGFRα1-CHis and hGDNF^mat^ were mixed at 20µM with 2 molar equivalents of heparin DP20 (Iduron) in GFB+Ca and purified using a Superdex 200 Increase 10/300 GL column (Cytiva).

Holey carbon film support, 300 R1.2/R1.3 copper grids (QUANTIFOIL) were glow discharged (45 mA, 45 s). Using a Vitrobot Mark IV (Thermo Fisher Scientific), 4 µL sample (at approx. 0.5 mg/mL) was applied to grids, blotted (100% humidity at RT, 20 s wait time, blot force 0-1, blot time 5 s) and vitrified by plunging into liquid ethane. Grids were clipped, loaded into Nanocab cassettes and screened on a Talos Arctica (200 kV, Falcon 3 detector)(Thermo Fisher Scientific) for suitability for automated data collection. Data collection was controlled with EPU software (version 3.11, Thermo Fisher Scientific) using a Titan Krios G2 (300 kV, Falcon 4i detector)(Thermo Fisher Scientific) microscope with a Selectris energy filter (10 eV slit width). Movies were collected in electron-event representation (EER) format^3^ with 1674 raw frames, which were fractionated to 31 during motion correction

### Preparation of components for the cell surface model (SLB)

Small unilamellar vesicles (SUVs) were prepared using a previously reported protocol^4^. 1,2-dioleoyl-*sn*-glycero3-phosphocho-line (DOPC, Avanti Polar Lipids) and 1,2-dioleoyl-sn-glycero-3-phosphoethanolamine-N-(cap biotinyl) (DOPE-biotin, Avanti Polar Lipids) were mixed to a 95:5 molar ratio in chloroform and deposited as a thin film in a glass vial through evaporation of the solvent with a nitrogen stream while the vial was rotated. The film was then further dried in vacuum (1 h), rehydrated in HBS (10 mM HEPES pH 7.4, 150 mM NaCl in ultrapure water) (final concentration 1 mg/mL). The lipid suspension was homogenised by five cycles of freezing, thawing and vortexing, and converted into SUVs by tip sonication in pulse mode (1 s on/1 s off for 30 min) with refrigeration. The SUV suspension was then centrifuged at 12,100 × *g* for 10 min to remove titanium debris (shed from the sonicator tip), and stored at 4 °C under nitrogen gas until use.

Biotinylated HS (HS-b) was prepared using a previously reported protocol^5,6^ using HS polysaccharides of average molecular weight 33 kDa and 1.6 sulfates/disaccharide (Fraction C as described in reference^7^).

### QCM-D assays

We used a quartz crystal microbalance with dissipation (QCM-D) technology to monitor the formation of model cell surfaces and protein binding through changes in the resonance frequency and damping of a silica-coated QCM-D sensor, which are interpreted as adsorption to the sensor (a decrease in the resonance frequency; negative Δ*F*) and changes to the layer softness (softness entails energy dissipation; positive Δ*D*) respectively^8^. A HS-based fluid-supported lipid bilayer (SLB) was constructed by rupture and spread of biotinylated SUVs on the sensor surface, followed by the sequential addition of a streptavidin monolayer which served as a molecular print board to accommodate a film of end-grafted HS-b **(Fig. 5E, S14A)**.

QCM-D was conducted using a QSense Analyser (Biolin Scientific) with SiO2 coated sensors (QSX303, Biolin Scientific) and QSoft control software, at a working temperature of 23.2°C. Four flow modules were operated in parallel with a custom-made syringe pump system that continuously pulled liquid through the flow modules at a flow rate of 20µL/min. Normalised frequency shifts Δ*F* = Δ*f*/*i* and Δ*D* were collected at six odd overtones (*i* = 3 to 13); data for *i* = 5 are graphed in figures throughout but all overtones gave qualitatively similar results. Measurements were performed in duplicate on separate days with newly constructed model cell surfaces. The sensors were cleaned prior to use by immersion in 2% w/v sodium dodecyl sulfate (SDS)(30 min) followed by rinsing with ultrapure water, blow-drying with nitrogen gas, and exposure to UV/ozone (Jelight Company; 30 min). Following a check for stable Δ*F* and Δ*D* baselines in air and then in HEPES buffered saline with calcium (HBS; 10mM HEPES pH 7.4, 150mM NaCl, 1mM CaCl_2_), the model cell surface was self-assembled on the sensor surface by the following steps with 10 min HBS steps inbetween: SUVs (50 µg/mL, 20 min), streptavidin (SAv)(20 µg/mL, 15 min) to form a dense monolayer on the SLB, and HS-b (10 µg/mL, 15 min) to form a brush of end-grafted polysaccharides with a root-mean-square anchor spacing of approximately 5 nm^9,10^. For each experiment, two of the chambers were not exposed to HS-b for use as negative controls to validate all binding of analytes was specific to the HS. Each analyte solution was prepared in HBS from purified individual proteins immediately prior to use in a QCM-D assay step. Each component was diluted to 680 nM, a concentration equivalent to that of 5 µg/mL of hGDNF^mat^. Following the protein binding step, HBS was flowed over the model cell surface for at least 15 min to monitor the rate of protein unbinding, followed by regeneration of the model cell surface with a high salt buffer (10 mM HEPES pH 7.4, 2 M NaCl).

## Author information

Correspondence and requests for material should be addressed to N. Q. McDonald

## Acknowledgments

We would like to acknowledge the work of the Francis Crick Structural Biology, Chemical Biology, Proteomics and Scientific Computing Science Technology Platforms. We would like to acknowledge the work of Frankie Houghton for guidance on conformational dynamics and for establishing the limited proteolysis assay. We thank Simone Kunzelmann for MST training and assistance with data processing.

We acknowledge the work of Erika Holz (University of Leeds) for providing SUVs and assistance with QCM-D experiments. We thank Barbara Mulloy for advice on HS assays and experimental design. We thank Jean-Paul Vincent and David Willnow for guidance on data interpretation. We acknowledge the advice and support of Erhard Hohenester and Katrin Rittinger.

## Author contributions

M.I.Z.-H. and N.Q.M. prepared and wrote the manuscript. M.I.Z.-H. performed general protein expression and purification, the NGL GFRα1 microarray, MST, cryo-EM sample preparation and structure reconstruction and QCM-D. B.R. performed MCF-7 signalling assays. S.B.-L. performed SH-SY5Y signalling assays. M.Z.-P. assisted with cryo-EM data processing. D.C.B. assisted with structural model refinement. A.N. performed cryo-EM data acquisition. A.B. carried out insect cell culture and expression. S.L.M. performed the HDX experiment and data analysis.

A.M. performed the microarray analysis using the GAG NGL microarrays. J.P. made reagents for the QCM-D model cell surface, and J.P. and R.P.R. assisted with QCM-D experiments and data analysis. All authors reviewed the manuscript.

## References

1. Honma, Y., Kawano, M., Kohsaka, S., and Ogawa, M. (2010). Axonal projections of mechanoreceptive dorsal root ganglion neurons depend on Ret. Development 137, 2319–2328. 10.1242/dev.046995.

2. Kramer, E.R., Aron, L., Ramakers, G.M.J., Seitz, S., Zhuang, X., Beyer, K., Smidt, M.P., and Klein, R. (2007). Absence of Ret Signaling in Mice Causes Progressive and Late Degeneration of the Nigrostriatal System. PLoS Biol. 5, e39. 10.1371/journal.pbio.0050039.

3. Oo, T.F., Kholodilov, N., and Burke, R.E. (2003). Regulation of Natural Cell Death in Dopaminergic Neurons of the Substantia Nigra by Striatal Glial Cell Line-Derived Neurotrophic Factor *In Vivo*. J. Neurosci. 23, 5141–5148. 10.1523/JNEUROSCI.23-12-05141.2003.

4. Schuchardt, A., D’Agati, V., Larsson-Blomberg, L., Costantini, F., and Pachnis, V. (1994). Defects in the kidney and enteric nervous system of mice lacking the tyrosine kinase receptor Ret. Nature 367, 380–383. 10.1038/367380a0.

5. Trupp, M., Belluardo, N., Funakoshi, H., and Ibáñez, C.F. (1997). Complementary and Overlapping Expression of Glial Cell Line-Derived Neurotrophic Factor (GDNF), c-ret Proto-Oncogene, and GDNF Receptor-α Indicates Multiple Mechanisms of Trophic Actions in the Adult Rat CNS. J. Neurosci. 17, 3554–3567. 10.1523/JNEUROSCI.17-10-03554.1997.

6. Costantini, F. (2010). GDNF/Ret signaling and renal branching morphogenesis. Organogenesis 6, 252–262. 10.4161/org.6.4.12680.

7. Meng, X., Lindahl, M., Hyvönen, M.E., Parvinen, M., De Rooij, D.G., Hess, M.W., Raatikainen-Ahokas, A., Sainio, K., Rauvala, H., Lakso, M., et al. (2000). Regulation of Cell Fate Decision of Undifferentiated Spermatogonia by GDNF. Science 287, 1489–1493. 10.1126/science.287.5457.1489.

8. Lin, L.-F.H., Doherty, D.H., Lile, J.D., Bektesh, S., and Collins, F. (1993). GDNF: a Glial Cell Line-Derived Neurotrophic Factor for Midbrain Dopaminergic Neurons. Science 260, 1130–1132. 10.1126/science.8493557.

9. Parkash, V., Leppänen, V.-M., Virtanen, H., Jurvansuu, J.M., Bespalov, M.M., Sidorova, Y.A., Runeberg-Roos, P., Saarma, M., and Goldman, A. (2008). The Structure of the Glial Cell Line-derived Neurotrophic Factor-Coreceptor Complex. J. Biol. Chem. 283, 35164–35172. 10.1074/jbc.M802543200.

10. Adams, S.E., Purkiss, A.G., Knowles, P.P., Nans, A., Briggs, D.C., Borg, A., Earl, C.P., Goodman, K.M., Nawrotek, A., Borg, A.J., et al. (2021). A two-site flexible clamp mechanism for RET-GDNF-GFRα1 assembly reveals both conformational adaptation and strict geometric spacing. Structure 29, 694–708.e7. 10.1016/j.str.2020.12.012.

11. Goodman, K.M., Kjær, S., Beuron, F., Knowles, P.P., Nawrotek, A., Burns, E.M., Purkiss, A.G., George, R., Santoro, M., Morris, E.P., et al. (2014). RET Recognition of GDNF-GFRα1 Ligand by a Composite Binding Site Promotes Membrane-Proximal Self-Association. Cell Rep. 8, 1894–1904. 10.1016/j.celrep.2014.08.040.

12. Li, J., Shang, G., Chen, Y.J., Brautigam, C.A., Liou, J., Zhang, X., and Bai, X.C. (2019). Cryo-EM analyses reveal the common mechanism and diversification in the activation of RET by different ligands. eLife 8, 1–26. 10.7554/eLife.47650.001.

13. Zol-Hanlon, M., and McDonald, N.Q. (2026). RET receptor tyrosine kinase architecture, assemblies, and activation. Endocr. Relat. Cancer 33, e250322. 10.1530/ERC-25-0322.

14. Mullican, S.E., Lin-Schmidt, X., Chin, C.-N., Chavez, J.A., Furman, J.L., Armstrong, A.A., Beck, S.C., South, V.J., Dinh, T.Q., Cash-Mason, T.D., et al. (2017). GFRAL is the receptor for GDF15 and the ligand promotes weight loss in mice and nonhuman primates. Nat. Med. 23, 1150–1157. 10.1038/nm.4392.

15. Bullock, S.L., Fletcher, J.M., Beddington, R.S.P., and Wilson, V.A. (1998). Renal agenesis in mice homozygous for a gene trap mutation in the gene encoding heparan sulfate 2-sulfotransferase. Genes Dev. 12, 1894–1906. 10.1101/gad.12.12.1894.

16. Enomoto, H., Araki, T., Jackman, A., Heuckeroth, R.O., Snider, W.D., Johnson, E.M., and Milbrandt, J. (1998). GFRα1-deficient mice have deficits in the enteric nervous system and kidneys. Neuron. 10.1016/S0896-6273(00)80541-3.

17. Li, J.-P., Gong, F., Hagner-McWhirter, Å., Forsberg, E., Åbrink, M., Kisilevsky, R., Zhang, X., and Lindahl, U. (2003). Targeted Disruption of a Murine Glucuronyl C5-epimerase Gene Results in Heparan Sulfate Lacking l-Iduronic Acid and in Neonatal Lethality. J. Biol. Chem. 278, 28363–28366. 10.1074/jbc.c300219200.

18. Moore, M.W., Klein, R.D., Fariñas, I., Sauer, H., Armanini, M., Phillips, H., Reichardt, L.F., Ryan, A.M., Carver-Moore, K., and Rosenthal, A. (1996). Renal and neuronal abnormalities in mice lacking GDNF. Nature 382, 76–79. 10.1038/382076a0.

19. Schneeberger, P.E., Von Elsner, L., Barker, E.L., Meinecke, P., Marquardt, I., Alawi, M., Steindl, K., Joset, P., Rauch, A., Zwijnenburg, P.J.G., et al. (2020). Bi-allelic Pathogenic Variants in HS2ST1 Cause a Syndrome Characterized by Developmental Delay and Corpus Callosum, Skeletal, and Renal Abnormalities. Am. J. Hum. Genet. 107, 1044–1061. 10.1016/j.ajhg.2020.10.007.

20. Ai, X., Kitazawa, T., Do, A.T., Kusche-Gullberg, M., Labosky, P.A., and Emerson, C.P. (2007). SULF1 and SULF2 regulate heparan sulfate-mediated GDNF signaling for esophageal innervation. Development 134, 3327–3338. 10.1242/dev.007674.

21. Alfano, I., Vora, P., Mummery, R.S., Mulloy, B., and Rider, C.C. (2007). The major determinant of the heparin binding of glial cell-line-derived neurotrophic factor is near the N-terminus and is dispensable for receptor binding. Biochem. J. 404, 131–140. 10.1042/BJ20061747.

22. Houghton, F.M., Adams, S.E., Ríos, A.S., Masino, L., Purkiss, A.G., Briggs, D.C., Ledda, F., and McDonald, N.Q. (2023). Architecture and regulation of a GDNF-GFRα1 synaptic adhesion assembly. Nat. Commun. 14, 7551. 10.1038/s41467-023-43148-8.

23. Stapornwongkul, K.S., and Vincent, J.-P. (2021). Generation of extracellular morphogen gradients: the case for diffusion. Nat. Rev. Genet. 22, 393–411. 10.1038/s41576-021-00342-y.

24. Fleming, M.S., Vysochan, A., Paixão, S., Niu, J., Klein, R., Savitt, J.M., and Luo, W. (2015). Cis and trans RET signaling control the survival and central projection growth of rapidly adapting mechanoreceptors. eLife 4, e06828. 10.7554/eLife.06828.

25. Ibáñez, C.F., Paratcha, G., and Ledda, F. (2020). RET-independent signaling by GDNF ligands and GFRα receptors. Cell Tissue Res. 382, 71–82. 10.1007/s00441-020-03261-2.

26. Ledda, F., Paratcha, G., and Ibáñez, C.F. (2002). Target-derived GFRα1 as an attractive guidance signal for developing sensory and sympathetic axons via activation of Cdk5. Neuron 36, 387–401. 10.1016/S0896-6273(02)01002-4.

27. Encinas, M., Tansey, M.G., Tsui-Pierchala, B.A., Comella, J.X., Milbrandt, J., and Johnson, E.M. (2001). c-Src Is Required for Glial Cell Line-Derived Neurotrophic Factor (GDNF) Family Ligand-Mediated Neuronal Survival via a Phosphatidylinositol-3 Kinase (PI-3K)-Dependent Pathway. J. Neurosci. 21, 1464–1472. 10.1523/JNEUROSCI.21-05-01464.2001.

28. Tansey, M.G., Baloh, R.H., Milbrandt, J., and Johnson, E.M. (2000). GFRα-mediated localization of RET to lipid rafts is required for effective downstream signaling, differentiation, and neuronal survival. Neuron 25, 611–623. 10.1016/S0896-6273(00)81064-8.

29. Tsui, C.C., Gabreski, N.A., Hein, S.J., and Pierchala, B.A. (2015). Lipid rafts are physiologic membrane microdomains necessary for the morphogenic and developmental functions of glial cell line-derived neurotrophic factor in vivo. J. Neurosci. 35, 13233–13243. 10.1523/JNEUROSCI.2935-14.2015.

30. Paratcha, G., Ledda, F., and Ibáñez, C.F. (2003). The Neural Cell Adhesion Molecule NCAM Is an Alternative Signaling Receptor for GDNF Family Ligands. Cell 113, 867–879. 10.1016/S0092-8674(03)00435-5.

31. Sergaki, M.C., and Ibáñez, C.F. (2017). GFRα1 Regulates Purkinje Cell Migration by Counteracting NCAM Function. Cell Rep. 18, 367–379. 10.1016/j.celrep.2016.12.039.

32. Paratcha, G., Ibáñez, C.F., and Ledda, F. (2006). GDNF is a chemoattractant factor for neuronal precursor cells in the rostral migratory stream. Mol. Cell. Neurosci. 31, 505–514. 10.1016/j.mcn.2005.11.007.

33. Ledda, F., Paratcha, G., Sandoval-Guzmán, T., and Ibáez, C.F. (2007). GDNF and GFRα1 promote formation of neuronal synapses by ligand-induced cell adhesion. Nat. Neurosci. 10, 293–300. 10.1038/nn1855.

34. Xu, D., and Esko, J.D. (2014). Demystifying heparan sulfate-protein interactions. Annu. Rev. Biochem. 83, 129–157. 10.1146/annurev-biochem-060713-035314.

35. Merry, C.L.R., Bullock, S.L., Swan, D.C., Backen, A.C., Lyon, M., Beddington, R.S.P., Wilson, V.A., and Gallagher, J.T. (2001). The Molecular Phenotype of Heparan Sulfate in theHs2st−/− Mutant Mouse. J. Biol. Chem. 276, 35429–35434. 10.1074/jbc.M100379200.

36. Pallerla, S.R., Lawrence, R., Lewejohann, L., Pan, Y., Fischer, T., Schlomann, U., Zhang, X., Esko, J.D., and Grobe, K. (2008). Altered Heparan Sulfate Structure in Mice with Deleted NDST3 Gene Function. J. Biol. Chem. 283, 16885–16894. 10.1074/jbc.M709774200.

37. Schlessinger, J., Plotnikov, A.N., Ibrahimi, O.A., Eliseenkova, A.V., Yeh, B.K., Yayon, A., Linhardt, R.J., and Mohammadi, M. (2000). Crystal Structure of a Ternary FGF-FGFR-Heparin Complex Reveals a Dual Role for Heparin in FGFR Binding and Dimerization. Mol. Cell 6, 743–750. 10.1016/S1097-2765(00)00073-3.

38. Jin, L., Abrahams, J.P., Skinner, R., Petitou, M., Pike, R.N., and Carrell, R.W. (1997). The anticoagulant activation of antithrombin by heparin. Proc. Natl. Acad. Sci. 94, 14683–14688. 10.1073/pnas.94.26.14683.

39. Nagy, G.N., Zhao, X.-F., Karlsson, R., Wang, K., Duman, R., Harlos, K., El Omari, K., Wagner, A., Clausen, H., Miller, R.L., et al. (2024). Structure and function of Semaphorin-5A glycosaminoglycan interactions. Nat. Commun. 15, 2723. 10.1038/s41467-024-46725-7.

40. Coles, C.H., Shen, Y., Tenney, A.P., Siebold, C., Sutton, G.C., Lu, W., Gallagher, J.T., Jones, E.Y., Flanagan, J.G., and Aricescu, A.R. (2011). Proteoglycan-Specific Molecular Switch for RPTP Clustering and Neuronal Extension. Science 332, 484–488. 10.1126/science.1200840.

41. Li, Z., and Feizi, T. (2018). The neoglycolipid (NGL) technology-based microarrays and future prospects. FEBS Lett. 592, 3976–3991. 10.1002/1873-3468.13217.

42. Liu, L., Chopra, P., Li, X., Bouwman, K.M., Tompkins, S.M., Wolfert, M.A., de Vries, R.P., and Boons, G.-J. (2021). Heparan Sulfate Proteoglycans as Attachment Factor for SARS-CoV-2. ACS Cent. Sci. 7, 1009–1018. 10.1021/acscentsci.1c00010.

43. Scott, R.P., and Ibáñez, C.F. (2001). Determinants of ligand binding specificity in the glial cell line-derived neurotrophic factor family receptor αs. J. Biol. Chem. 276, 1450–1458. 10.1074/jbc.M006157200.

44. Punjani, A., Rubinstein, J.L., Fleet, D.J., and Brubaker, M.A. (2017). cryoSPARC: algorithms for rapid unsupervised cryo-EM structure determination. Nat. Methods 14, 290–296. 10.1038/nmeth.4169.

45. Zivanov, J., Nakane, T., and Scheres, S.H.W. (2019). A Bayesian approach to beam-induced motion correction in cryo-EM single-particle analysis. IUCrJ 6, 5–17. 10.1107/S205225251801463X.

46. Liebschner, D., Afonine, P.V., Baker, M.L., Bunkóczi, G., Chen, V.B., Croll, T.I., Hintze, B., Hung, L.-W., Jain, S., McCoy, A.J., et al. (2019). Macromolecular structure determination using X-rays, neutrons and electrons: recent developments in *Phenix*. Acta Crystallogr. Sect. Struct. Biol. 75, 861–877. 10.1107/S2059798319011471.

47. Mulloy, B., Forster, M.J., Jones, C., and Davies, D.B. (1993). N.m.r. and molecular-modelling studies of the solution conformation of heparin. Biochem. J. 293, 849–858. 10.1042/bj2930849.

48. Virtanen, H., Yang, J., Bespalov, M.M., Hiltunen, J.O., Leppänen, V.-M., Kalkkinen, N., Goldman, A., Saarma, M., and Runeberg-Roos, P. (2005). The first cysteine-rich domain of the receptor GFRα1 stabilizes the binding of GDNF. Biochem. J. 387, 817–824. 10.1042/BJ20041257.

49. Eketjall, S. (1999). Distinct structural elements in GDNF mediate binding to GFRalpha 1 and activation of the GFRalpha 1-c-Ret receptor complex. EMBO J. 18, 5901–5910. 10.1093/emboj/18.21.5901.

50. Sandmark, J., Dahl, G., Öster, L., Xu, B., Johansson, P., Akerud, T., Aagaard, A., Davidsson, P., Bigalke, J.M., Winzell, M.S., et al. (2018). Structure and biophysical characterization of the human full-length neurturin-GFRa2 complex: A role for heparan sulfate in signaling. J. Biol. Chem. 293, 5492–5508. 10.1074/jbc.RA117.000820.

51. Horibata, S., Rice, E.J., Mukai, C., Marks, B.A., Sams, K., Zheng, H., Anguish, L.J., Coonrod, S.A., and Danko, C.G. (2018). ER-positive breast cancer cells are poised for RET-mediated endocrine resistance. PLOS ONE 13, e0194023. 10.1371/journal.pone.0194023.

52. Marks, B.A., Pipia, I.M., Mukai, C., Horibata, S., Rice, E.J., Danko, C.G., and Coonrod, S.A. (2023). GDNF-RET signaling and EGR1 form a positive feedback loop that promotes tamoxifen resistance via cyclin D1. BMC Cancer 23, 138. 10.1186/s12885-023-10559-1.

53. Xicoy, H., Wieringa, B., and Martens, G.J.M. (2017). The SH-SY5Y cell line in Parkinson’s disease research: a systematic review. Mol. Neurodegener. 12, 10. 10.1186/s13024-017-0149-0.

54. Yamada, S., Nomura, T., Uebersax, L., Matsumoto, K., Fujita, S., Miyake, M., and Miyake, J. (2007). Retinoic acid induces functional c-Ret tyrosine kinase in human neuroblastoma: NeuroReport 18, 359–363. 10.1097/WNR.0b013e32801299b4.

55. Merkouris, S., Barde, Y.-A., Binley, K.E., Allen, N.D., Stepanov, A.V., Wu, N.C., Grande, G., Lin, C.-W., Li, M., Nan, X., et al. (2018). Fully human agonist antibodies to TrkB using autocrine cell-based selection from a combinatorial antibody library. Proc. Natl. Acad. Sci. 115. 10.1073/pnas.1806660115.

56. Encinas, M., Iglesias, M., Liu, Y., Wang, H., Muhaisen, A., Ceña, V., Gallego, C., and Comella, J.X. (2000). Sequential treatment of SH-SY5Y cells with retinoic acid and brain-derived neurotrophic factor gives rise to fully differentiated, neurotrophic factor-dependent, human neuron-like cells. J. Neurochem. 75, 991–1003. 10.1046/j.1471-4159.2000.0750991.x.

57. Moccia, M., Yang, D., Lakkaniga, N.R., Frett, B., McConnell, N., Zhang, L., Brescia, A., Federico, G., Zhang, L., Salerno, P., et al. (2021). Targeted activity of the small molecule kinase inhibitor Pz-1 towards RET and TRK kinases. Sci. Rep. 11, 16103. 10.1038/s41598-021-95612-4.

58. Bespalov, M.M., Sidorova, Y.A., Tumova, S., Ahonen-Bishopp, A., Magalhães, A.C., Kulesskiy, E., Paveliev, M., Rivera, C., Rauvala, H., and Saarma, M. (2011). Heparan sulfate proteoglycan syndecan-3 is a novel receptor for GDNF, neurturin, and artemin. J. Cell Biol. 192, 153–169. 10.1083/jcb.201009136.

59. Häcker, U., Nybakken, K., and Perrimon, N. (2005). Heparan sulphate proteoglycans: the sweet side of development. Nat. Rev. Mol. Cell Biol. 6, 530–541. 10.1038/nrm1681.

60. McKeithan, T.W. (1995). Kinetic proofreading in T-cell receptor signal transduction. Proc. Natl. Acad. Sci. 92, 5042–5046. 10.1073/pnas.92.11.5042.

61. Patel, A., Harker, N., Moreira-Santos, L., Ferreira, M., Alden, K., Timmis, J., Foster, K., Garefalaki, A., Pachnis, P., Andrews, P., et al. (2012). Differential RET signaling pathways drive development of the enteric lymphoid and nervous systems. Sci. Signal. 5, 1–12. 10.1126/scisignal.2002734.

62. Enomoto, H., Hughes, I., Golden, J., Baloh, R.H., Yonemura, S., Heuckeroth, R.O., Johnson, E.M., and Milbrandt, J. (2004). GFRα1 Expression in Cells Lacking RET Is Dispensable for Organogenesis and Nerve Regeneration. Neuron 44, 623–636. 10.1016/j.neuron.2004.10.032.

63. Barnett, M.W., Fisher, C.E., Perona-Wright, G., and Davies, J.A. (2002). Signalling by glial cell line-derived neurotrophic factor (GDNF) requires heparan sulphate glycosaminoglycan. J. Cell Sci. 115, 4495–4503. 10.1242/jcs.00114.

64. Pratt, T., Conway, C.D., Tian, N.M.M.-L., Price, D.J., and Mason, J.O. (2006). Heparan Sulphation Patterns Generated by Specific Heparan Sulfotransferase Enzymes Direct Distinct Aspects of Retinal Axon Guidance at the Optic Chiasm. J. Neurosci. 26, 6911–6923. 10.1523/JNEUROSCI.0505-06.2006.

65. Sugaya, N., Habuchi, H., Nagai, N., Ashikari-Hada, S., and Kimata, K. (2008). 6-O-Sulfation of Heparan Sulfate Differentially Regulates Various Fibroblast Growth Factor-dependent Signalings in Culture. J. Biol. Chem. 283, 10366–10376. 10.1074/jbc.M705948200.

66. Sjöstrand, D., and Ibáñez, C.F. (2008). Insights into GFRα1 regulation of neural cell adhesion molecule (NCAM) function from structure-function analysis of the NCAM/GFRα1 receptor complex. J. Biol. Chem. 283, 13792–13798. 10.1074/jbc.M800283200.

67. Gude, F., Froese, J., Manikowski, D., Di Iorio, D., Grad, J.-N., Wegner, S., Hoffmann, D., Kennedy, M., Richter, R.P., Steffes, G., et al. (2023). Hedgehog is relayed through dynamic heparan sulfate interactions to shape its gradient. Nat. Commun. 14, 758. semaph.

68. Runeberg-Roos, P., Piccinini, E., Penttinen, A.-M., Mätlik, K., Heikkinen, H., Kuure, S., Bespalov, M.M., Peränen, J., Garea-Rodríguez, E., Fuchs, E., et al. (2016). Developing therapeutically more efficient Neurturin variants for treatment of Parkinson’s disease. Neurobiol. Dis. 96, 335–345. 10.1016/j.nbd.2016.07.008.

69. Salvatore, M., Ai, Y., Fischer, B., Zhang, A., Grondin, R., Zhang, Z., Gerhardt, G., and Gash, D. (2006). Point source concentration of GDNF may explain failure of phase II clinical trial. Exp. Neurol. 202, 497–505. 10.1016/j.expneurol.2006.07.015.

70. Liu, Y., De Castro Ribeiro, O., Haapanen, O., Craven, G.B., Sharma, V., Muench, S.P., and Goldman, A. (2022). Unexpected structures formed by the kinase RET C634R mutant extracellular domain suggest potential oncogenic mechanisms in MEN2A. J. Biol. Chem. 298, 102380. 10.1016/j.jbc.2022.102380.

71. Chandra, N., Liu, Y., Liu, J.-X., Frängsmyr, L., Wu, N., Silva, L., Lindström, M., Chai, W., Pedrosa Domellöf, F., Feizi, T., et al. (2019). Sulfated Glycosaminoglycans as Viral Decoy Receptors for Human Adenovirus Type 37. Viruses 11, 247. 10.3390/v11030247.

72. Silva, J.C., Denny, R., Dorschel, C.A., Gorenstein, M., Kass, I.J., Li, G.-Z., McKenna, T., Nold, M.J., Richardson, K., Young, P., et al. (2005). Quantitative Proteomic Analysis by Accurate Mass Retention Time Pairs. Anal. Chem. 77, 2187–2200. 10.1021/ac048455k.

73. Richter, R., Mukhopadhyay, A., and Brisson, A. (2003). Pathways of Lipid Vesicle Deposition on Solid Surfaces: A Combined QCM-D and AFM Study. Biophys. J. 85, 3035–3047. 10.1016/S0006-3495(03)74722-5.

74. Srimasorn, S., Souter, L., Green, D.E., Djerbal, L., Goodenough, A., Duncan, J.A., Roberts, A.R.E., Zhang, X., Débarre, D., DeAngelis, P.L., et al. (2022). A quartz crystal microbalance method to quantify the size of hyaluronan and other glycosaminoglycans on surfaces. Sci. Rep. 12, 10980. 10.1038/s41598-022-14948-7.

75. Thakar, D., Migliorini, E., Coche-Guerente, L., Sadir, R., Lortat-Jacob, H., Boturyn, D., Renaudet, O., Labbe, P., and Richter, R.P. (2014). A quartz crystal microbalance method to study the terminal functionalization of glycosaminoglycans. Chem Commun 50, 15148–15151. 10.1039/C4CC06905F.

76. Taylor, S.L., Hogwood, J., Guo, W., Yates, E.A., and Turnbull, J.E. (2019). By-Products of Heparin Production Provide a Diverse Source of Heparin-like and Heparan Sulfate Glycosaminoglycans. Sci. Rep. 9, 2679. 10.1038/s41598-019-39093-6.

77. Reviakine, I., Johannsmann, D., and Richter, R.P. (2011). Hearing What You Cannot See and Visualizing What You Hear: Interpreting Quartz Crystal Microbalance Data from Solvated Interfaces. Anal. Chem. 83, 8838–8848. 10.1021/ac201778h.

78. Dubacheva, G.V., Araya-Callis, C., Geert Volbeda, A., Fairhead, M., Codée, J., Howarth, M., and Richter, R.P. (2017). Controlling Multivalent Binding through Surface Chemistry: Model Study on Streptavidin. J. Am. Chem. Soc. 139, 4157–4167. 10.1021/jacs.7b00540.

79. Migliorini, E., Thakar, D., Sadir, R., Pleiner, T., Baleux, F., Lortat-Jacob, H., Coche-Guerente, L., and Richter, R.P. (2014). Well-defined biomimetic surfaces to characterize glycosaminoglycan-mediated interactions on the molecular, supramolecular and cellular levels. Biomaterials 35, 8903–8915. 10.1016/j.biomaterials.2014.07.017.

