## Supplementary Information for "A long-chain heparan sulfate capture mechanism directs paracrine GDNF-GFRα1 signalling through RET"

### Supplementary Figures

**A.**

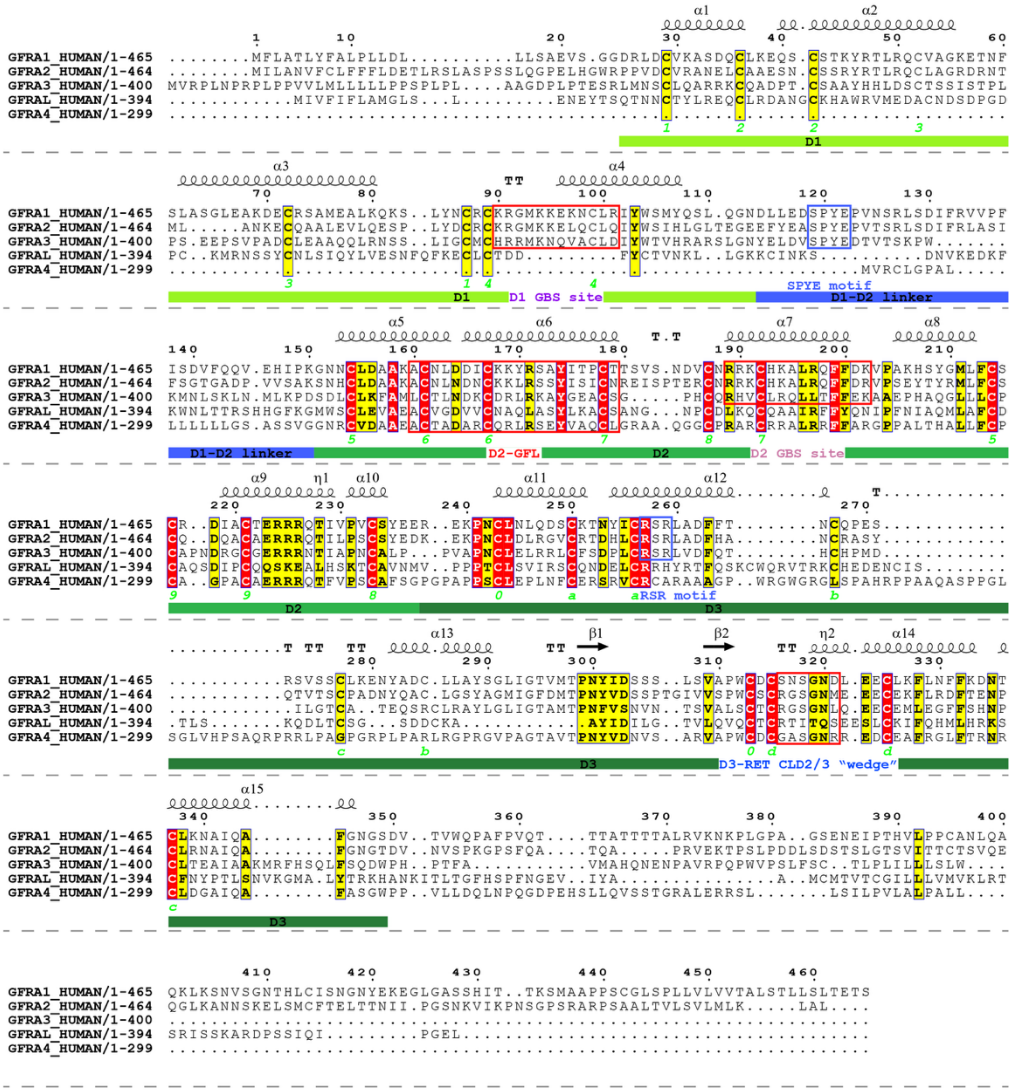


**B.**

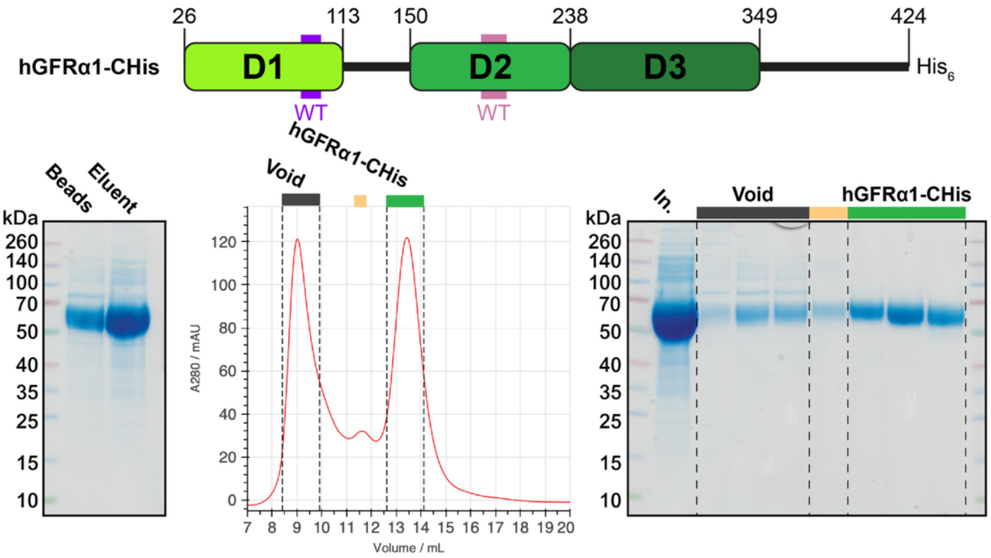

**Figure S1: (A)** Annotated sequence alignment of the five RET co-receptors. Generated using ESPript[1] (https://espript.ibcp.fr). **(B)** Domain diagram, SDS-PAGE analysis of Ni-NTA extraction and SEC elution profile monitored by absorbance at 280 nm and SDS-PAGE for hGFRα1-CHis.

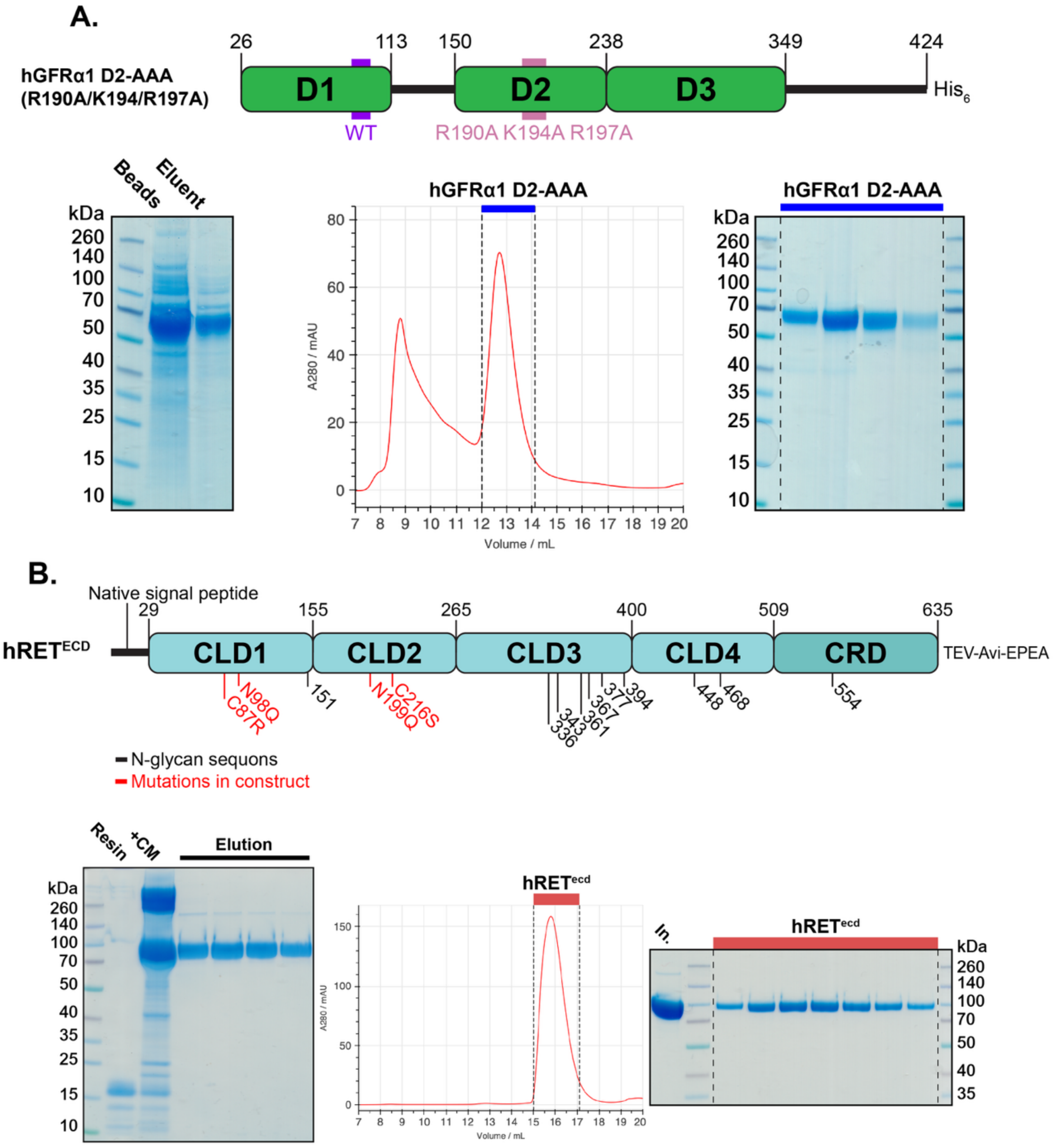


**Figure S2: (A)** Domain diagram, SDS-PAGE analysis of Ni-NTA extraction and SEC elution profile monitored by absorbance at 280 nm and SDS-PAGE for hGFRα1-CHis D2-AAA. **(B)** Domain diagram, SDS-PAGE analysis of C-Tag resin extraction and SEC elution profile monitored by absorbance at 280 nm and SDS-PAGE for hRET$^{ECD}$.

**A.**

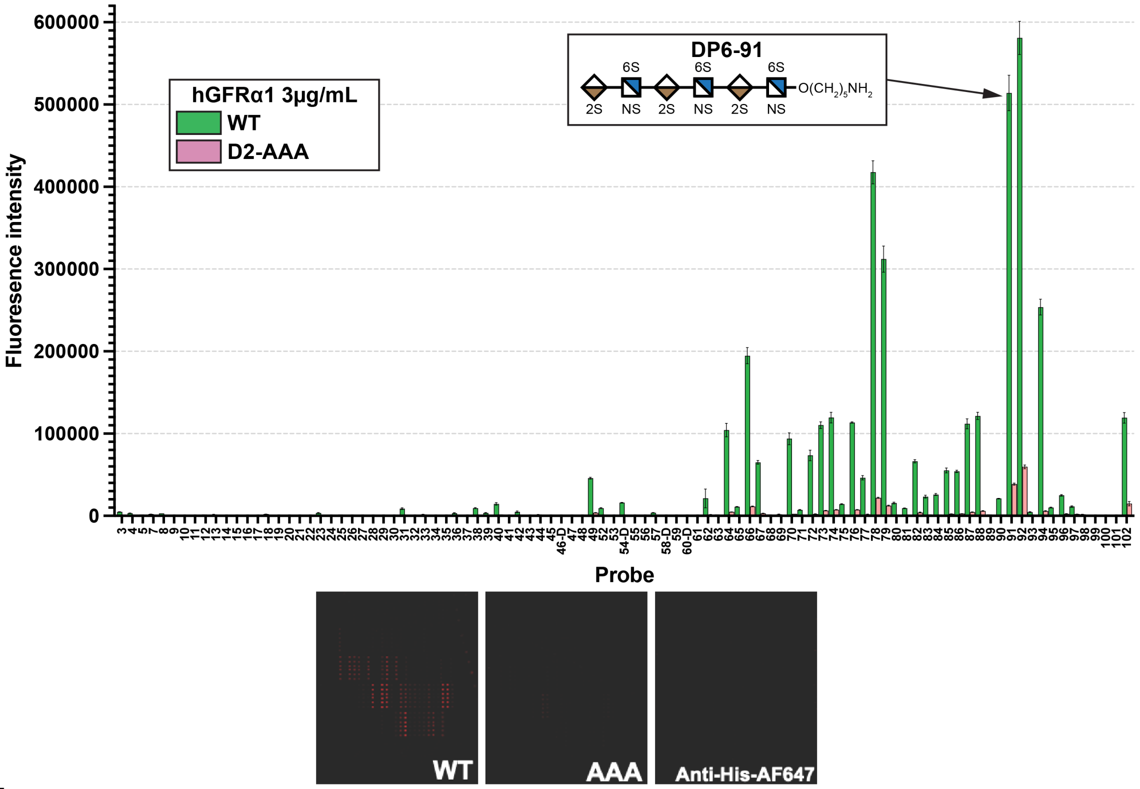


**B.**

| Array# | HS | sulfates | structure |
|---|---|---|---|
| 3 | tetra | 5 | GlcA-GlcNS6S-IdoA2S-GlcNS6S |
| 4 | tetra | 4 | IdoA-GlcNS6S-GlcA-GlcNS6S |
| 5 | tetra | 3 | GlcA-GlcNAc6S-IdoA2S-GlcNAc6S |
| 7 | tetra | 2 | IdoA-GlcNAc6S-IdoA-GlcNAc6S |
| 8 | tetra | 4 | GlcA-GlcNS6S-IdoA-GlcNS6S |
| 9 | tetra | 2 | IdoA-GlcNAc6S-GlcA-GlcNAc6S |
| 10 | tetra | 2 | GlcA-GlcNAc6S-IdoA-GlcNAc6S |
| 11 | tetra | 3 | IdoA-GlcNAc6S-IdoA2S-GlcNAc6S |
| 12 | tetra | 4 | GlcA-GlcNS6S-GlcA-GlcNS6S |
| 13 | tetra | 4 | IdoA-GlcNS6S-IdoA-GlcNS6S |
| 14 | tetra | 2 | GlcA-GlcNAc6S-GlcA-GlcNAc6S |
| 15 | tetra | 1 | GlcA-GlcNAc-IdoA2S-GlcNAc |
| 16 | tetra | 3 | GlcA-GlcNS-IdoA2S-GlcNS |
| 17 | tetra | 2 | GlcA-GlcNAc-GlcA2S-GlcNAc6S |
| 18 | tetra | 4 | GlcA-GlcNS-GlcA2S-GlcNS6S |
| 19 | tetra | 1 | GlcA-GlcNAc-GlcA2S-GlcNAc |
| 20 | tetra | 3 | GlcA-GlcNS-GlcA2S-GlcNS |
| 21 | tetra | 0 | GlcA-GlcNAc-GlcA-GlcNAc |
| 22 | tetra | 2 | GlcA-GlcNAc-IdoA-GlcNS6S |
| 23 | tetra | 4 | GlcA-GlcNS-IdoA2S-GlcNS6S |
| 24 | tetra | 2 | GlcA-GlcNAc-IdoA2S-GlcNAc6S |
| 25 | tetra | 3 | GlcA-GlcNAc-IdoA2S-GlcNS6S |
| 26 | tetra | 1 | GlcA-GlcNAc-IdoA-GlcNAc6S |
| 27 | tetra | 3 | GlcA-GlcNS-IdoA-GlcNS6S |
| 28 | tetra | 0 | GlcA-GlcNAc-IdoA-GlcNAc |
| 29 | di | 1 | IdoA-GlcNAc6S |
| 30 | tetra | 1 | IdoA-GlcNAc6S-GlcA-GlcNAc |
| 31 | tetra | 5 | IdoA2S-GlcNS6S-GlcA-GlcNS6S |
| 32 | tetra | 3 | IdoA-GlcNS6S-GlcA-GlcNS |
| 33 | tetra | 3 | IdoA2S-GlcNAc6S-GlcA-GlcNAc6S |
| 34 | tetra | 2 | IdoA2S-GlcNAc6S-GlcA-GlcNAc |
| 35 | tetra | 1 | IdoA2S-GlcNAc-GlcA-GlcNAc |
| 36 | tetra | 4 | IdoA2S-GlcNS6S-GlcA-GlcNS |
| 37 | tetra | 3 | IdoA2S-GlcNS-GlcA-GlcNS |
| 38 | tetra | 5 | IdoA-GlcNS6S-IdoA2S-GlcNS6S |
| 39 | tetra | 4 | IdoA2S-GlcNAc6S-IdoA2S-GlcNAc6S |
| 40 | tetra | 6 | IdoA2S-GlcNS6S-IdoA2S-GlcNS6S |
| 41 | hexa | 2 | GlcA-GlcNAc-IdoA2S-GlcNAc6S-GlcA-GlcNAc |
| 42 | hexa | 5 | GlcA-GlcNS-IdoA2S-GlcNS6S-GlcA-GlcNS |
| 43 | di | 2 | IdoA2S-GlcNAc6S |
| 44 | di | 3 | IdoA2S-GlcNS6S |
| 45 | di | 1 | GlcA-GlcNAc6S |
| 46 | tetra-CS | 2 | GlcA-GlcNS6S-GlcA-GlcNAc |
| 47 | tetra | 1 | IdoA-GlcNAc-IdoA-GlcNAc6S |
| 48 | tetra | 1 | IdoA-GlcNAc6S-IdoA-GlcNAc |
| 49 | tetra | 6 | GlcA-GlcNS3S6S-IdoA2S-GlcNS6S |

| Array# | HS | sulfates | structure |
|---|---|---|---|
| 52 | tetra | 5 | GlcA-GlcNS3S-IdoA2S-GlcNS6S |
| 53 | tetra-CS | 1 | GlcA-GalNAc-GlcA-GalNAc4S |
| 54 | octa | 7 | GlcA-GlcNS6S-IdoA-GlcNS-IdoA2S-GlcNS6S-IdoA-GlcNAc6S |
| 55 | tetra | 2 | GlcA-GlcNS6S-GlcA-GlcNAc |
| 56 | tetra | 2 | IdoA-GlcNS-IdoA-GlcNS |
| 57 | tetra | 4 | IdoA-GlcNS-IdoA2S-GlcNS6S |
| 58 | tetra | 3 | GlcA-GlcNS6S-IdoA-GlcNS |
| 59 | tetra | 4 | IdoA2S-GlcNS6S-IdoA-GlcNAc6S |
| 60 | tetra | 3 | IdoA-GlcNS-IdoA-GlcNS6S |
| 62 | Hexa | 7 | GlcA-GlcNS6S-GlcA-GlcNS3S6S-GlcA-GlcNS6S |
| 63 | Hexa | 6 | GlcA-GlcNS6S-GlcA-GlcNS6S-GlcA-GlcNS6S |
| 64 | Hexa | 7 | GlcA-GlcNS6S-IdoA-GlcNS3S6S-GlcA-GlcNS6S |
| 65 | Hexa | 6 | GlcA-GlcNS6S-IdoA-GlcNS6S-GlcA-GlcNS6S |
| 66 | Hexa | 8 | GlcA-GlcNS6S-IdoA2S-GlcNS3S6S-GlcA-GlcNS6S |
| 67 | Hexa | 7 | GlcA-GlcNS6S-IdoA2S-GlcNS6S-GlcA-GlcNS6S |
| 68 | Hexa | 7 | GlcA-GlcNS6S-GlcA-GlcNS3S6S-IdoA-GlcNS6S |
| 69 | Hexa | 6 | GlcA-GlcNS6S-GlcA-GlcNS6S-IdoA-GlcNS6S |
| 70 | Hexa | 7 | GlcA-GlcNS6S-IdoA-GlcNS3S6S-IdoA-GlcNS6S |
| 71 | Hexa | 6 | GlcA-GlcNS6S-IdoA-GlcNS6S-IdoA-GlcNS6S |
| 72 | Hexa | 8 | GlcA-GlcNS6S-IdoA2S-GlcNS3S6S-IdoA-GlcNS6S |
| 73 | Hexa | 7 | GlcA-GlcNS6S-IdoA2S-GlcNS6S-IdoA-GlcNS6S |
| 74 | Hexa | 8 | GlcA-GlcNS6S-GlcA-GlcNS3S6S-IdoA2S-GlcNS6S |
| 75 | Hexa | 7 | GlcA-GlcNS6S-GlcA-GlcNS6S-IdoA2S-GlcNS6S |
| 76 | Hexa | 8 | GlcA-GlcNS6S-IdoA-GlcNS3S6S-IdoA2S-GlcNS6S |
| 77 | Hexa | 7 | GlcA-GlcNS6S-IdoA-GlcNS6S-IdoA2S-GlcNS6S |
| 78 | Hexa | 9 | GlcA-GlcNS6S-IdoA2S-GlcNS3S6S-IdoA2S-GlcNS6S |
| 79 | Hexa | 8 | GlcA-GlcNS6S-IdoA2S-GlcNS6S-IdoA2S-GlcNS6S |
| 80 | Hexa | 6 | GlcA-GlcNS6S-GlcA-GlcNS3S-GlcA-GlcNS6S |
| 81 | Hexa | 6 | GlcA-GlcNS6S-IdoA-GlcNS3S-GlcA-GlcNS6S |
| 82 | Hexa | 7 | GlcA-GlcNS6S-IdoA2S-GlcNS3S-GlcA-GlcNS6S |
| 83 | Hexa | 6 | GlcA-GlcNS6S-GlcA-GlcNS3S-IdoA-GlcNS6S |
| 84 | Hexa | 6 | GlcA-GlcNS6S-IdoA-GlcNS3S-IdoA-GlcNS6S |
| 85 | Hexa | 7 | GlcA-GlcNS6S-IdoA2S-GlcNS3S-IdoA-GlcNS6S |
| 86 | Hexa | 6 | GlcA-GlcNS6S-GlcA-GlcNS3S-IdoA2S-GlcNS6S |
| 87 | Hexa | 7 | GlcA-GlcNS6S-IdoA-GlcNS3S-IdoA2S-GlcNS6S |
| 88 | Hexa | 8 | GlcA-GlcNS6S-IdoA2S-GlcNS3S-IdoA2S-GlcNS6S |
| 89 | Di | 3 | IdoA2S-GlcNS6S |
| 90 | Tetra | 6 | IdoA2S-GlcNS6S-IdoA2S-GlcNS6S |
| 91 | Hexa | 9 | IdoA2S-GlcNS6S-IdoA2S-GlcNS6S-IdoA2S-GlcNS6S |
| 92 | Octa | 12 | IdoA2S-GlcNS6S-IdoA2S-GlcNS6S-IdoA2S-GlcNS6S-IdoA2S- |
| 93 | Hexa | 5 | GlcA-GlcNAc-IdoA2S-GlcNS6S-IdoA2S-GlcNS |
| 94 | Hexa | 7 | GlcA-GlcNS6S-IdoA2S-GlcNS6S-IdoA2S-GlcNS |
| 95 | Hexa | 6 | GlcA-GlcNS-IdoA2S-GlcNS6S-IdoA2S-GlcNS |
| 96 | Hexa | 6 | GlcA-GlcNAc6S-IdoA2S-GlcNS6S-IdoA2S-GlcNS |
| 97 | Tetra | 5 | IdoA2S-GlcNS6S-IdoA2S-GlcNS |
| 98 | Tetra | 5 | GlcA-GlcNS6S-GlcA2S-GlcNS6S |
| 99 | Tetra | 3 | GlcA-GlcNAc6S-GlcA2S-GlcNAc6S |
| 100 | Tetra | 1 | IdoA-GlcNS-IdoA-GlcNAc |
| 101 | Tetra | 1 | IdoA-GlcNAc-IdoA-GlcNS |

**Figure S3: (A)** Histogram of fluorescence intensities and image of sequence-defined HS arrays assayed with hGFRα1-CHis WT/D2-AAA at 3µg/mL. **(B)** Table of structures of the probes printed on the sequence-defined HS array.

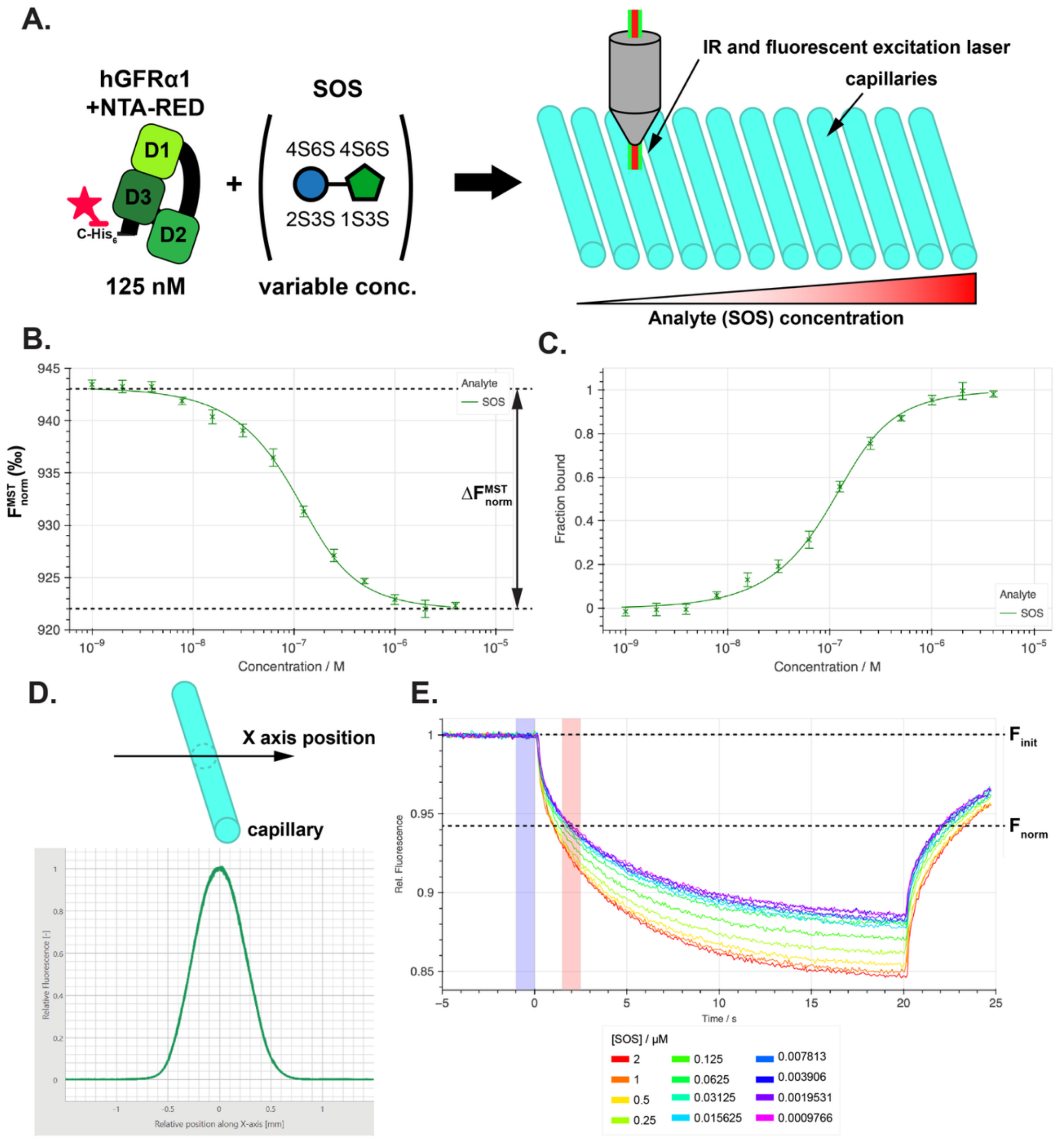


**Figure S4: (A)** Schematic detailing experimental setup for measurement of hGFRα1-CHis-glycan interaction by MST. **(B)** hGFRα1-CHis/SOS binding curve measured using MST. $\Delta F_{norm}$ is calculated from the best-fit binding model to the data. MST of the full SOS concentration series was performed in triplicate. **(C)** The same data plotted in terms of fraction bound (0-1). **(D)** Measurement of fluorescence profile transverse to the capillaries indicates no sticking of labelled protein to glass. **(E)** Raw MST traces with cold (blue) and hot (red) regions used for deriving binding curve data.

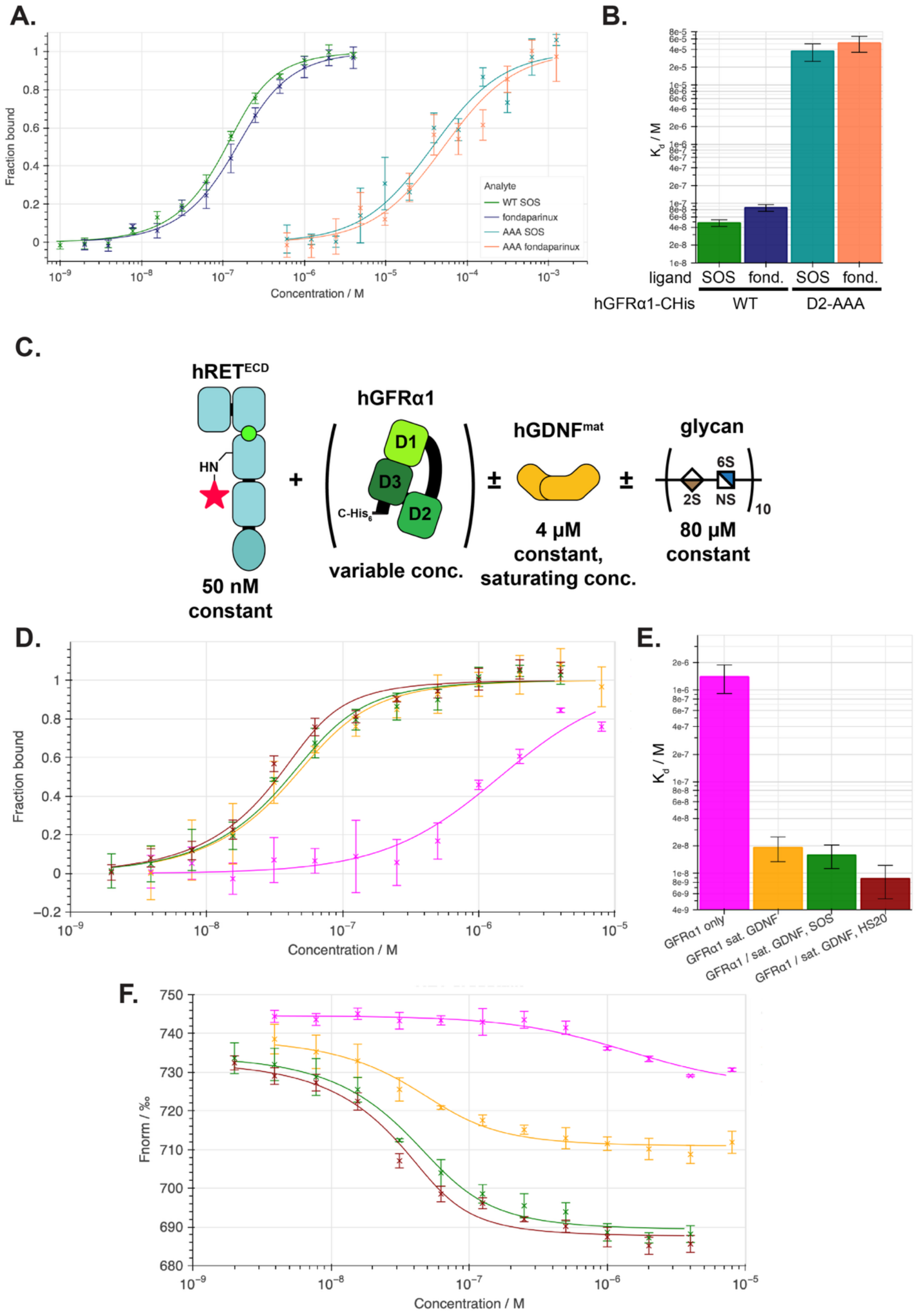


**Figure S5: (A/B)** Binding curves and calculated $K_d$ values for hGFRα1-CHis WT/D2-AAA t to SOS and fondaparinux measured by MST. Binding data points are plotted ±SD as the mean for each concentration, normalised to the fraction bound of the binding model. Derived $K_d$ values for each ligand are plotted with ±the standard error of the least-squares fit of the 1:1 binding model to the data. **(C)** Schematic detailing experimental setup for measuring the $K_d$ of formation of the hRET$^{ECD*}$ ternary complex by MST. **(D-F)** Binding curves (fraction bound, $F^{MST}{}_{norm}$) and derived $K_d$ measured by MST for hRET$^{ECD*}$ to hGFRα1-CHis with and without of hGDNF$^{mat}$ and glycans.

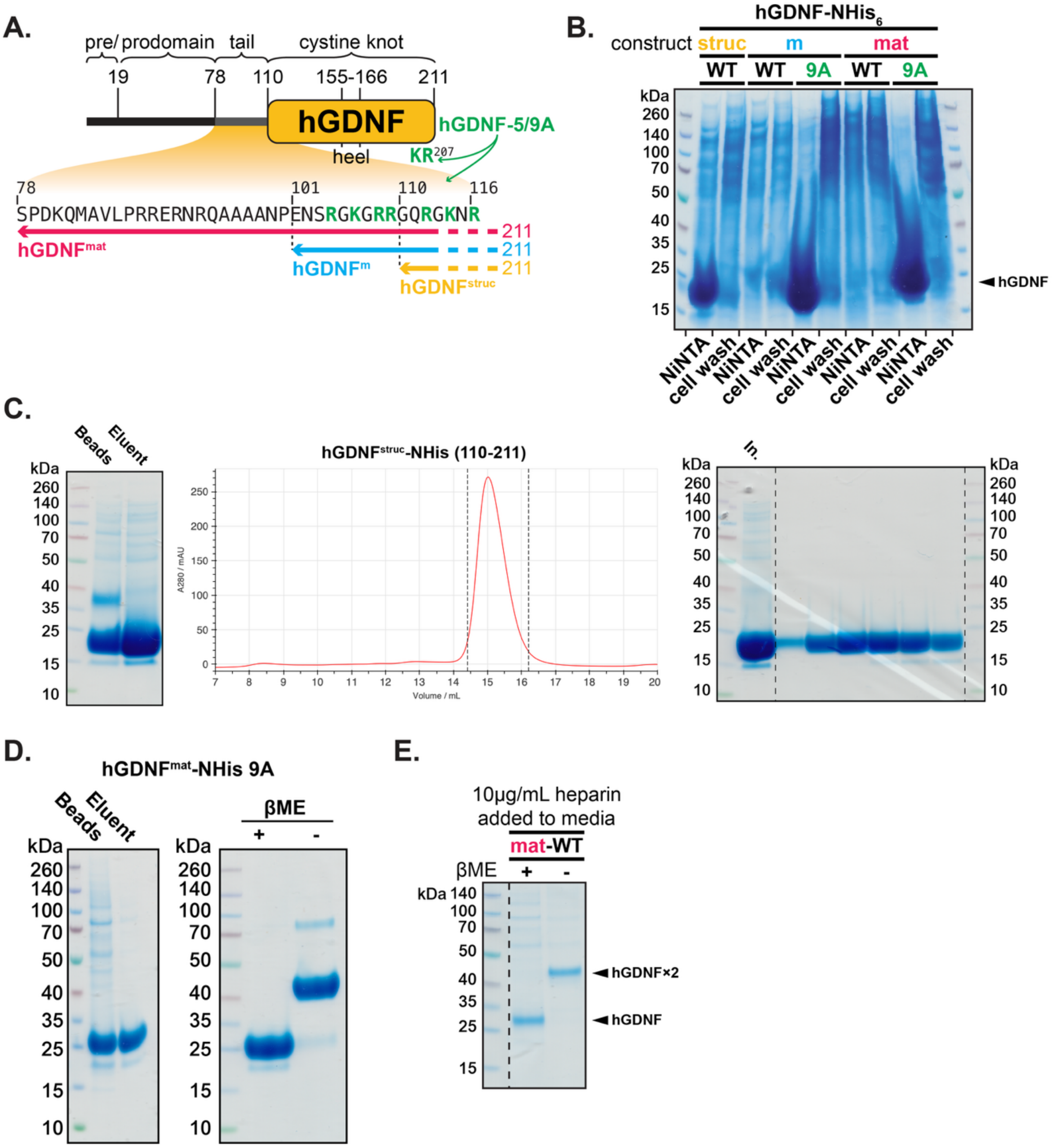


**Figure S6: (A)** Schematic of hGDNF sequence with the domain boundaries and mutations used for production of the recombinant proteins hGDNF$^{mat}$ WT/9A, hGDNF$^{m}$ and hGDNF$^{struc}$ WT/5A. **(B)** SDS-PAGE analysis of Ni-NTA resin extractions of CM or cell pellet washed with high salt buffer from HEK293 cells transfected with different hGDNF constructs. Samples were boiled with DTT prior to SDS-PAGE. **(C)** SDS-PAGE analysis of Ni-NTA extraction and SEC elution profile monitored by absorbance at 280 nm and SDS-PAGE for hGDNF$^{struc}$. Samples were boiled with DTT prior to SDS-PAGE. **(D)** SDS-PAGE analysis of Ni-NTA extraction for hGDNF$^{mat}$ 9A and SDS-PAGE with and without boiling protein with DTT. **(E)** Expression of hGDNF$^{mat}$-NHis with 10 µg/mL heparin added to cell media 24h after transfection. SDS-PAGE of Ni-NTA extraction from transfected CM.

**A.**

| | | | labelling time | | | |
|---|---|---|---|---|---|---|
| Sequence | Start | End | 3s | 30s | 300s | 3000s |
| ADRLDCVKASDQCLKE | 24 | 39 | -0.1561 | 0.0288 | 0.0078 | 0.0274 |
| LASGLEA | 62 | 68 | -0.0694 | -0.0087 | -0.0677 | 0.093 |
| SLYNCRCKRGMKKEKNCLRIYWSMYQ | 83 | 108 | -0.4291 | -0.226 | 0.4896 | |
| KNCLRIYWSMYQSLQGNDLLED | 97 | 118 | 0.0972 | -0.081 | -0.018 | 0.2408 |
| YQSLQGNDLLEDSPYEPVNSRL | 107 | 128 | -1.2189 | -0.6827 | -0.2238 | 0.0526 |
| LLEDSPYEPVNSRL | 115 | 128 | -1.3032 | -0.8322 | -0.1209 | 0.0399 |
| LEDSPYEPVNSRL | 116 | 128 | -1.342 | -0.8835 | -0.1663 | 0.0171 |
| EDSPYEPVNSRL | 117 | 128 | -1.2105 | -0.8348 | -0.1211 | 0.0403 |
| SDIFRVVPFISDV | 129 | 141 | -0.3819 | -0.0689 | 0.3177 | -0.0929 |
| SPYEPVNSRLSDIFRVVPFISDVFQQVE HIPKGNNCLDAAKACNLDDICKKYRSA | 119 | 174 | -2.3037 | -4.9006 | -4.1883 | |
| KKYRSAY | 168 | 174 | 0.0436 | -0.009 | 0.0566 | 0.0159 |
| SAYITPCT | 172 | 179 | -0.7004 | -0.5837 | -0.2893 | -0.0266 |
| LRQFFDKVPAKHSYGMLFCSCRDIACTE | 196 | 223 | -0.0041 | -0.2634 | 0.299 | -0.2652 |
| NLQDSCKTNYICRSRLADFFTNCQP | 245 | 269 | -0.4856 | -0.5457 | 0.2621 | |
| PESRSVSSCLKENYADCLLAYSGL | 269 | 292 | 0.1866 | 0.2349 | 0.6685 | 0.3272 |
| YADCLLAYSGLIGTVMTPNYIDSSSLS | 282 | 308 | -0.2098 | -0.0776 | -0.3163 | 0.2141 |
| LIGTVMTPNYIDSSSLSVAPWCDCSN | 292 | 317 | -0.3107 | -0.0335 | 0.2926 | |
| SSLSVAPWCDCSNSGNDLEECL | 305 | 326 | -0.0468 | 0.0177 | 0.0265 | -0.0903 |

**B.**

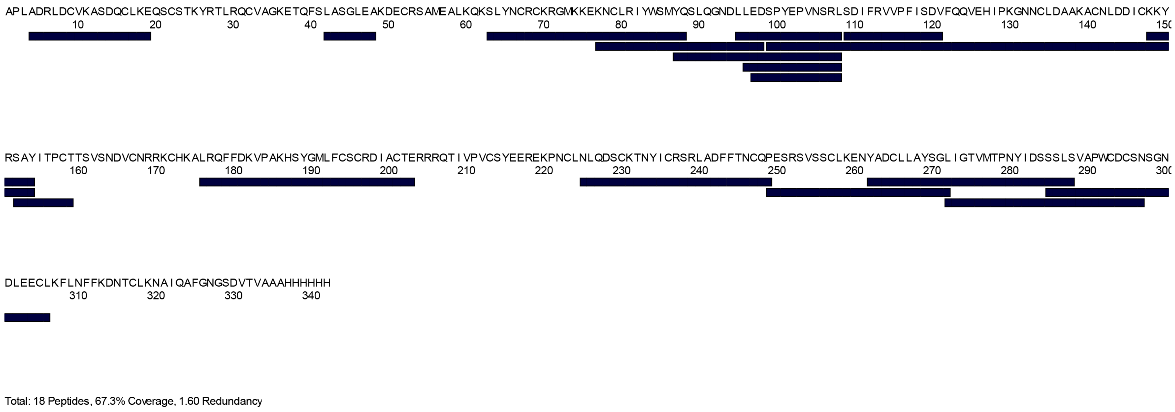


**Figure S7: (A)** Peptides detected by HDX for hGFRα1-CHis N59Q both with and without fondaparinux and the detected mass change for each in Daltons. Mass changes lower than -0.5 are highlighted in blue. Empty cells indicate peptide was not detected. **(B)** Schematic of the above data aligned to the hGFRα1-CHis N59Q sequence.

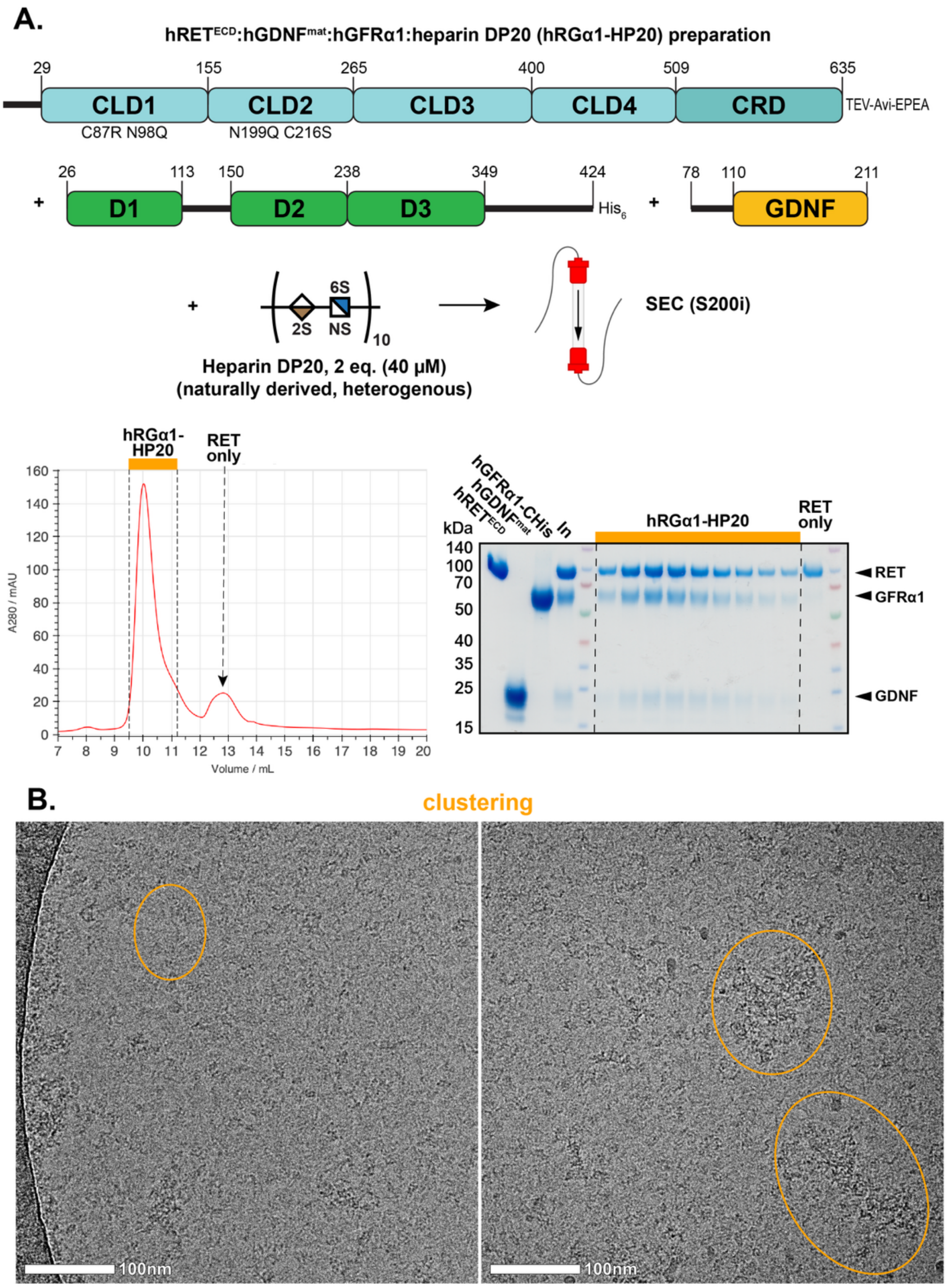


**Figure S8: (A)** Schematic detailing the reconstitution and purification of the hRGα1-HP20 sample and SEC purification of hRGα1-HP20. **(B)** Representative cryo-EM micrographs (200 kV) for RGα1-HP20 grids. Regions of high clustering or aggregation are highlighted. Effective magnification: ×120,000.

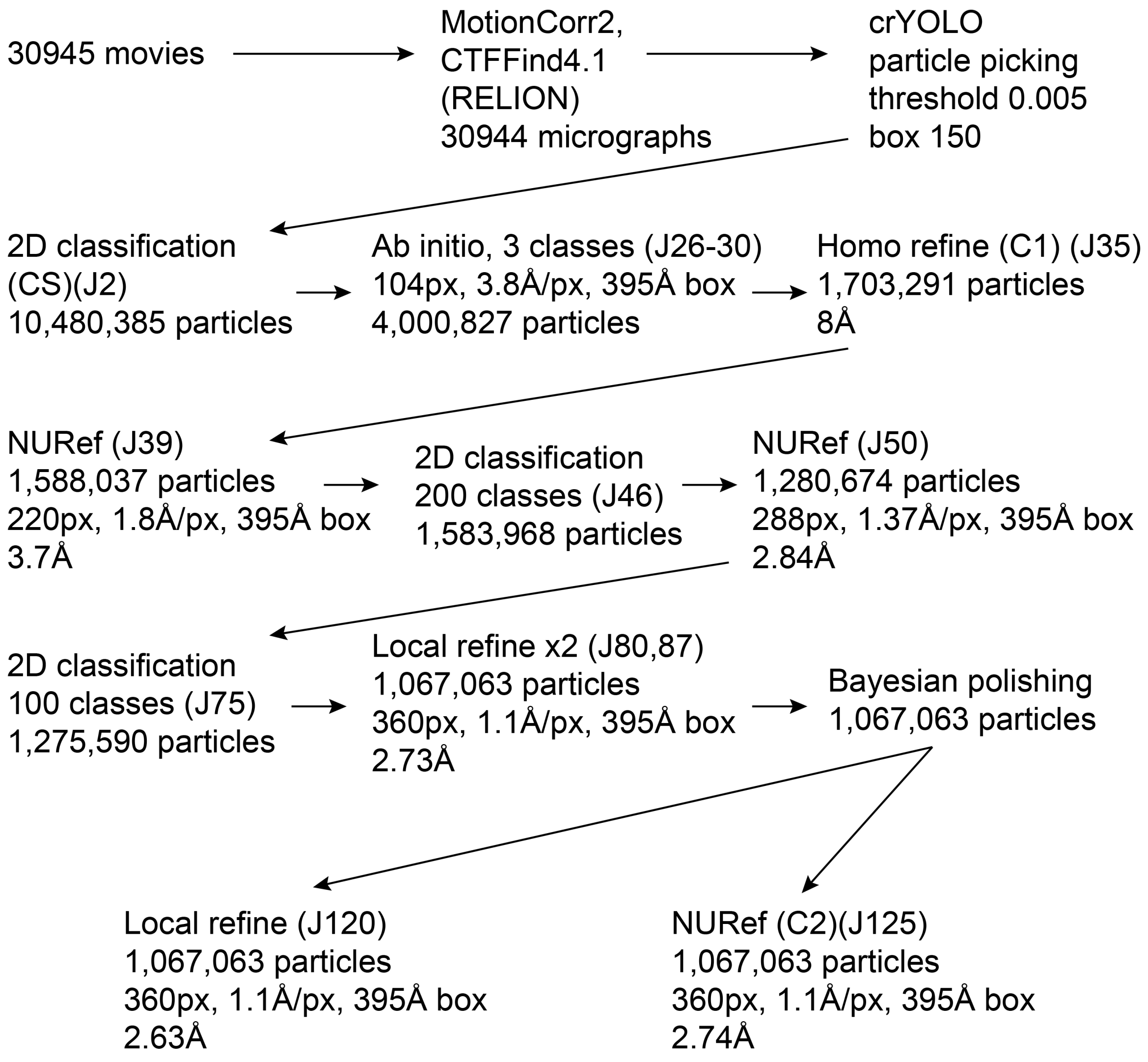


**Figure S9:** Flowchart summarising data processing for the 300kV dataset collected of hRGα1-HP20 grids.

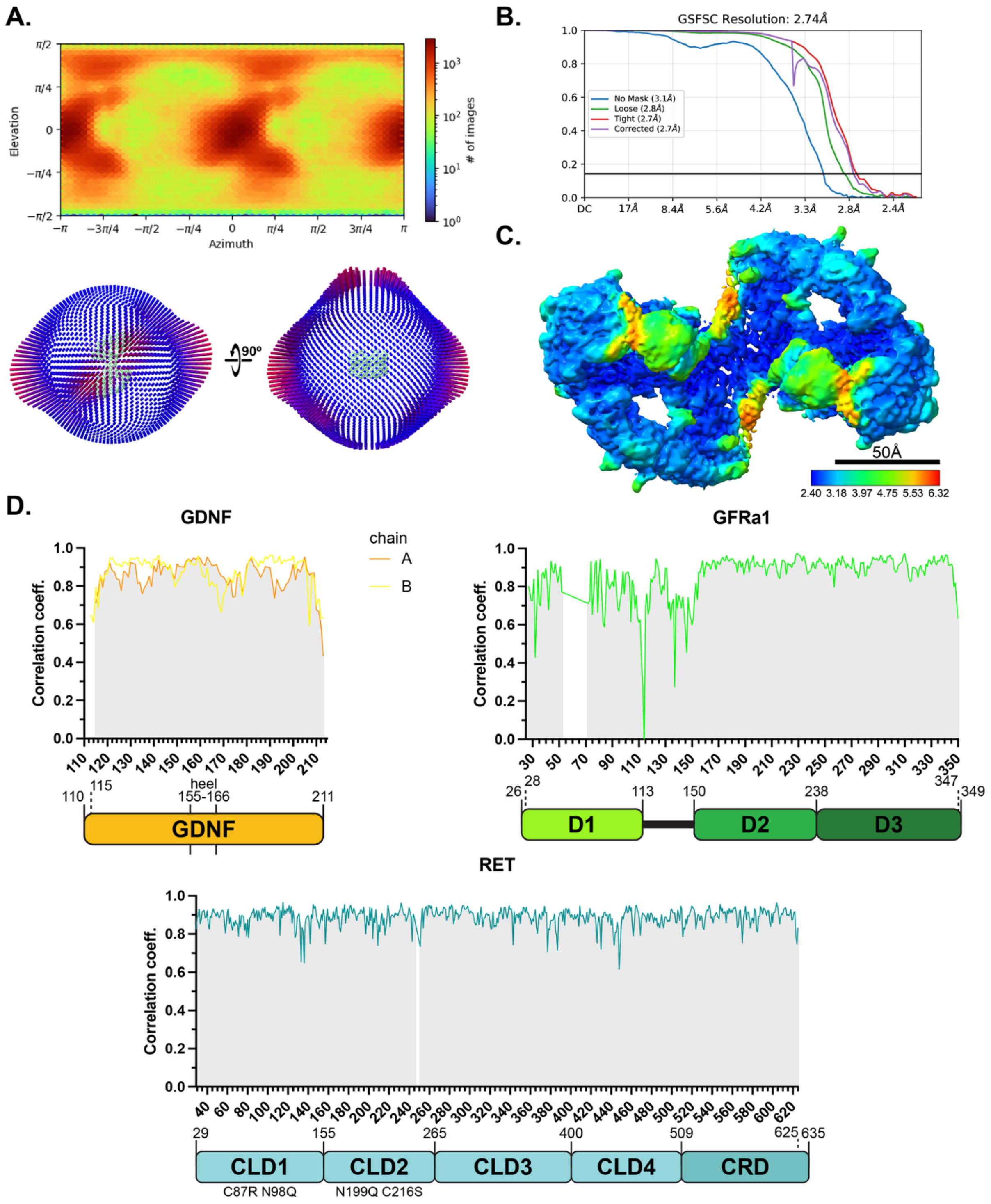


**Figure S10: (A)** Orientation distribution of particles which reconstruct to the final hRGα1-HP20 map, represented as a heatmap and in 3D space. **(B)** Gold-standard Fourier shell correlation (GSFSC) plot for the final hRGα1-HP20 map. **(C)** Surface-colour representation of the computed local resolution of the hRGα1-HP20 map as calculated by CryoSPARC. **(D)** Per-residue map-model correlation coefficient plots for each chain in the hRGα1-HP20 model aligned to domain diagrams. Calculated using Phenix[2].

A.

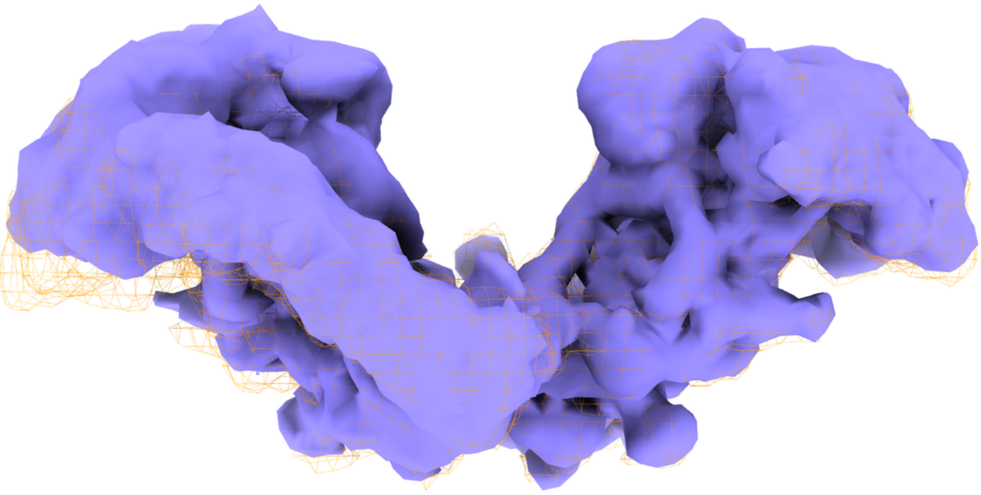

B.

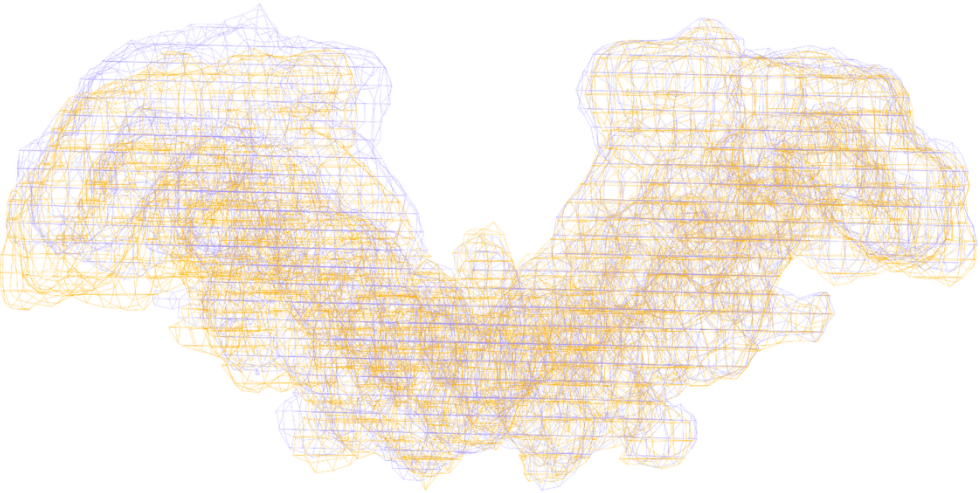

**Figure S11**: 3D classification of hRGα1-HP20 particles (5 classes, 1,588,037 particles) comparing the inter-batwing angles between two classes, with one **(A)** and both **(B)** rendered as wireframes. The maps were aligned on the right half of the complex.

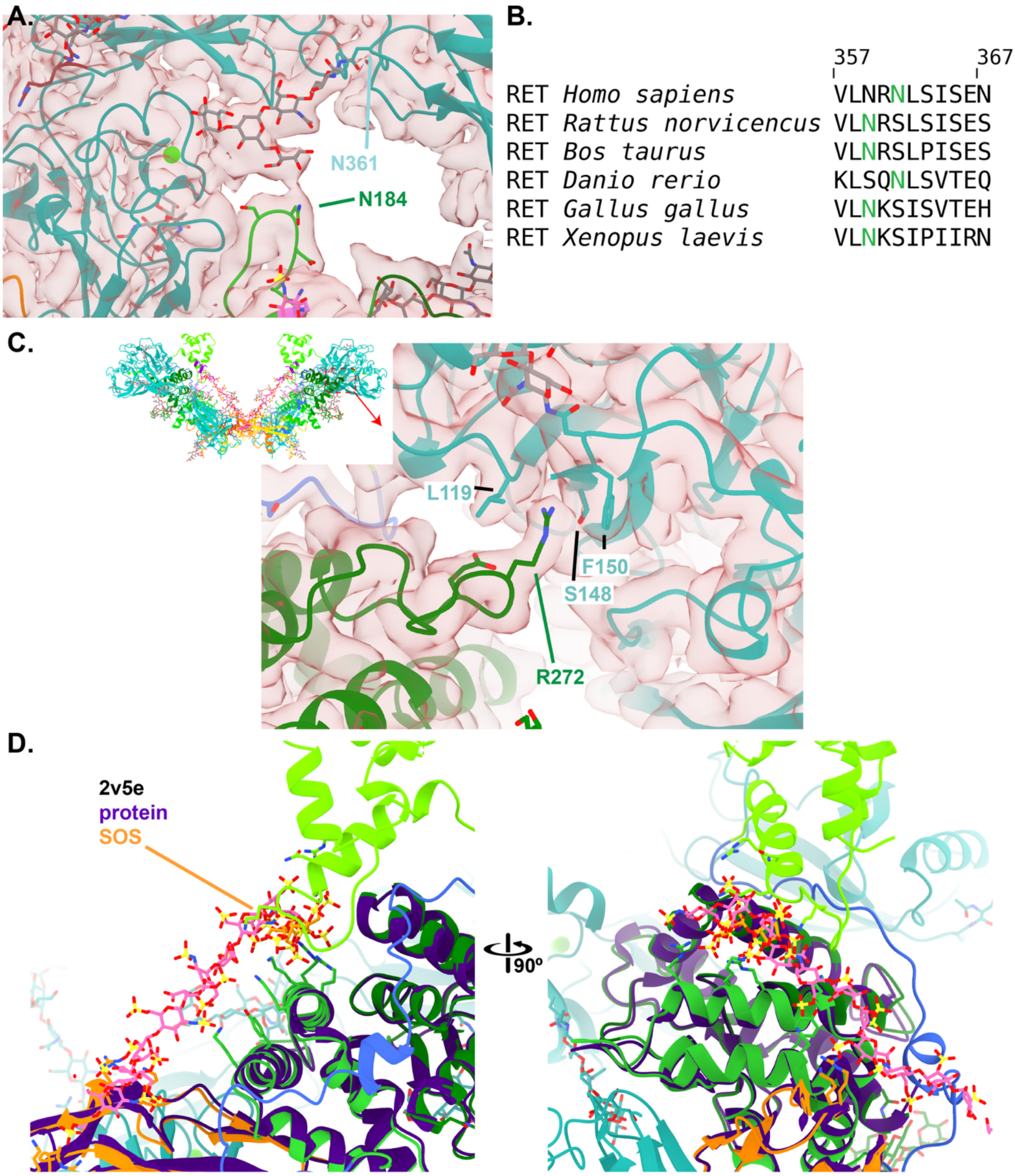


**Figure S12: (A)** Detail of the putative interaction of the hRET$^{ECD}$ N-glycan attached to N361 and the sidechain of hGFRα1 residue N184. **(B)** Conservation of the N-glycan sequon at RET N361 across vertebrate clades. **(C)** Detail of the interaction of the sidechain hGFRα1 R272 and a pocket on RET CLD1. **(D)** Comparison of the binding modes of heparin to the hGFRα1 GBS in the hRGα1-HP20 structure and SOS to a ∆D1 construct of GFRα1 (PDB: 2V5E).

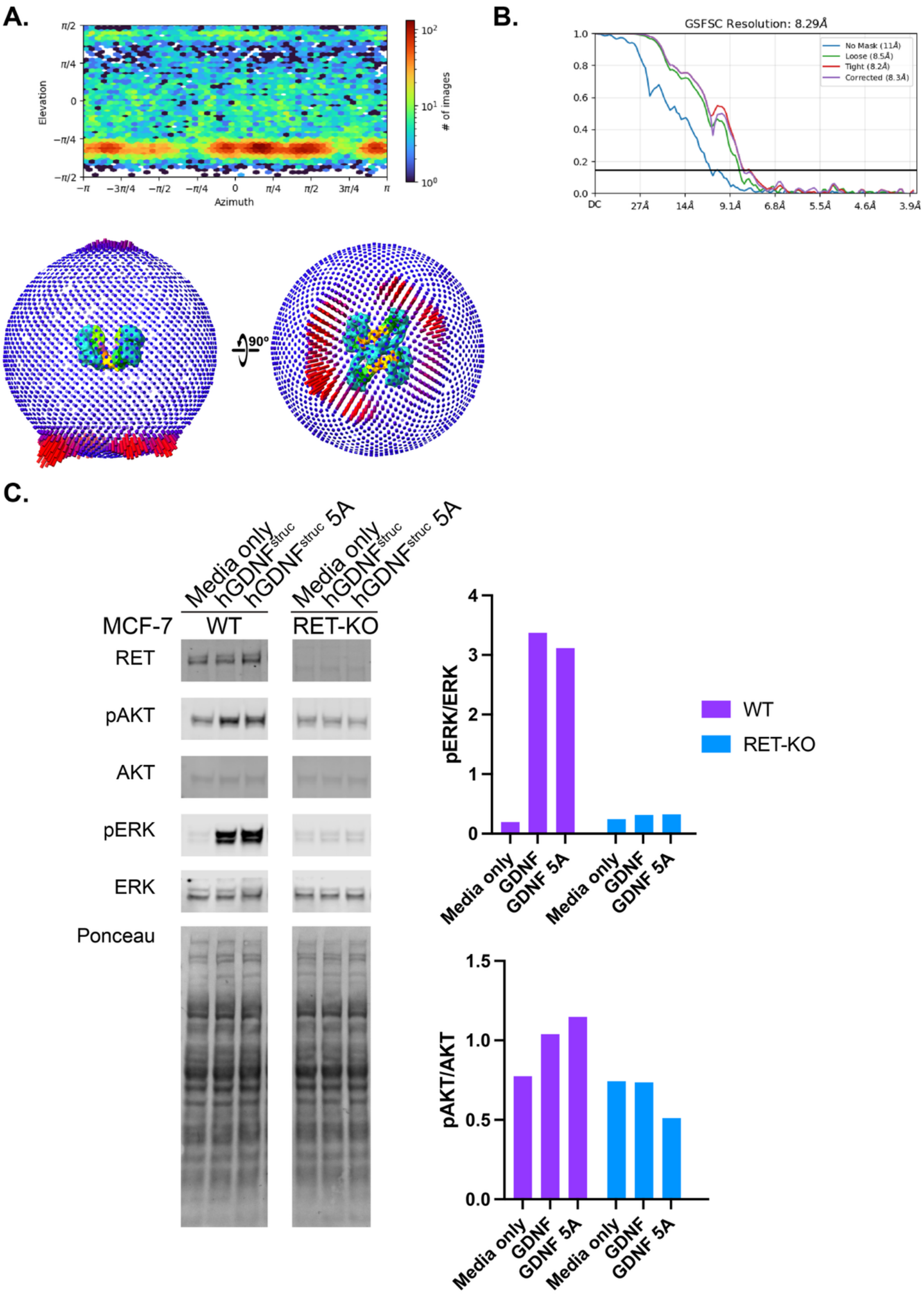


**Figure S13:** **(A)** Orientation distribution of particles which reconstruct to the final hRGα1-HP20 tetrameric form map, represented as a heatmap and in 3D space. **(B)** Gold-standard Fourier shell correlation (GSFSC) plot for the final hRGα1-HP20 tetrameric map with C2 imposed. **(C)** Stimulation of Akt and ERK phosphorylation in WT and RET KO MCF7 cells with hGDNF$^{struc}$.

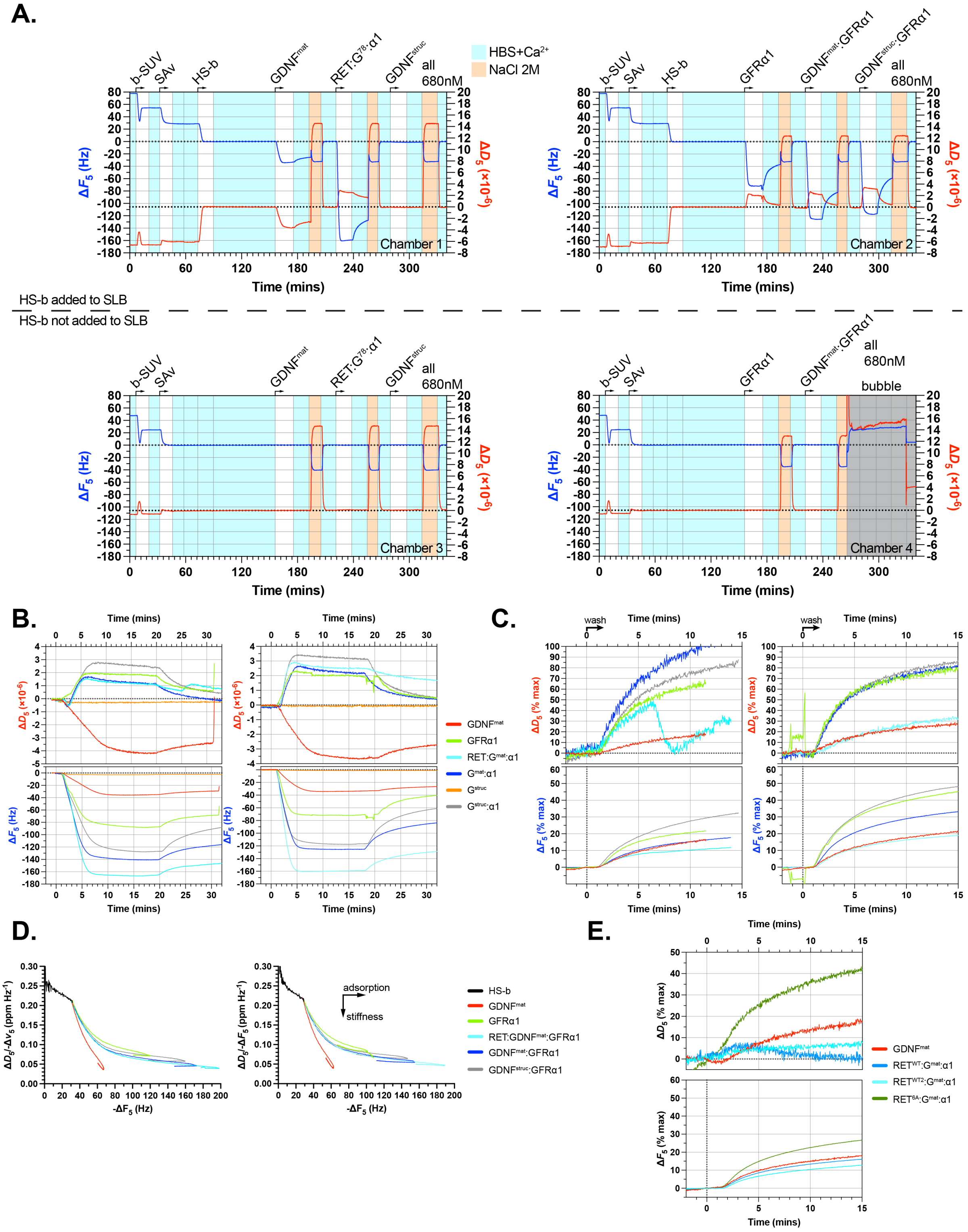


**Figure S14: (A)** QCM-D traces (Δ*F* and Δ*D*, both *i*=5) for all four parallel flow chambers detailing model cell surface assembly steps (SLB formation from SUVs, SAv monolayer and HS-b ‘brush’ formation), analyte binding and elution steps, and model cell surface regeneration with high salt buffer. Model cell surfaces were consistently of good quality, with the expected QCM-D responses established previously for SLB and SAv monolayer formation. The large increase in Δ*D* on HS-b incubation (by ~6 $\times$ $10^{-6}$ at saturation) alongside the decrease in Δ*F* (by -25 Hz at saturation) indicates a soft film is being formed, as expected for a brush of HS polysaccharides[3]. Chambers 1 & 3 featured the HS-b brush, but chambers 2 & 4 did not for us as negative binding controls. The model cell surfaces were fully stable throughout the experiments, as demonstrated by the return of QCM-D responses to Δ*F* = Δ*D* = 0 (as offset in the graphs) after each regeneration step. Note that the transient large decreases in Δ*F* and increases in Δ*D* in the regeneration step are mainly caused by increases in the viscosity and density of the solution in the presence of high salt and thus should not be interpreted as surface effects. Virtually no binding to plain SAv-on-SLB surfaces (chambers 2 & 4) was observed for any of the analytes, demonstrating good specificity of observed interactions. **(B)** QCM-D data for analyte adsorption and desorption assays aligned in time. Left and right plots are two independent replicates. The decrease in Δ*D* observed here for $hGDNF^{mat}$ indicates a substantive stiffening of the HS film. **(C)** QCM-D data for the analyte desorption aligned in time with Δ*F* and Δ*D* offset and normalised such that 0% corresponds to the start of the elution phase and 100% to full protein desorption (i.e., the signal for the plain HS-b film). Left and right plots are two independent replicates. **(D)** Parametric plots of Δ*D*/-Δ*F* (a measure of the softness of the HS film, alone or with bound analyte) vs. -Δ*F* (a measure of HS surface coverage) for each analyte[4,5], derived from the QCM-D data shown in A-B. Note that in these plots Δ*F* and Δ*D* were offset to zero at the point before the SAv monolayer was functionalised with HS-b, such that responses represent the HS film on its own (grey part of the data traces) or with bound analyte (coloured parts of the data traces; the turns at the end of these traces correspond to the analyte desorption phases). The Δ*D*/-Δ*F* traces for $hGDNF^{mat}$ are consistently lower than for any of the other analytes, demonstrating that this protein on its own has a particularly high propensity to stiffen the HS film which we propose is due to protein-mediated HS cross-linking not present in any complexes of hGDNF. Left and right plots represent independent replicates. **(E)** Plot (presented as C) illustrating the desorption kinetics including $hRET^{ECD}$-6A (R417A/R418A/K471A/R474A/R475A/K477A) targeting the CLD4 HS binding site.

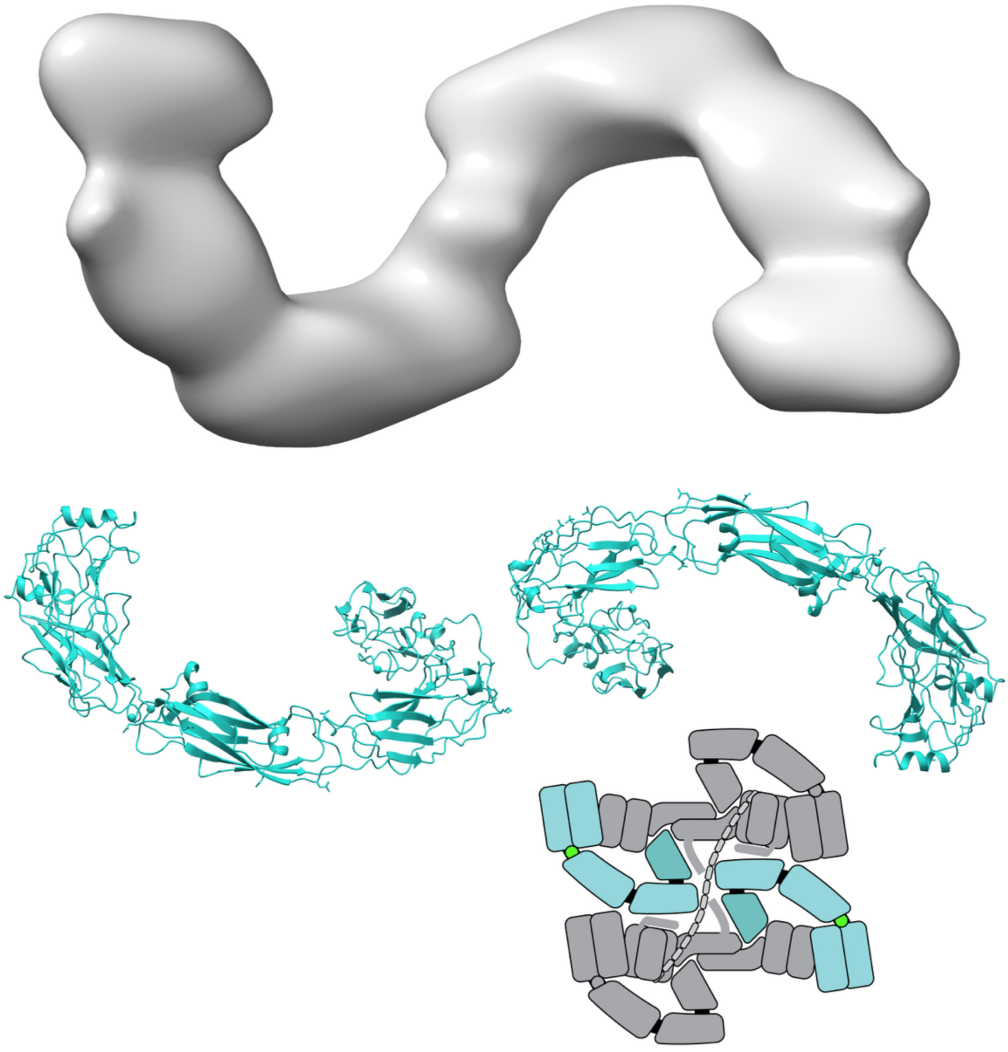

**Figure S15:** Comparison of the aberrant dimer formed by $RET^{ECD}$ with the MEN2A mutation C634 with the arrangement of RETs formed in the hRGα1-HP20 tetrameric structure. A schematic of the hRGα1-HP20 tetrameric structure is shown with the hidden chains in grey.

#### Supplementary Tables

| **Protein** | **Analyte (variable conc.)** | $K_d$/nM | $\Delta F^{MST}_{norm}$/‰ | **colour** |
|---|---|---|---|---|
| GFRα1-CHis | SOS | 47±6 | 21 | green |
| GFRα1-CHis | Heparin DP4 | 1700±300 | 22 | gold |
| GFRα1-CHis | Heparin DP6 | 180±20 | 25 | orange |
| GFRα1-CHis | HS-HepIII DP10 | 27±4 | 27 | red |
| GFRα1-CHis | Heparin DP20 | 16±4 | 25 | dark red |
| GFRα1-CHis | CS-A/C DP10 | 12000±2000 | 24 | turquoise |
| GFRα1-CHis | CS-D DP10 | 12000±1000 | 22 | dark olivegreen |
| GFRα1-CHis | DP6-91 | 29±6 | 22 | magenta |
| GFRα1-CHis | fondaparinux | 90±10 | 24 | midnight blue |
| GFRα1-CHis D2-AAA | SOS | 40000±10000 | 9 | dark cyan |
| GFRα1-CHis D2-AAA | fondaparinux | 50000±20000 | 9 | coral |

**Table S1. Analysis of GFRα1-GAG interactions by MST.** $K_d$ values are given ±the standard error for the least squares fit of the 1:1 binding model to the data as calculated in the MO.Affinity application. $\Delta F^{MST}_{norm}$ is calculated from the fitted binding model.

| **Protein (variable concentration)** | **Protein (4μM constant)** | **Ligand (constant 80 μM)** | $K_d$/nM | **Δ**$F_{norm}$/‰ | **colour** |
|---|---|---|---|---|---|
| hGFRα1-CHis | - | - | 1400±500 | 18 | magenta |
| hGFRα1-CHis | $hGDNF^{struc}$ | - | 19±6 | 28 | orange |
| hGFRα1-CHis | $hGDNF^{struc}$ | SOS | 16±5 | 45 | green |
| hGFRα1-CHis | $hGDNF^{struc}$ | Heparin DP20 | 9±3 | 45 | crimson |

**Table S2. Analysis of $hRET^{ECD*}$ interactions by MST.** $K_d$ values are given ±the standard error for the least squares fit of the 1:1 binding model to the data as calculated in the MO.Affinity application. $\Delta F^{MST}_{norm}$ is calculated from the fitted binding model.

| Position | Probe | hGFRa1 WT | hGFRa1 D2-AAA |
|---|---|---|---|
| 1 | HA-S4* | - | - |
| 2 | HA-S6* | - | - |
| 3 | HA-S8* | - | - |
| 4 | HA-S10* | - | - |
| 5 | HA-S12* | - | - |
| 6 | HA-S14* | - | - |
| 7 | HA-S16* | - | - |
| 8 | HA-S18* | - | - |
| 9 | CSA-2-AO* (Nian) | 2289 | - |
| 10 | CSA-4-AO* (Nian) | 1341 | - |
| 11 | CSA-6-AO* (Nian) | - | - |
| 12 | CSA-8-AO* (Nian) | 1765 | - |
| 13 | CSA-10-AO* (Nian) | - | - |
| 14 | CSA-12-AO* (Nian) | - | - |
| 15 | CSA-14-AO* (Nian) | - | - |
| 16 | CSA-16-AO* (Nian) | - | - |
| 17 | CSA-18-AO* (Nian) | - | - |
| 18 | CSA-20-AO* (Nian) | 2334 | - |
| 19 | CSB-2-AO* (Nian) | 7604 | - |
| 20 | CSB-4-AO* (Nian) | 5746 | - |
| 21 | CSB-6-AO* (Nian) | 7750 | - |
| 22 | CSB-8-AO* (Nian) | 7674 | - |
| 23 | CSB-10-AO* (Nian) | 11961 | - |
| 24 | CSB-12-AO* (Nian) | 6973 | - |
| 25 | CSB-14-AO* (Nian) | 4681 | - |
| 26 | CSB-16-AO* (Nian) | 8048 | - |
| 27 | CSB-18-AO* (Nian) | 9959 | - |
| 28 | CSB-20-AO* (Nian) | 7487 | - |
| 29 | CSC-2-AO* (Nian) | 3096 | - |
| 30 | CSC-4-AO* (Nian) | 1168 | - |
| 31 | CSC-6-AO* (Nian) | 1605 | - |
| 32 | CSC-8-AO* (Nian) | 3274 | - |
| 33 | CSC-10-AO* (Nian) | 1520 | - |
| 34 | CSC-12-AO* (Nian) | 1971 | - |
| 35 | CSC-14-AO* (Nian) | 6092 | - |
| 36 | CSC-16-AO* (Nian) | 6681 | - |
| 37 | CSC-18-AO* (Nian) | 6509 | - |
| 38 | CSC-20-AO* (Nian) | 6050 | - |
| 39 | Hep-2-AO*(Nian) | 6251 | - |

| | | | |
|---|---|---|---|
| 40 | Hep-4-AO*(Nian) | 9264 | 1869 |
| 41 | Hep-6-AO*(Nian) | 13262 | 1370 |
| 42 | Hep-8-AO*(Nian) | 9337 | 4205 |
| 43 | Hep-10-AO*(Nian) | 12110 | 3866 |
| 44 | Hep-12-AO*(Nian) | 15475 | 7060 |
| 45 | Hep-14-AO*(Nian) | 15126 | 2550 |
| 46 | Hep-16-AO*(Nian) | 18020 | 2796 |
| 47 | Hep-18-AO*(Nian) | 19159 | 4779 |
| 48 | Hep-20-AO*(Nian) | 16695 | 4946 |
| 49 | HS-S4-AO* | 7405 | - |
| 50 | HS-S6-AO* | 14499 | 257 |
| 51 | HS-S8-AO* | 13782 | 2855 |
| 52 | HS-S4a-AO | 14260 | - |
| 53 | HS-S6a-AO | 19628 | 2402 |
| 54 | HS-S8a-AO | 9874 | - |
| 55 | Hep-S4-AO* | 12672 | 4484 |
| 56 | Hep-S6-AO* | 18597 | 1540 |
| 57 | Hep-S8-AO* | 16440 | 4549 |
| 58 | Hep-S8a-AO | 12307 | 2385 |
| 59 | KS-2-AO*[K'ase I, 23-27h] | 638 | - |
| 60 | KS-4-AO*[K'ase I, 23-27h] | 938 | - |
| 61 | KS-8-AO*[K'ase I, 23-27h] | 3230 | - |
| 62 | KS-6-AO*[K'ase I, 23-27h] | 1004 | - |
| 63 | KS-10-AO*[K'ase I, 23-27h] | - | - |
| 64 | KS-12-AO*[K'ase I, 23-27h] | 1377 | - |
| 65 | KS-14-AO*[K'ase I, 23-27h] | - | - |
| 66 | KS-16-AO*[K'ase I, 23-27h] | 1073 | - |
| 67 | KS-18-AO*[K'ase I, 23-27h] | 2135 | - |
| 68 | KS-20-AO*[K'ase I, 23-27h] | 3624 | - |
| 69 | KS-2-AO*[K'aseII,7.5h] | 2946 | - |
| 70 | KS-4-AO*[K'aseII,7.5h] | 425 | - |
| 71 | KS-6-AO*[K'aseII,7.5h] | - | - |
| 72 | KS-8-AO*[K'aseII,7.5h] | - | - |
| 73 | KS-10-AO* [K'aseII,7.5h] | - | - |
| 74 | KS-12-AO*[K'aseII,7.5h] | 335 | - |
| 75 | KS-14-AO*[K'aseII,7.5h] | 23 | - |
| 76 | KS-16-AO*[K'aseII,7.5h] | 1632 | - |
| 77 | KS-18-AO*[K'aseII,7.5h] | - | - |
| 91 | Lac-AO | - | - |
| 92 | SU-Tyr | 4496 | - |

**Table S3.** Binding of hGFRα1-CHis WT/D2-AAA to an NGL microarray. Shows fluorescence intensity of detected binding to each probe. Cells are colour-coded to value relative to the highest in the experiment: red (>70%), orange (30-70%), yellow (10-30%), blue (<10%). Lac-AO is a lactose negative control and SU-Tyr is a sulfated tyrosine control.

| | WT | | D2-AAA | | Probe | | | |
|---|---|---|---|---|---|---|---|---|
| **HS#** | **Mean** | **SD** | **Mean** | **SD** | **DP** | **# Sulf** | **Ido 2S** | **3S** |
| **3** | 4885 | 183 | 597 | 126.8 | 4 | 5 | 1 | |
| **4** | 3314 | 152 | 524 | 60.22 | 4 | 4 | 0 | |
| **5** | 1143 | 109 | 301.5 | 35.02 | 4 | 3 | 1 | |
| **7** | 2147 | 208 | 426.5 | 72.43 | 4 | 2 | 0 | |
| **8** | 3054 | 81 | 467.3 | 13.44 | 4 | 4 | 0 | |
| **9** | 261 | 162 | 79.5 | 48.38 | 4 | 2 | 0 | |
| **10** | 90 | 62 | 212.5 | 24.09 | 4 | 2 | 0 | |
| **11** | 304 | 280 | 114 | 49.03 | 4 | 3 | 1 | |
| **12** | 1228 | 331 | 190.8 | 115 | 4 | 4 | 0 | |
| **13** | 1471 | 543 | 166.8 | 64.64 | 4 | 4 | 0 | |
| **14** | 274 | 83 | 200.8 | 74.54 | 4 | 2 | 0 | |
| **15** | 52 | 57 | 105.8 | 45.07 | 4 | 1 | 1 | |
| **16** | 582 | 186 | 144 | 102.1 | 4 | 3 | 1 | |
| **17** | 400 | 22 | 118.5 | 74.3 | 4 | 2 | 1 | |
| **18** | 2145 | 87 | 362.5 | 16.87 | 4 | 4 | 1 | |
| **19** | 160 | 111 | 154.8 | 38.09 | 4 | 1 | 1 | |
| **20** | 741 | 103 | 227.5 | 75.74 | 4 | 3 | 1 | |
| **21** | 1 | 0 | 3 | 3.46 | 4 | 0 | 0 | |
| **22** | 235 | 45 | 115.8 | 34.52 | 4 | 2 | 0 | |
| **23** | 3541 | 548 | 563.8 | 139.6 | 4 | 4 | 1 | |
| **24** | 504 | 75 | 196.3 | 51.67 | 4 | 2 | 1 | |
| **25** | 900 | 154 | 266.8 | 51.02 | 4 | 3 | 1 | |
| **26** | 36 | 43 | 81.25 | 49.42 | 4 | 1 | 0 | |
| **27** | 407 | 200 | 166.5 | 11.67 | 4 | 3 | 0 | |
| **28** | 45 | 44 | 20.5 | 24.94 | 4 | 0 | 0 | |
| **29** | 1 | 0 | 30 | 50.23 | 2 | 1 | 0 | |
| **30** | 69 | 51 | 61.75 | 60.18 | 4 | 1 | 0 | |
| **31** | 8769 | 1055 | 839.8 | 14.92 | 4 | 5 | 1 | |
| **32** | 886 | 72 | 211.5 | 48.53 | 4 | 3 | 0 | |
| **33** | 1816 | 145 | 343.3 | 40.39 | 4 | 3 | 1 | |
| **34** | 412 | 84 | 103.8 | 63.65 | 4 | 2 | 1 | |
| **35** | 46 | 67 | 17.5 | 17.04 | 4 | 1 | 1 | |
| **36** | 3373 | 351 | 225 | 224.5 | 4 | 4 | 1 | |

| | | | | | | | | |
|---|---|---|---|---|---|---|---|---|
| **37** | 747 | 222 | 112.5 | 87.1 | 4 | 3 | 1 | |
| **38** | 9448 | 655 | 505.3 | 300.3 | 4 | 5 | 1 | |
| **39** | 3634 | 518 | 364.8 | 74.61 | 4 | 4 | 2 | |
| **40** | 14547 | 1595 | 1374 | 132.1 | 4 | 6 | 2 | |
| **41** | 323 | 191 | 123 | 91.51 | 4 | 2 | 1 | |
| **42** | 4959 | 1071 | 335.8 | 99.13 | 4 | 5 | 1 | |
| **43** | 796 | 179 | 157 | 83.14 | 2 | 2 | 1 | |
| **44** | 1217 | 415 | 208 | 105.2 | 2 | 3 | 1 | |
| **45** | 57 | 23 | 54 | 44.34 | 2 | 1 | 0 | |
| **46-D** | 50 | 47 | 128.8 | 106.1 | 4 | 2 | 0 | |
| **47** | 99 | 97 | 42.25 | 41.29 | 4 | 1 | 0 | |
| **48** | 988 | 49 | 444.8 | 77.15 | 4 | 1 | 0 | |
| **49** | 45790 | 1060 | 3873 | 303.1 | 4 | 6 | 1 | yes |
| **52** | 9443 | 760 | 742.8 | 63.81 | 4 | 5 | 1 | yes |
| **53** | 1 | 0 | 19.5 | 24.63 | 4 | 1 | 0 | |
| **54-D** | 16063 | 416 | 648.3 | 109.6 | 8 | 7 | 1 | |
| **55** | 24 | 28 | 89 | 56.44 | 4 | 2 | 0 | |
| **56** | 37 | 61 | 84.25 | 74.93 | 4 | 2 | 0 | |
| **57** | 3796 | 177 | 288.3 | 73.19 | 4 | 4 | 1 | |
| **58-D** | 124 | 160 | 49.25 | 48.09 | 4 | 3 | 0 | |
| **59** | 1 | 0 | 23.25 | 38.54 | 4 | 4 | 1 | |
| **60-D** | 4 | 5 | 1 | 0 | 4 | 3 | 0 | |
| **62** | 21294 | 11463 | 13.25 | 21.22 | 6 | 7 | 0 | yes |
| **63** | 1153 | 360 | 1505 | 197 | 6 | 6 | 0 | |
| **64** | 104227 | 8127 | 219.3 | 51.45 | 6 | 7 | 0 | yes |
| **65** | 11125 | 388 | 4811 | 135.5 | 6 | 6 | 0 | |
| **66** | 194748 | 9955 | 770 | 124.2 | 6 | 8 | 1 | yes |
| **67** | 65019 | 2228 | 11547 | 423.2 | 6 | 7 | 1 | |
| **68** | 1658 | 20 | 2999 | 205.8 | 6 | 7 | 0 | yes |
| **69** | 2049 | 290 | 15.25 | 14.46 | 6 | 6 | 0 | |
| **70** | 93766 | 7202 | 191.5 | 14.15 | 6 | 7 | 0 | yes |
| **71** | 7339 | 446 | 2478 | 97.11 | 6 | 6 | 0 | |
| **72** | 73528 | 6486 | 590.3 | 25.62 | 6 | 8 | 1 | yes |
| **73** | 110320 | 4109 | 2297 | 191.3 | 6 | 7 | 1 | |
| **74** | 119451 | 6332 | 6364 | 209.3 | 6 | 8 | 1 | yes |
| **75** | 14294 | 579 | 7449 | 176.6 | 6 | 7 | 1 | |
| **76** | 113561 | 799 | 1400 | 57.48 | 6 | 8 | 1 | yes |
| **77** | 46350 | 2564 | 7417 | 325.1 | 6 | 7 | 1 | |
| **78** | 417639 | 13998 | 1884 | 228.2 | 6 | 9 | 2 | yes |
| **79** | 312102 | 15780 | 22046 | 674.8 | 6 | 8 | 2 | |

| | | | | | | | | |
|---|---|---|---|---|---|---|---|---|
| **80** | 15614 | 1139 | 12508 | 504.5 | 6 | 6 | 0 | |
| **81** | 9293 | 257 | 952.3 | 39.88 | 6 | 6 | 0 | yes |
| **82** | 66398 | 1930 | 452.3 | 65.97 | 6 | 7 | 1 | yes |
| **83** | 23343 | 1688 | 4196 | 414.1 | 6 | 6 | 0 | yes |
| **84** | 25972 | 1191 | 1136 | 179 | 6 | 6 | 0 | yes |
| **85** | 55258 | 2870 | 1064 | 101.8 | 6 | 7 | 1 | yes |
| **86** | 54079 | 1391 | 2461 | 155.9 | 6 | 7 | 1 | yes |
| **87** | 111940 | 5985 | 2636 | 125.1 | 6 | 7 | 1 | yes |
| **88** | 121490 | 4411 | 4562 | 303.4 | 6 | 8 | 2 | yes |
| **89** | 1312 | 40 | 6007 | 336.7 | 2 | 3 | 1 | |
| **90** | 21013 | 391 | 138.3 | 23.94 | 4 | 6 | 2 | |
| **91** | 514067 | 21598 | 1748 | 85.17 | 6 | 9 | 3 | |
| **92** | 581062 | 20288 | 38595 | 1243 | 8 | 12 | 4 | |
| **93** | 4712 | 442 | 59625 | 2157 | 6 | 5 | 2 | |
| **94** | 253744 | 9616 | 655.3 | 74.55 | 6 | 7 | 2 | |
| **95** | 10101 | 456 | 5936 | 497.8 | 6 | 6 | 2 | |
| **96** | 24969 | 981 | 1081 | 69.2 | 6 | 6 | 2 | |
| **97** | 11307 | 986 | 2416 | 289.9 | 4 | 5 | 2 | |
| **98** | 1855 | 204 | 2128 | 215.8 | 4 | 5 | 1 | |
| **99** | 825 | 71 | 318.8 | 56.94 | 4 | 3 | 1 | |
| **100** | 89 | 25 | 261 | 27.12 | 4 | 1 | 0 | |
| **101** | 42 | 29 | 77 | 83.32 | 4 | 1 | 0 | |
| **102** | 119173 | 6415 | 120.5 | 77.4 | ~40 | | | |

**Table S4.** HS sequence-defined array data. Fluorescence intensities and standard deviation is listed for the 3 µg/mL assay for hGFRα1-CHis WT and D2-AAA. Mean intensities are coloured indicating binding relative to the highest binding probe (blue: low, red: high).

| **Antibody** | **Clone** | **Supplier** | **Catalogue no.** | **Dilution** | **Host** |
|---|---|---|---|---|---|
| ERK | Clone-16 (pan ERK) | BD bioscience | 610124 | 1:2500 | Mouse |
| pERK (T202/Y204) | D13.14.4E | Cell Signalling Technologies (CST) | 4370 | 1:2000 | Rabbit |
| AKT | 40D4 | CST | 2920 | 1:2000 | Mouse |
| AKT-biotinylated | 40D4 | CST | 4821 | 1:2000 | Mouse-biotinylated |
| pAKT (ser473) | D9E | CST | 4060 | 1:2000 | Rabbit |
| Anti-Mouse | | LiCor | 926-68070 | 1:20,000 | Goat |
| Anti-Rabbit | | LiCor | 926-32211 | 1:20,000 | Goat |
| Streptavidin | | LiCor | 925-68079 | 1:5000 | N/A |

**Table S5.** Antibodies used for Western blotting.

| Position | Probe | hGDNF$^{mat}$ | hGDNF$^{mat}$-9A |
|---|---|---|---|
| 1 | HA-S6 | - | 51 |
| 2 | HA-S10 | - | - |
| 3 | HA-S14 | - | - |
| 4 | HA-S18 | - | - |
| 5 | CSA-6-AO | 166 | 42 |
| 6 | CSA-10-AO | 68 | - |
| 7 | CSA-14-AO | 499 | - |
| 8 | CSA-18-AO | 933 | - |
| 9 | CSB-6-AO | 2767 | 150 |
| 10 | CSB-10-AO | 5453 | - |
| 11 | CSB-14-AO | 1981 | - |
| 12 | CSB-18-AO | 14143 | - |
| 13 | CSC-6-AO | 2364 | 186 |
| 14 | CSC-10-AO | 248 | - |
| 15 | CSC-14-AO | 3786 | - |
| 16 | CSC-18-AO | 4376 | - |
| 17 | CSD-4-AO | 791 | - |
| 18 | CSD-6-AO | 827 | - |
| 19 | CSD-8-AO | 79 | 118 |
| 20 | CSD-10-AO | 410 | - |
| 21 | CSD-12-AO | 2247 | 117 |
| 22 | CSD-14-AO | 2683 | - |
| 23 | Hep-4-AO (new) | 3856 | 147 |
| 24 | Hep-6-AO (ex new) | 7374 | 142 |
| 25 | Hep-6-AO (new) | 5168 | - |
| 26 | Hep-8-AO (new) | 8053 | 100 |
| 27 | Hep-10-AO (ex new) | 10004 | - |
| 28 | Hep-10-AO(new) | 9086 | - |
| 29 | Hep-12-AO (new) | 10855 | 227 |
| 30 | Hep-14-AO (ex new) | 12239 | - |
| 31 | Hep-18-AO (ex new) | 22807 | - |
| 32 | HS-S6-AO | 8743 | 111 |
| 33 | HS-S8-AO | 11217 | - |
| 34 | KS-6-AO | - | - |
| 35 | KS-10-AO | - | - |
| 36 | KS-14-AO | - | 137 |
| 37 | KS-18-AO | 2365 | - |
| 38 | 2-O-DeS Hep-8-AO | 6611 | - |
| 39 | 2-O-DeS Hep-10-AO | 7547 | 88 |
| 40 | 6-O-DeS Hep-8-AO | 4787 | - |

| | | | |
|---|---|---|---|
| 41 | 6-O-DeS Hep-10-AO | **5159** | **166** |
| 42 | N-DeS Hep-8-AO | **5586** | **-** |
| 43 | N-DeS Hep-10-AO | **7390** | **127** |
| 44 | N-DeS ReAc Hep-8-AO | **1805** | **-** |
| 45 | N-DeS ReAc Hep-10-AO | **2465** | **171** |
| 46 | N-DeS ReAc Hep-12-AO | **9290** | **30** |

**Table S7.** Binding of hGDNF$^{mat}$ WT/9A to an NGL GAG microarray. Shows fluorescence intensity of detected binding to each probe. Cells for WT are colour-coded to value relative to the highest in the experiment: red (>70%), orange (30-70%), yellow (10-30%), blue (<10%).

#### References


1. Gouet, P., Courcelle, E., Stuart, D.I., and M√©toz, F. (1999). ESPript: analysis of multiple sequence alignments in PostScript. Bioinformatics *15*, 305–308. https://doi.org/10.1093/bioinformatics/15.4.305.

2. Liebschner, D., Afonine, P.V., Baker, M.L., Bunkóczi, G., Chen, V.B., Croll, T.I., Hintze, B., Hung, L.-W., Jain, S., McCoy, A.J., et al. (2019). Macromolecular structure determination using X-rays, neutrons and electrons: recent developments in *Phenix*. Acta Crystallogr D Struct Biol *75*, 861–877. https://doi.org/10.1107/S2059798319011471.

3. Srimasorn, S., Souter, L., Green, D.E., Djerbal, L., Goodenough, A., Duncan, J.A., Roberts, A.R.E., Zhang, X., Débarre, D., DeAngelis, P.L., et al. (2022). A quartz crystal microbalance method to quantify the size of hyaluronan and other glycosaminoglycans on surfaces. Sci Rep *12*, 10980. https://doi.org/10.1038/s41598-022-14948-7.

4. Migliorini, E., Thakar, D., Sadir, R., Pleiner, T., Baleux, F., Lortat-Jacob, H., Coche-Guerente, L., and Richter, R.P. (2014). Well-defined biomimetic surfaces to characterize glycosaminoglycan-mediated interactions on the molecular, supramolecular and cellular levels. Biomaterials *35*, 8903–8915. https://doi.org/10.1016/j.biomaterials.2014.07.017.

5. Reviakine, I., Johannsmann, D., and Richter, R.P. (2011). Hearing What You Cannot See and Visualizing What You Hear: Interpreting Quartz Crystal Microbalance Data from Solvated Interfaces. Anal. Chem. *83*, 8838–8848. https://doi.org/10.1021/ac201778h.
